# Modulation of the speed of Intrinsic Alpha Frequency (IAF) with tACS modulates variability of illusory speech perception

**DOI:** 10.64898/2026.10.07.756537

**Authors:** Shreya Deb, Debottam Bhunia, Vinsea A V Singh, Vignesh Muralidharan, Dipanjan Roy

## Abstract

Navigating the world requires our brains to constantly process a massive influx of information from the environment. One of the most remarkable yet complex feats the human brain performs daily is speech perception. It is a multifaceted, hierarchical process that relies heavily on the integration of auditory and visual cues, particularly in noisy environments. Neural integration of these streams is hypothesized to depend on temporal mechanisms governed by individual alpha frequency (IAF, 8-12 Hz), which serves as a key neural correlate of the temporal binding window (TBW). While IAF has been linked to multisensory integration, its role in shaping multisensory illusions remains largely unexplored. Here, we tested this directly using transcranial alternating current stimulation (tACS) to modulate occipital activity during multisensory conflict. In a two-day protocol involving 27 healthy participants, resting-state EEG was recorded to determine IAF. On Day 2, participants then performed a multisensory speech task involving congruent (pa/, /ta/, /ka/) and incongruent McGurk stimuli (auditory /pa/, visual /ka/), both at baseline and during tACS applied at IAF and offset frequencies (IAF ± 2 Hz) over occipital electrodes. While accuracy for congruent syllables remained high across conditions, results revealed that tACS significantly reduced the perception of illusory /ta/ percepts during McGurk trials. Finally, using a phenomenological model based on Kuramoto oscillators, we provided a mechanistic explanation for the percentage of illusory responses observed across participants under the influence of tACS at IAF and offsets. This study highlights the potential for targeted neuromodulatory interventions to influence multisensory processing in clinical or professional settings.

## 1. Introduction

Speech perception is a hierarchical process that integrates multiple sensory modalities (O’Sullivan et al., 2021), particularly when comprehension is challenged by background noise or competing speakers. Segmentation and analysis of phonemes, syllables, and words depend on both auditory and visual cues (Mitterer & Reinisch, 2017), such as the rhythmic movement of a speaker’s lips, which fundamentally shapes auditory perception. A quintessential demonstration of this cross-modal integration is the McGurk effect (Mcgurk & Macdonald, 1976), an illusion in which incongruent audiovisual (AV) stimuli (e.g., auditory /pa/ paired with visual /ka/) yield a fused percept such as /ta/. Its magnitude and occurrence are modulated by both bottom-up sensory factors, such as auditory noise or reduced visual salience, which systematically scale illusory percept (Munhall et al., 2004; Stacey et al., 2020), and top-down factors including spatial attention, perceptual expectation, and task instructions (Alsius et al., 2005; Getz & Toscano, 2021; Tiippana et al., 2004; Windmann, 2004).

Neuroimaging and electrophysiological studies have mapped the networks subserving this integration. The superior temporal sulcus (STS) and posterior STS (pSTS) are consistently identified as primary hubs for multisensory speech integration (for review, see Beauchamp 2015), though AV integration extends beyond the STS to require dynamic coordination among primary auditory and visual cortices (Paulesu et al., 2003) and frontoparietal networks (Kumar et al., 2016, 2020; Shyamchand Singh et al., 2024). Using fMRI, Benoit and colleagues showed that BOLD signal in primary auditory and visual cortices not just the STS tracked perception of the McGurk fusion, implying a distributed network for multisensory integration, (Benoit et al., 2010). Consistent with this, AV temporal correspondence modulates activity jointly in the multisensory STS and primary sensory cortices, indicating that binding is expressed at early sensory stages as well as associative cortex (Noesselt et al., 2007). Furthermore, the motor theory of speech perception proposes that listening to and observing speech modulates the excitability of sensorimotor circuits involved in speech production (GALANTUCCI et al., 2006; Liessens & Ladouce, 2024). Given that AV integration recruits such spatially distributed regions, from primary sensory cortices to the pSTS and sensorimotor areas, a central question emerges: how does the brain coordinate activity across these regions with sufficient temporal precision to bind auditory and visual signals into a unified percept? Neural oscillations, particularly the alpha rhythm, have increasingly been proposed as the answer (Cecere et al., 2015; Keil & Senkowski, 2018; Samaha & Postle, 2015).

## 1.1 Neural Oscillations and the Role of Alpha

Spatially distributed neuronal populations achieve the necessary synchrony for binding through neural oscillations (Senkowski et al., 2008; Senkowski & Engel, 2024). Van Rullen proposed that perception is parsed into discrete ‘perceptual cycles’ organised by endogenous brain rhythms rather than flowing continuously (VanRullen, 2016; VanRullen & Koch, 2003). Moreover, the alpha band (8-12 Hz) acts as a sensory gating mechanism and internal pacemaker, with the Individual Alpha Frequency (IAF) each person’s peak alpha frequency dictating the temporal resolution of visual and multisensory perception (Cecere et al., 2015; Venskus & Hughes, 2021).

In multisensory speech perception, the brain faces a ‘causal inference’ problem: determining whether auditory and visual signals arise from a common source before integrating them (Chen & Spence, 2017; Körding et al., 2007; Welch & Warren, 1980). This is governed partly by the Temporal Window of Integration (TWI), the interval within which two stimuli are judged to originate from the same event. Sensory inputs falling within the same alpha cycle are proposed to be perceptually bound, while those in separate cycles are segregated (Cecere et al., 2015) suggesting that an individual’s alpha speed correlates with their TWI width and, in turn, their susceptibility to temporal illusions.

Moreover, prestimulus brain states also biases AV integration, offering correlational evidence for alpha’s functional relevance beyond IAF alone. Prestimulus alpha and beta power and phase strongly predict trial-by-trial McGurk responses (Keil et al., 2012; Roa Romero et al., 2015). Notably, this extends to aperiodic activity: a flatter 1/f slope and lower aperiodic offset in temporo-occipital sensors prior to stimulus onset predict McGurk perception (Singh et al., 2025), directly implicating occipital regions in illusion gating. Moreover, convergent evidence from computational neural mass models similarly assigns a central role to visual processing nodes, showing that the comparatively slow dynamics of occipital/visual nodes, interacting with faster auditory nodes and intermediate hubs such as the STS, generate inter-individual variability in AV speech perception (Kumar et al., 2020). Collectively, this indicates that occipital alpha activity whether a trial-by-trial state or a stable trait indexed by IAF is tightly coupled to illusion susceptibility.

### 1.2 Neuromodulation and Causal Evidence

Establishing a causal role for alpha and occipital cortex requires direct neuromodulation. Occipital tACS reliably entrains endogenous alpha when delivered at an individual’s own frequency (Helfrich et al., 2014; Zaehle et al., 2010). The clearest causal precedent linking occipital alpha to temporal binding comes from outside the speech domain: Cecere et al. (2015) applied occipital tACS at IAF and at offset frequencies (IAF ± 2 Hz), reporting that slower stimulation enlarged the temporal window of a sound-induced double-flash illusion while faster stimulation significantly reduced it. This pattern has since been replicated across two-flash fusion and segregation tasks (Battaglini et al., 2020; Ronconi et al., 2018; Santoni et al., 2025), visual feature binding (Zhang et al., 2019), and time perception (Mioni et al., 2020), extending the effect to a broader temporal binding window (TBW) construct (Venskus & Hughes, 2021). However, the effect is not universal, the same protocol applied to a suprasecond time-generalisation task found correlational but no causal evidence (Mokhtarinejad et al., 2024), and parietal stimulation has yielded mixed results (Ronconi et al., 2020; Schuhmann et al., 2019).

Previous stimulation-based studies have focused on modulating parietal or multisensory integration (MSI), not occipital area for understanding the causal evidence in case of McGurk illusion. Single-pulse TMS over left pSTS reduces McGurk incidence (Beauchamp et al., 2010), while bilateral cathodal tDCS over STS decreases, and bilateral anodal tDCS over posterior parietal cortex increases, McGurk illusory responses (Marques et al., 2014). This leaves a critical gap: occipital tACS at IAF has repeatedly shaped temporal binding in visual and audiovisual illusions but has never been applied to McGurk illusion speech integration, and McGurk-specific neurostimulation has never targeted occipital cortex. Whether entraining occipital alpha can causally shape susceptibility to an ecologically valid, speech-specific illusion remains untested. A further consideration for occipital tACS studies is distinguishing a selective neuromodulatory effect from a generalised perceptual deficit driven by nonspecific sensory or expectancy confounds (Antal et al., 2017); prior alpha-tACS work has addressed this by showing effects confined to specific conditions rather than a uniform shift in performance (Cecere et al., 2015).

Here, we address this gap by testing whether occipital tACS at IAF and offsets causally alters McGurk susceptibility, using congruent AV stimuli as an internal control against a generalised perceptual deficit and thereby extending causal evidence for alpha-paced temporal binding to naturalistic audiovisual speech. We hypothesised that tACS at IAF and at offset frequencies (IAF ± 2 Hz) would selectively reduce the illusory McGurk percept relative to baseline, while leaving congruent perception largely unaffected. Across a two-day protocol, resting-state EEG was used to determine each participant’s IAF, before personalised occipital tACS was applied at IAF and its offsets while participants categorised congruent and incongruent (McGurk) audiovisual stimuli. Responses were analysed using a Bayesian GLMM to test whether stimulation condition interacted with stimulus type in predicting illusory percept rates. To complement this empirical approach with a mechanistic account of the observed synchronisation dynamics, we additionally employed a computational framework based on the Kuramoto model of coupled oscillators. The Kuramoto model portrays the collective dynamics of interacting oscillatory systems by capturing how oscillators with distinct intrinsic frequencies spontaneously synchronise that yields a quantitative measure of phase coherence (Acebron et al., 2005). Here, each oscillator represents the phase dynamics of a neural population involved in multisensory processing; their interaction depends on coupling strength and time delays. Since tACS can entrain neural oscillations by aligning their specific frequencies, dynamical systems provide a suitable platform to characterise how external stimulation interacts with intrinsic brain dynamics (Vogeti et al., 2022; Madadi Asl & Valizadeh, 2025). Therefore, we employed a Kuramoto model as a parsimonious framework to examine how tACS changes the phase synchronisation and affects the multisensory perception. Synchronisation is quantified using the complex order parameter, whose magnitude ranges from 0, indicating complete desynchronisation, to 1, indicating perfect synchronisation. The model provides a mechanistic framework to relate stimulation-dependent changes in neural phase coordination to experimentally observed variations in illusory perception by looking at how the order parameter changes with the stimulation frequency. Furthermore, comparing the stimulated synchronisation dynamics with behavioural measures of illusion perception enables us to examine the potential relationship between neural phase synchronisation and the probability of experiencing the multisensory illusion.

## 2. Methodology

### 2.1 Experiment

The experiment paradigm developed and used in this study has been built upon a previously designed experiment (Kumar et al., 2020).

#### 2.1.1 Participants

A total of 32 healthy participants were recruited from the Indian Institute of Technology, Jodhpur. One participant was excluded due to a misunderstanding of task instructions, and four were excluded because their congruent performance accuracy fell below 50% on congruent trials in the no-stimulation condition (averaged across all five blocks). The final sample, therefore, comprised twenty-seven participants (10 females) with a mean age of 24.96 (SD=2.75years) and age range from 20-31. Twenty-four of them were right handed, 2 left handed and 1 ambidextrous, as assessed by the Edinburgh Handedness Inventory (Oldfield, 1971). None of the participants had any history of neurological or psychiatric disorders, and they fell under the criteria for administration of tACS (Antal et al., 2017). All participants had normal or corrected-to-normal vision and reported no hearing impairment. Written informed consent was obtained in accordance with the experimental protocol, and ethical clearance was taken from the Institutional Human Ethics Committee (IHEC) of the Indian Institute of Technology, Jodhpur.

#### 2.1.2 Experiment Design and Trials

The experiment spanned two sessions on two different days (average interval: 3.3 ± 2.5 days). On Day 1, a 5-minute resting-state EEG (5min) was recorded to estimate individual alpha frequency (IAF). Participants then completed a multisensory speech perception task, reporting their subjective experiences of incoming audio-visual (AV) stimuli. On Day 2, participants repeated the task while receiving tACS at the occipital area delivered at their respective IAF and IAF ± 2 Hz (offsets) (Figure 2b).

**Figure 1:**
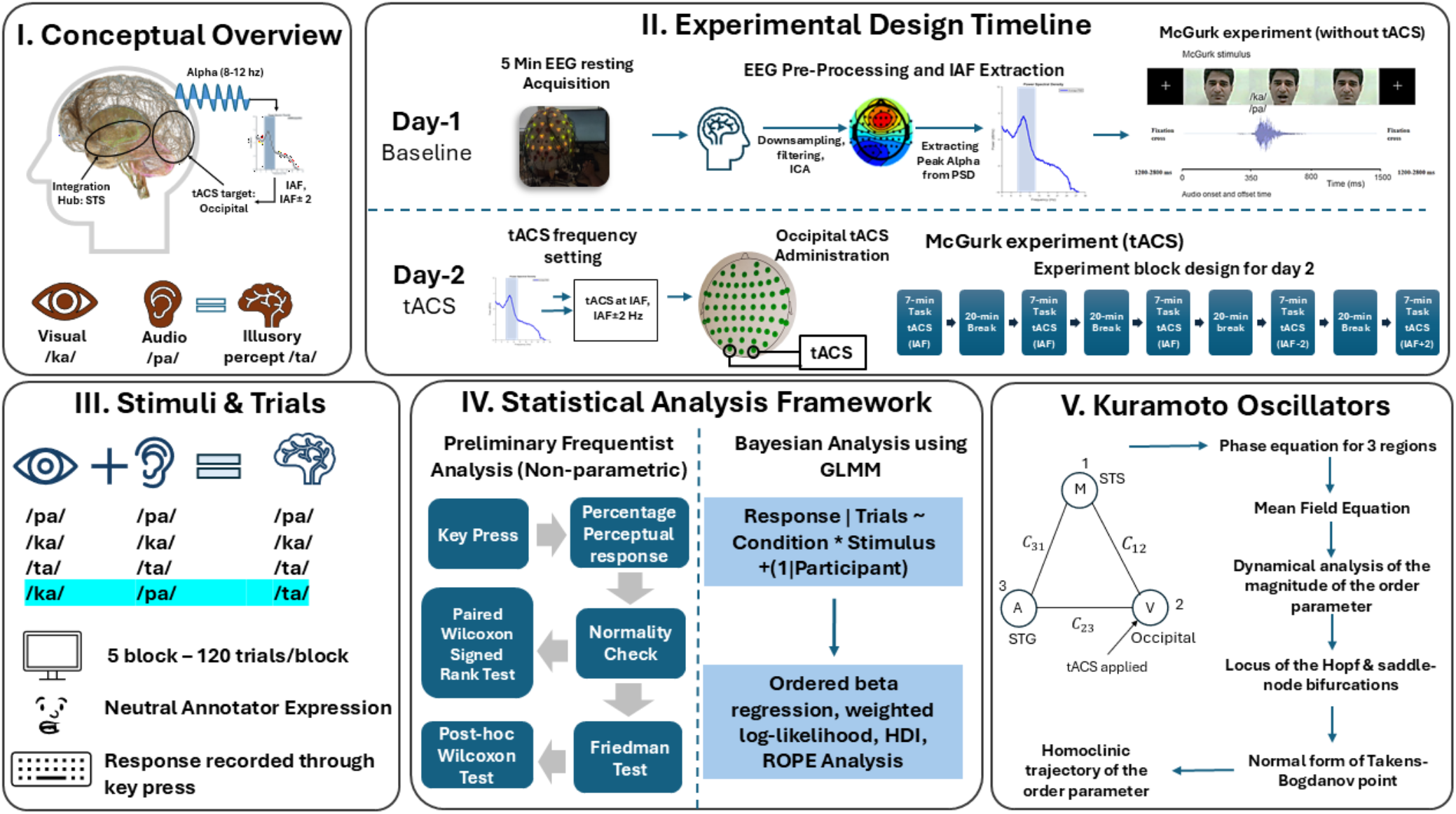
Comprehensive schematic of the experimental design, analytical framework, and theoretical modelling pipeline. (Panel I) Grounded in the theoretical framework linking occipital alpha-band oscillations to multisensory integration, the study evaluates causal changes in the McGurk effect via transcranial alternating current stimulation (tACS). (Panel II) The two-day protocol encompasses baseline individual alpha frequency (IAF) extraction from resting-state EEG (Day 1), followed by personalised occipital tACS administration at IAF and IAF ± 2 Hz across five separated task blocks (Day 2). (Panel III) During these sessions, participants perceive congruent audiovisual syllables and incongruent McGurk stimuli (visual /ka/ and auditory /pa/ eliciting an illusory /ta/ percept). (Panel IV) Behavioural data are subsequently evaluated using a Bayesian Generalised Linear Mixed Model (GLMM) with ordered beta regression, complemented by preliminary non-parametric testing. (Panel V) Finally, a three-node Kuramoto oscillator network mechanistically models the synchronisation dynamics, bifurcations, and homoclinic trajectories underlying individual susceptibility to the multisensory illusion. The person depicted in experimental design pipeline is one of the authors, who provided consent for publication of these images.

**Figure 2:**
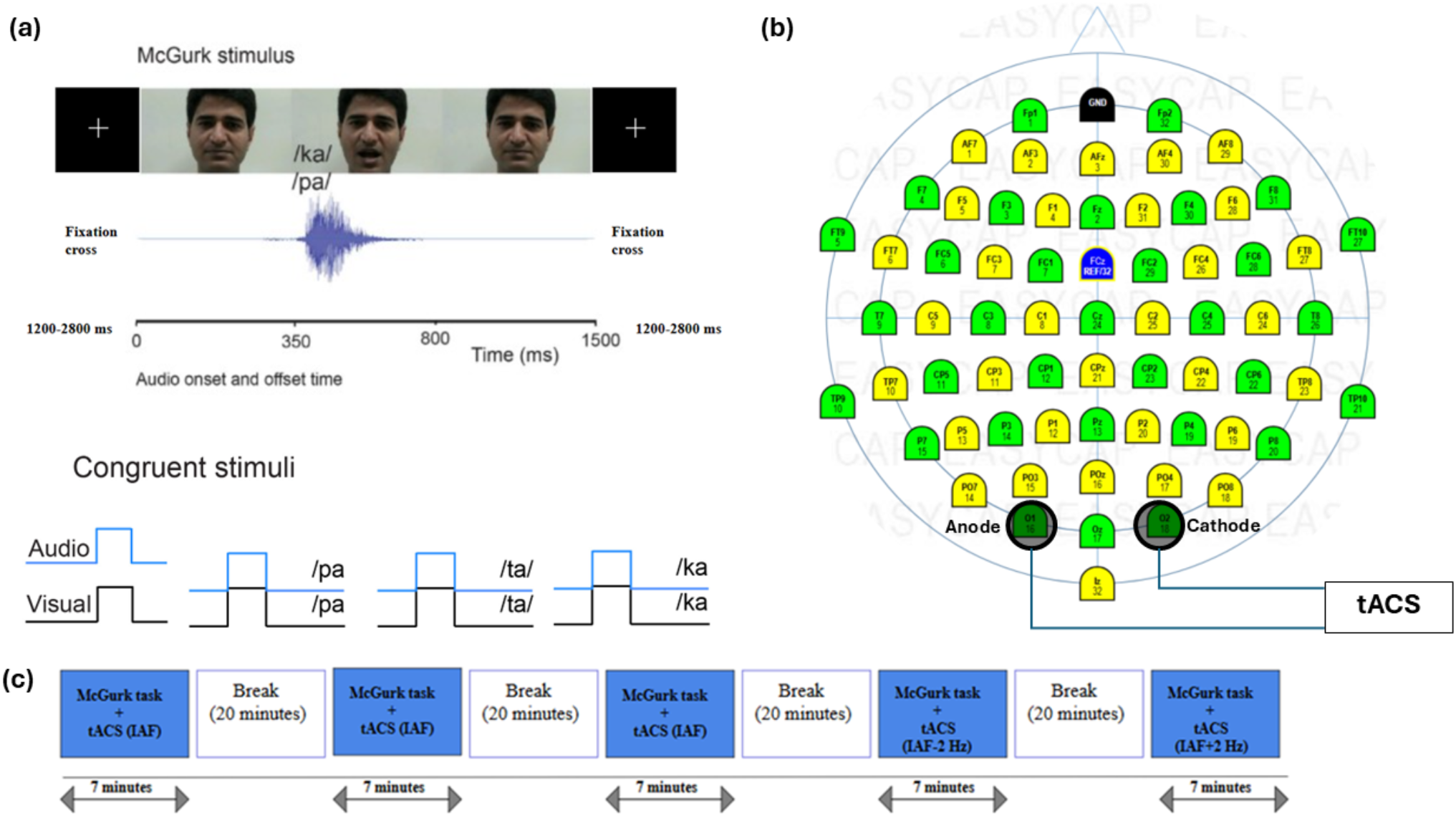
Schematic of the experimental design. (a) Top: Example trial depicting three sequential video frames from the incongruent McGurk stimulus (auditory /pa/ dubbed onto a visual articulation of /ka/), together with the corresponding audio waveform showing audio onset and offset (350–800 ms); the inter-trial interval (fixation cross) was pseudo-randomly varied between 1200 and 2800 ms. Bottom: Schematic timing of the three congruent audiovisual stimuli (/pa/-/pa/, /ta/-/ta/, /ka/-/ka/), illustrating synchronous onset of the auditory and visual syllable in each case, in contrast to the audiovisually mismatched McGurk stimulus above. (b) Electroencephalogram (EEG) montage illustrating the transcranial alternating current stimulation (tACS) electrode placement at occipital sites O1 (anode) and O2 (cathode). (c) Block design timeline detailing the active tACS experimental session on Day 2. The person depicted in experimental design pipeline is one of the authors, who provided consent for publication of these images.

The perception experiment included four kinds of Audio-Visual (AV) stimuli: three congruent (video articulation matching the audio syllable) and one incongruent (mismatch between video articulation and audio syllable). The three congruent syllables used were (/pa/, /ta/, /ka/) and the incongruent (auditory /pa/, visual /ka/) stimulus eliciting the /ta/ illusion (Figure 2a). The experiment was divided into five blocks, each consisting of 120 trials. Out of these 120 trials, congruent and incongruent stimuli were equally divided (30 for each) and randomly presented. The inter-stimulus interval was pseudo-randomised between 1200 ms to 2800 ms to minimise expectancy effects (Minor, 1970). Each participant performed 600 trials in total of each day of the experiment, with a total of 150 incongruent McGurk trials (for one participant (P01), Block 1 of both the Day-1 (wo_tACS) and Day-2 (tACS) sessions failed to register at the response-logging stage; these trials were treated as null and excluded from all condition-level aggregates and analyses. This participant therefore contributed 120 (rather than 150) trials per stimulus to the wo_tACS condition, and 60 (rather than 90) to the tACS_IAF condition). The participants reported their responses through key presses of four keys that denoted the following responses /pa/, /ta/, /ka/ or “something else”.

A within-subject design was implemented in which each participant took part in two sessions of the experiment, Day 1 was without tACS (control) and Day 2 was with tACS. For the second session of the experiment, participants performed the first three blocks while receiving tACS at their respective IAF and the remaining two blocks at offsets: IAF-2 and IAF+2 Hz. Each block had a 20-minute gap during which no stimulation was provided to wash out any carryover from the previous block (Mastakouri et al., 2019). The experiment from the Day 1, lasted for 40-50 minutes and on Day 2, for 2.5 hours, both excluding preparation time. The stimuli were in the form of video of the annotator articulating the syllables. Each trial had one 0 lag audio-video of a Hindi-speaking male annotator, which were preceded and succeeded by a fixation cross. The male annotator had a neutral expression and the AV stimuli appeared in the interval of 300ms-800ms from the appearance of the annotator (Figure 2a).

The stimuli were presented via Presentation® software (Neurobehavioral Systems, Inc., Berkeley, CA, www.neurobs.com) and displayed on a 60 cm LED monitor placed 60 cm from the participants. Articulatory sounds were delivered through H111 Stereo Headset Logitech headphones (Logitech, Lausanne, Switzerland).

#### 2.1.3 Neuromodulation Protocol and tACS Application

Transcranial Alternating Current Stimulation (tACS) was administered via the Nurostym tES device (Brainbox Ltd, Cardiff, UK, www.brainbox-neuro.com). The current was applied through two felt pads of dimensions 5cmX5cm each and were placed over the occipital electrodes, anode at O1 location and cathode at O2 location of a 10-10 montage (see Figure 2b). Each participant was provided with current stimulation at 1.5 mA at their respective Intrinsic Alpha Frequency (IAF) for 30s as a trial to check for any phosphene effect. None of the participants reported any. Each of the 5 blocks of the experiment was approximately 6 minutes and 45 seconds long. tACS was applied to participants for the entirety of the block at their respective IAFs for 3 blocks, and at an offset (IAF-2) for the other two remaining blocks. The fade-in and fade-out of each stimulation were set to 10 seconds each. A 20-minute rest period was provided between each block.

### 2.2 EEG acquisition and pre-processing

On Day 1 of the experiment, 5-minute eyes-closed resting-state EEG data were collected to calculate IAF. 64-channel Ag/AgCl active gel-based electrodes from the Brain Products actiCHamp were placed in a 10-10 montage system using ActiCap snap cap for data collection and recorded using the BrainVision recorder software from Brain Products GmbH, Gilching, Germany. Cz was selected as the reference. The recording sampling rate was kept at 1,000 Hz, with channel impedance check set to 10 kΩ. Pre-processing and IAF calculation were done using EEGLAB(Delorme & Makeig, 2004) and customised MATLAB codes (version R2023b, https://www.mathworks.com/). The data was first downsampled to 250 Hz, then filtered using a Basic FIR filter with a high-pass frequency cut-off at 1 Hz and a low-pass frequency cut-off at 40 Hz. After common average rereferencing, ICA (runica) was performed to remove eye movements and muscle artifact after visual inspection. The preprocessing pipeline has been adapted from EEGLAB (Delorme & Makeig, 2004). Following preprocessing and independent component analysis (ICA), the retained EEG signals were analysed to determine the Individual Alpha Frequency (IAF) for each participant. Power spectral density (PSD) estimates were computed utilising Welch’s averaged, modified periodogram method via MATLAB’s *pwelch* function. Specifically, the time-series data were segmented into 20-second windows with zero overlap, providing a high-frequency resolution suitable for precise peak detection. The resulting power spectra were converted to decibels (dB) and averaged across all channels to construct a single global power spectrum for each subject. Finally, the IAF was identified by extracting the frequency corresponding to the maximum power peak within the canonical alpha band (8–14 Hz). For the power spectral density (PSD) of each individual’s IAF, and average PSD plot refer to Supplementary Material 2 Figure S1(a) and S1(b).

### 2.3 Analysis

Statistical analyses were performed using R v 4.4.3 (R Core Team, 2025). Behavioural responses were recorded as discrete key presses corresponding to the participant’s perceived syllable. To quantify perceptual performance, these raw responses were aggregated and then calculated as Percentage Perceptual Response (PPR) rates for each condition and stimulus type. During this calculation, invalid trials, specifically omitted responses (misses) were excluded. During congruent audiovisual trial only correctly identified trials were scored as a successful attempt leading up to the PPR. For incongruent trials, illusory /ta/ response from participants were considered under the illusory PPR, while the non-illusory /pa/ or /ka/ were considered unifiedly as non-illusory PPR (as key presses response for /ka/ in the incongruent stimuli were extremely low-only 148 trials out of 3980 incongruent McGurk valid trials-summed over all 27 subjects). This approach was taken to ensure the metric accurately reflected true perceptual alterations in the analysis. These aggregated percentage perceptual responses were utilised to compute individual baseline susceptibility to the McGurk illusion and for data visualisation purpose. Finally, to meet the mathematical assumptions of the Bayesian Ordered Beta Regression model (which requires data to be bounded strictly between 0 and 1), these percentage scores (or PPR) were converted into decimal proportions.

To determine the appropriate statistical framework, behavioural response distributions were first assessed for normality using visual inspection of Q-Q plots (See Figure S2, Supplementary Material 2) and the Shapiro-Wilk test. As the percentage perceptual Responses for both tACS and without-tACS (wo_tACS) conditions significantly deviated from a normal distribution (see Table S3, Supplementary Material 3), the data were analysed using a combination of non-parametric frequentist tests and Bayesian modelling.

#### 2.3.1 Behavioural Scoring and Non-Parametric Analysis

To evaluate within condition differences between illusory (McGurk /ta/) and non-illusory (McGurk /pa/-/ka/) responses, paired Wilcoxon signed-rank tests were conducted. And, to assess the effect of tACS conditions on individual response types (e.g., congruent /pa/, /ta/, /ka/, and illusory McGurk responses) across multiple sessions, the Friedman test which is a non-parametric alternative to repeated-measures ANOVA was utilised. For all frequentist analyses yielding main effects, post-hoc pairwise Wilcoxon signed-rank tests were performed. To control for multiple comparisons, all frequentist post-hoc p-values were adjusted using the False Discovery Rate (FDR) method, maintaining the FDR at α = 0.05.

#### 2.3.2 Bayesian Generalised Linear Mixed Modelling (GLMM) for Trial-Weighted Response Proportions

To rigorously overcome the limitations of traditional frequentist tests and specifically to handle the unbalanced trial distributions inherent to the experimental design (i.e., 5 blocks for baseline wo_tACS, 3 blocks for IAF, and 1 block each for IAF-2 and IAF+2) a Bayesian Generalised Linear Mixed Model (GLMM) was implemented to evaluate whether tACS selectively influenced multisensory integration. The model was built using the *brms* (v 2.23.0) Rstan (2.32.7) (Bürkner, 2017, 2018, 2021; Carpenter et al., 2017) and *ordbetareg* (v 0.8; Kubinec 2023, 2025) packages in R.

Traditional linear models, such as Repeated-Measures ANOVA, assume normally distributed data and require balanced designs. They struggle significantly with unbalanced trial distributions, often requiring the aggregation of data which results in a loss of statistical power and trial-level variance (Baayen et al., 2008; Magezi, 2015). In contrast, mixed-effects models natively accommodate unbalanced datasets by utilising maximum likelihood estimation, ensuring that conditions with fewer trials (e.g., the offset stimulation blocks) are appropriately weighted without discarding data (Jaeger, 2008). Furthermore, while standard logistic regression accommodates binomial counts, it can struggle with aggregated continuous proportions (Douma & Weedon, 2019; Ferrari & Cribari-Neto, 2004). Therefore, an Ordered Beta Regression family was applied. Because the design was unbalanced across stimulation frequencies, a weighted likelihood was used, and the ordered beta family accommodated boundary values (0/1) without ad-hoc transformations. This approach natively models proportional data that is highly skewed and accommodates true boundary values (exact 0s and exact 1s) without requiring ad-hoc data transformations (Kubinec, 2023).

The formal model structure was specified as:

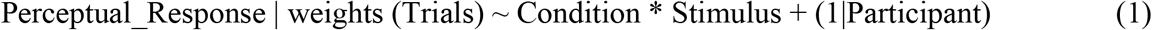

Where, Perceptual_Response holds the values of PPR converted to decimal format to have the values between 0 to 1.

This computational syntax corresponds to a generalised linear mixed-effects formulation where the expected perceptual response proportion for observation *i* and participant *j (μ_ij_)* is modelled via a logit link function. Specifically, the logit function transforms the bounded perceptual response proportions into unbounded log-odds *(η_ij_)*, thereby preserving the linearity of the predictors and constraining predictions to the valid probability space [0, 1]. The linear predictor is formally defined as:

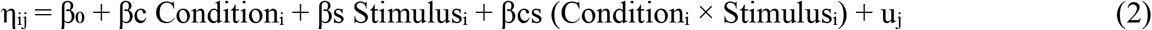

In this equation, *β*_*0*_ represents the global intercept, indexing the baseline condition (i.e., the absence of stimulation, wo_tACS, paired with the /ka/ stimulus). The *βc* and *βs* coefficients quantify the population-level main effects of the tACS condition and the phonetic stimulus, respectively. Crucially, *βcs* denotes the interaction term, which tests the primary hypothesis regarding whether the modulatory influence of tACS varies significantly as a function of the presented stimulus. Furthermore, the *u_j_* term incorporates a random intercept for each participant to account for repeated measures and inter-individual variability in baseline susceptibility to the McGurk illusion, with the assumption of a normal distribution where *u_j_ ∼ N (0, σ^2^ u)*. Finally, the model accommodated the unbalanced block design through a weighted log-likelihood approach. The mathematical contribution of each observation to the posterior was multiplied by its respective trial count *(w_i_)*, ensuring that condition estimates derived from a larger number of trials exerted a proportionally stronger influence on the model’s parameters than those derived from fewer trials.

To prevent overfitting and penalise large effect sizes, a weakly informative, regularising prior with a normal distribution centred at zero with a standard deviation of 1.5 was applied to all population-level β coefficients. As the model operated on the logit scale, a standard deviation of 1.5 restricts the model from making extreme, mathematically improbable predictions while remaining sufficiently broad to allow the observed data to dominate the posterior distribution (Gelman et al., 2008).

The model was fitted using Markov Chain Monte Carlo (MCMC) sampling via the Stan probabilistic programming language, running 4 chains with 4000 iterations each (total 8000 warmups). Post-hoc Bayesian inference was conducted using Estimated Marginal Means (*emmeans*) (v 2.0.2; Lenth and Piaskowski 2026) to extract posterior distributions for the condition-by-stimulus interactions. Differences between conditions were evaluated using 95% Highest Density Intervals (HDI) and Region of Practical Equivalence (ROPE) tests via the *bayestestR* package (v 0.17.0; Makowski, Ben-Shachar, Chen, et al. 2019; Makowski, Ben-Shachar, and Lüdecke 2019). Pairwise and interaction contrasts (difference of differences) were computed on the log-odds scale to avoid probability-boundary compression artifacts and were considered credibly different if their 95% HDIs strictly excluded zero. To quantify the strength of evidence for these effects, Bayes factors were computed using the Savage–Dickey density ratio method (Wagenmakers et al., 2010): parameter-level BFs were obtained via *bayestestR::bayesfactor_parameters()*, and BFs for all pairwise and interaction contrasts were derived from the equivalent ratio returned by *brms::hypothesis()* (Bürkner, 2017), inverting the reported evidence ratio (BF_01_) to yield BF_10_. These BFs were interpreted as follows: a BF between 1 (log_10_(BF) = 0) and 3 (log_10_(BF) = 0.477) was considered as anecdotal evidence while a BF more than 3 (log_10_(BF) > 0.477) moderate evidence in favour of the effect (i.e., in favor of the alternative hypothesis H_1_ that tACS credibly modulates multisensory integration), a BF greater than 10 (log_10_(BF) > 1) was considered strong evidence, and a BF greater than 100 (log_10_(BF) > 2) was considered high evidence. Conversely, BFs between 1 (log_10_(BF) = 0) and 1/3 (log_10_(BF) = -0.477) were anecdotal evidence while a BF less than 1/3 (log_10_(BF) < -0.477), 1/10 (log_10_(BF) < -1), and 1/100 (log_10_(BF) < -2) were interpreted as moderate, strong, and high evidence against the effect (in favour of the null hypothesis H_0_-“tACS has no effect on multisensory integration”), respectively. Because raw BF values for several comparisons spanned many orders of magnitude, we report log_10_(BF_10_) throughout the Results for readability and consistency with the corresponding figures, where positive log_10_(BF_10_) indicates evidence favouring the effect and negative log_10_(BF_10_) indicates evidence favouring the null.

In addition to the BF based inferential framework, we report approximate frequentist p-value for each post-hoc contrast to aid interpretation and to permit false-discovery-rate correction (Benjamini-Hochberg) across the family of comparisons within each analysis. For contrasts obtained via *hypothesis()* in *brms* (Bürkner, 2017), p-values were derived from the probability of direction (pd) the proportion of the posterior distribution with the same sign as the median computed directly on the posterior draws underlying each contrast using the *p_direction()* function from the *bayestestR* package (Makowski, Ben-Shachar, Chen, et al., 2019). Following the same source, pd was converted to an approximate two-sided frequentist-equivalent p-value via p = 2 × (1 - pd). For contrasts obtained via *emmeans* (Lenth et al., 2026), we applied the same procedure to the posterior draws of each contrast, extracted using the *tidybayes* package, rather than relying on a normal-theory approximation of the posterior mean and standard deviation so that the resulting p-value reflects the empirical shape of each contrast’s posterior rather than an assumed Gaussian form. As these p-values are computed from models fit to a large number of trials per cell, several reach extremely small magnitudes; these are reported as p < 0.001 rather than at full precision.

#### 2.3.3 Phenomenological model to understand the dynamics of illusory speech perception in tACS modulation

In order to understand the dynamics of McGurk illusory /ta/ responses for each participant, we employed Kuramoto oscillator model for three brain regions (STS, STG and Occipital). The superior temporal sulcus (STS), superior temporal gyrus (STG), and occipital cortex were selected as functionally complementary nodes that represent audiovisual integration, auditory speech processing, and visual speech processing, respectively. Previous neuroimaging and causal studies have implicated these regions in McGurk perception (Beauchamp et al., 2010; Calvert et al., 2000; Nath & Beauchamp, 2012). Moreover, dynamical modeling studies show that interactions and synchronization among sensory and multisensory brain regions explain why people perceive multisensory information differently (Thakur et al., 2016; Kumar et al., 2020). Because transcranial alternating current stimulation (tACS) modulates brain waves through frequency-dependent entrainment (Vogeti et al., 2022; Madadi Asl & Valizadeh, 2025), we used a three-node Kuramoto oscillator model (Figure 3). This model-comprising the STS, STG, and occipital cortex-characterizes how individual participants’ McGurk illusory /ta/ responses change under tACS stimulation. The equations of these three brain regions can be written as follows

**Figure 3:**
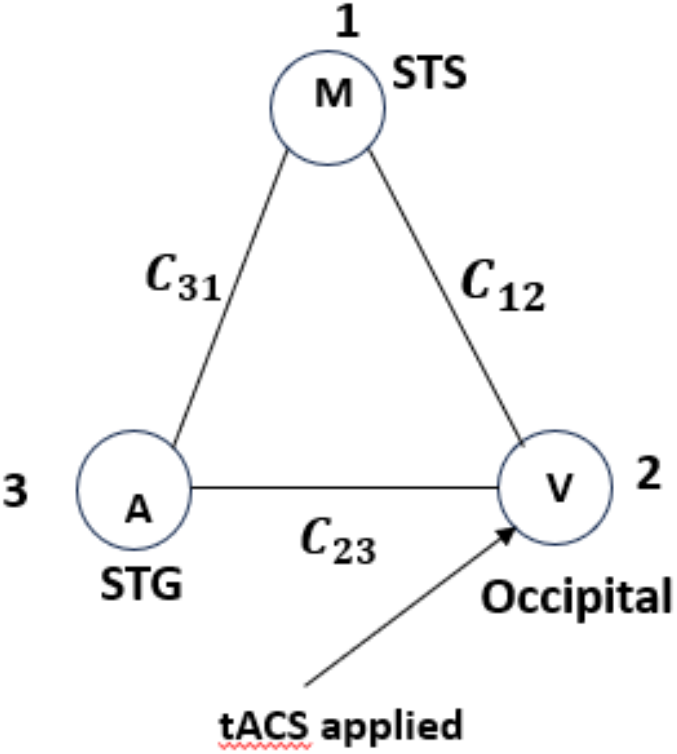
The schematic diagram of three brain regions represented by three Kuramoto oscillators. STS denotes superior temporal sulcus which is multisensory integration region, occipital is the visual region which is perturbed by tACS, STG denotes superior temporal gyrus which is auditory region. *C*_12_ denotes structural connectivity (fibre density) in between STS and occipital regions, *C*_23_ denotes structural connectivity (fibre density) in between STG and occipital regions, *C*_31_ denotes structural connectivity (fibre density) in between STG and STS regions.

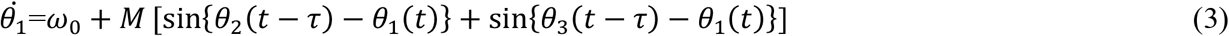

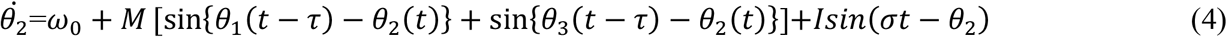

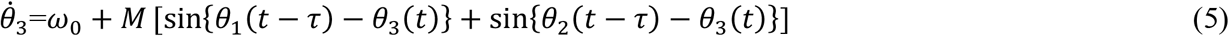

*Where, θ*_1_, *θ*_2_ and *θ*_3_ are the phases of STS, occipital and STG areas respectively. *τ* is the neurophysiological delay between two brain areas which we have kept the same for all these three brain nodes. We have also maintained identical intrinsic frequency for the three brain areas denoted by *ω*_0_ to avoid additional complexity. Since tACS perturbation were targeted to occipital region, we have applied a sinusoidal current to the occipital region with amplitude *I* and frequency *σ*. These three brain regions are interconnected through coupling strength and structural connectivity which is *M. M* can be written as follows

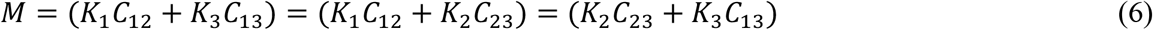

Here, *K*_1_ is the coupling strength between occipital and STS, *K*_2_ is the coupling strength between STG and occipital, and *K*_3_ is the coupling strength between STS and STG. Now, to understand the collective dynamics of three brain regions, we can formulate a single mean field equation as:

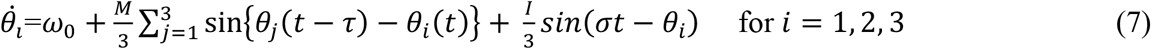

Here, *i* denotes the index of the brain region. We shift the system into a frame co-rotating with the stimulation frequency for analyzing the synchronization dynamics relative to the applied tACS. This transformation removes the explicit time dependence of the periodic stimulation term. It converts the dynamics into an autonomous system that describes the relative phase between the neural oscillators and the external tACS drive that facilitates the analysis of synchronization states. To achieve this, let

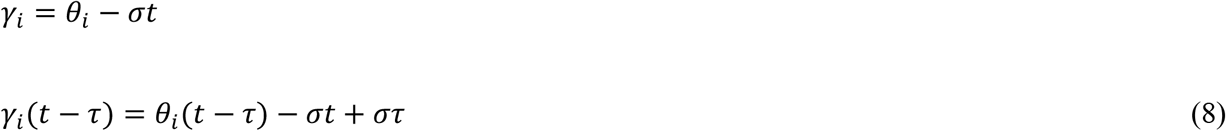

Substituting eq. (8) into eq. (7), we obtain

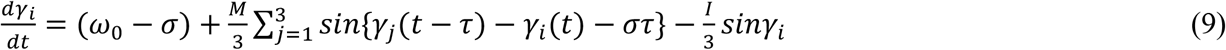

Now, we aim to understand the dynamics of order parameter *r*(*t*), which can be related to participant’s propensity towards McGurk illusion perception. When the magnitude of the order parameter approaches 1, the illusion is maximal, whereas when the order parameter approaches zero, the illusion is minimal. We can derive the following equation (see Supplementary Material 1 for the detailed derivation)

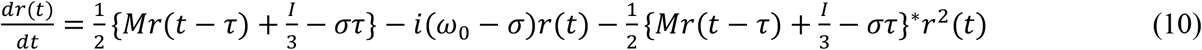

Here, * denote the complex conjugate. Let

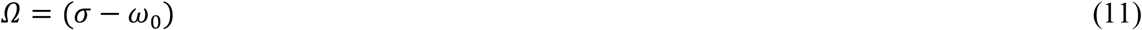

denotes detuning between tACS frequency and intrinsic alpha frequency (IAF) of each participant. *ω*_0_ would be different for each participant. Since order parameter is a complex quantity, we can write in polar coordinates,

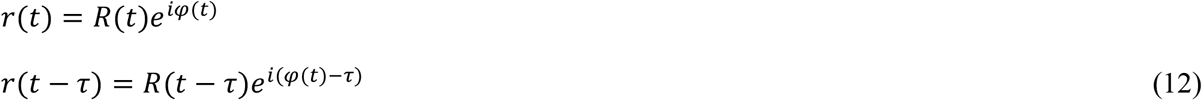

Here, *R*(*t*) and *φ*(*t*) are the magnitude and phase of the order parameter respectively. Substituting eq. (12) into eq. (10), then separating real and imaginary parts, we can write

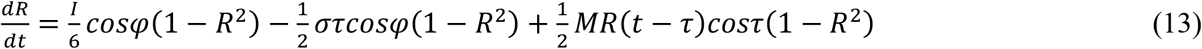

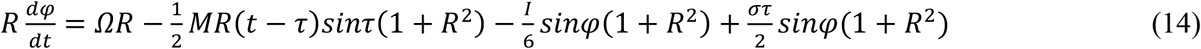

It is algebraically cumbersome to solve for the fixed points directly from eq. (13) and eq. (14). Instead, we can make progress by imposing appropriate bifurcation conditions and solving for the parameters in terms of the fixed points. To obtain mathematically tractable solutions, we vary only *Ω* while keeping *M* and *τ* fixed. This approach allows us to identify the stimulation frequency-dependent transitions between distinct dynamical regimes and thereby relate the bifurcation structure of the model to experimentally observed changes in McGurk /ta/ responses. Let us first find the locus of Hopf bifurcation where at steady state, 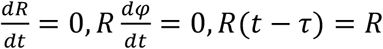 and the trace of Jacobian is zero.

From equations (13), (14), and (11) and putting Ω = Ω_*Hopf*_, we get,

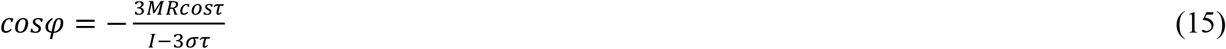

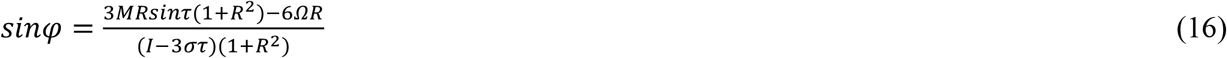

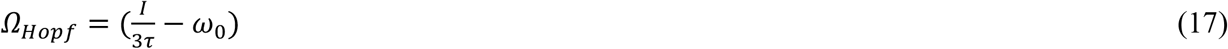

From eq. (17), we see that detuning along Hopf curve is independent of the order parameter. Next, let us find the locus of saddle-node bifurcation where the determinant of Jacobian is zero. Similarly, from equations (13), (14), and (11) and putting Ω = Ω_*SN*_, we get,

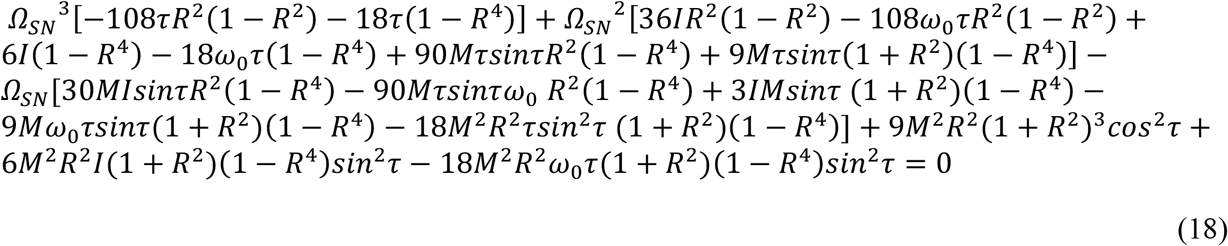

From eq. (18), we see that detuning along the saddle-node curve is dependent on the order parameter. Equation (18) is a cubic equation of Ω_*SN*_ and we can find the three roots of Ω_*SN*_ by Cardano’s approach (Lestari et al., 2020), then plot *R* versus these three branches of Ω_*SN*_ for each participant with their own IAF (by keeping *M* and *τ* fixed) to understand the variation of this order parameter. In the absence of tACS stimulation, the external input can be removed by setting *I* = 0 in eq. (18). The order parameter can then be computed for each participant individually using their corresponding IAF, while keeping *M* and *τ* fixed. When eq. (17) and eq. (18) intersects, they form a Takens-Bogdanov (TB) bifurcation point (Kuznetsov, 2004), from which a homoclinic manifold emerges along which the order parameter or the participants’ illusory responses can vary. From equations (11), (13) and (14), we can derive (see Supplementary Material 1 for the detailed derivation) the normal form of TB as follows

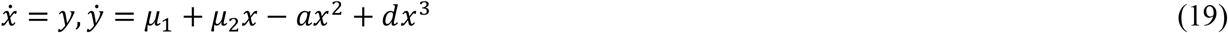

Where *μ*_1_ = Ω − Ω_*TB*_, *μ*_2_ = *σ* − *σ*_*TB*_, *a* = *MR*_*TB*_ *and* 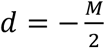

The equation of the homoclinic trajectory (derivation provided in Supplementary Material 1) is as follows

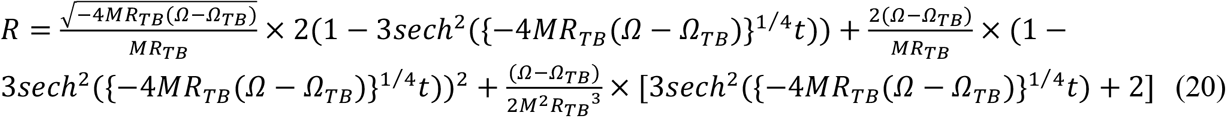

Now, we assume coupling strength is inversely proportional to the detuning frequency, that is when tACS frequency matches with the participants’ IAF, the coupling strength is maximum. Therefore, we can write,

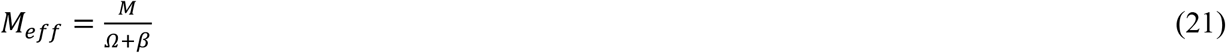

Where *M*_*eff*_ is the effective coupling strength and *β* is the damping parameter. Substituting eq. (21) into eq. (7), we can write the mean field equation as follows

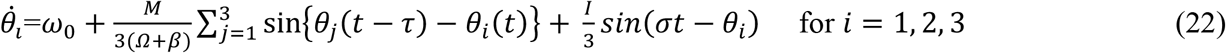

Here, *i* denotes the index of the brain region. For some participants, the behavioural data revealed a higher percentage of McGurk /ta/ responses at (IAF-2) than at (IAF+2) (See Table 4 in Supplementary Material 4), indicating a participant-specific asymmetry around the individual alpha frequency (IAF). To account for this asymmetry, we incorporated *M*_*eff*_ into eq. (18) and eq. (20) to determine the locus of the saddle node bifurcation, obtained from a quintic polynomial in Ω_*SN*_ and the homoclinic trajectory for each participant, respectively.

The Kuramoto model fitting was divided into two steps: % of McGurk /ta/ response at (IAF+2) is greater than or equal to at (IAF-2) (for 17 participants), and % of McGurk /ta/ response at (IAF-2) is greater than at (IAF+2) (for 10 participants) (see Supplementary Material 4). The order parameter (R) value for each of the 17 participants was uniquely fitted using the proportion of McGurk /ta/ responses, with participant-specific M and *τ* values (see Table 1 and Table 2 in supplementary material 4). Similarly, the order parameter (R) value of the 10 participants was fitted with the proportion of McGurk /ta/ responses with the same M, *τ*, and *β* values for each participant (see Table 3 and Table 4 in Supplementary Material 4) since all other M, *τ*, and *β* values provide almost similar R values for these 10 participants. MATLAB 2024a has been used to calculate all R values and simulation results.

**Table 1.**
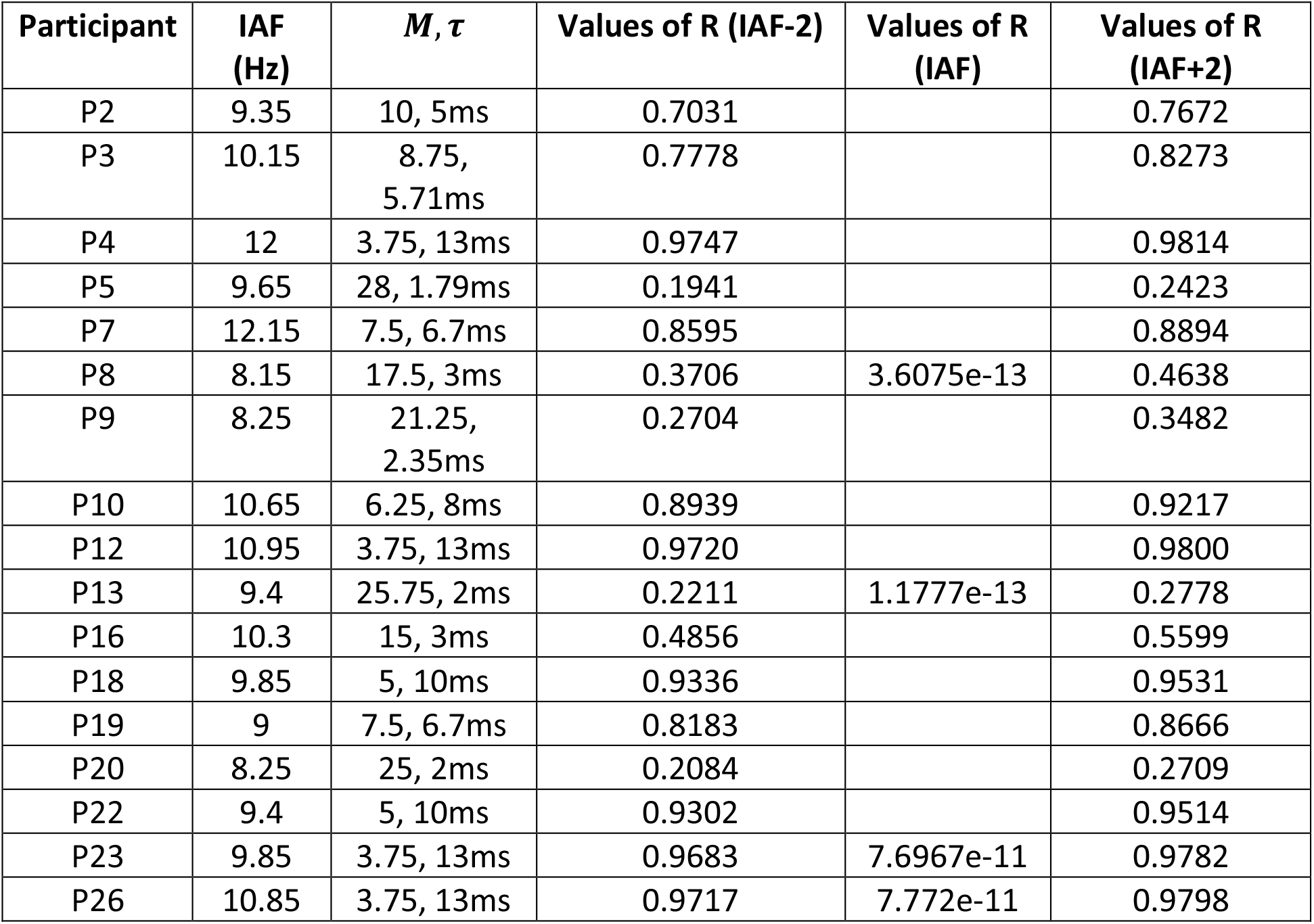
(order parameter (R) values)

**Table 2.**
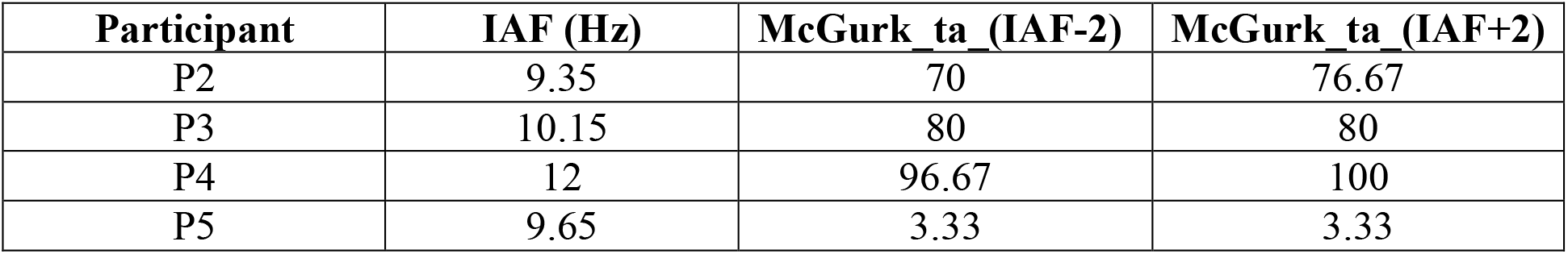
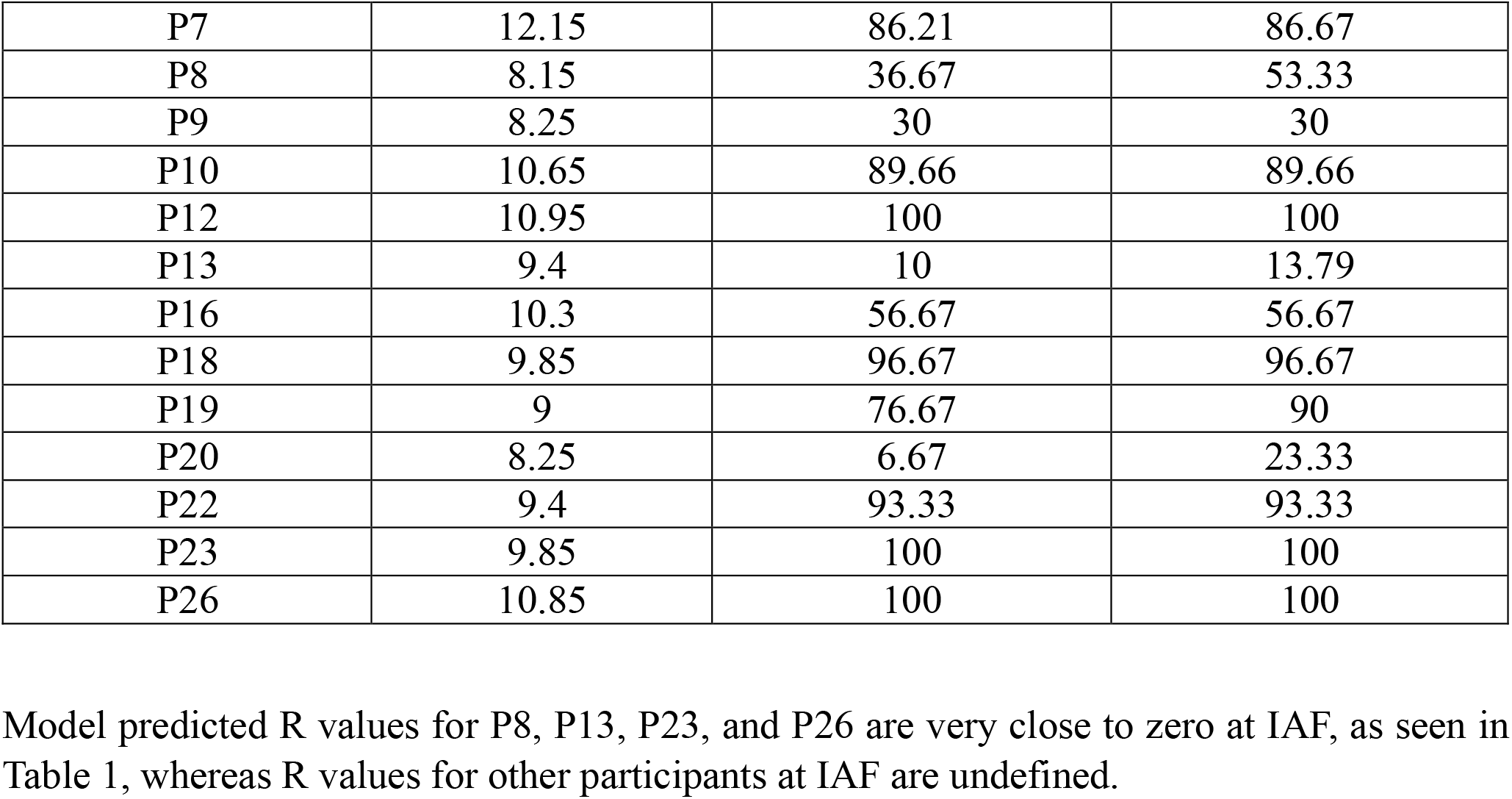
(%/ta/ responses)

| Participant | IAF (Hz) | McGurk ta (IAF-2) | McGurk ta (IAF+2) |
| --- | --- | --- | --- |
| P2 | 9.35 | 70 | 76.67 |
| P3 | 10.15 | 80 | 80 |
| P4 | 12 | 96.67 | 100 |
| P5 | 9.65 | 3.33 | 3.33 |
| P7 | 12.15 | 86.21 | 86.67 |
| P8 | 8.15 | 36.67 | 53.33 |
| P9 | 8.25 | 30 | 30 |
| P10 | 10.65 | 89.66 | 89.66 |
| P12 | 10.95 | 100 | 100 |
| P13 | 9.4 | 10 | 13.79 |
| P16 | 10.3 | 56.67 | 56.67 |
| P18 | 9.85 | 96.67 | 96.67 |
| P19 | 9 | 76.67 | 90 |
| P20 | 8.25 | 6.67 | 23.33 |
| P22 | 9.4 | 93.33 | 93.33 |
| P23 | 9.85 | 100 | 100 |
| P26 | 10.85 | 100 | 100 |

**Table 3.** (order parameter (R) values for M= 10, *τ* =5 ms and *β* = 1)

| Participant | IAF (Hz) | Values of R (IAF-2) | Values of R (IAF) | Values of R (IAF+2) |
| --- | --- | --- | --- | --- |
| P1 | 8.65 | 0.9995 | 0.0194 | 0.9974 |
| P6 | 11.2 | 0.9997 | 0.0219 | 0.9980 |
| P11 | 10.75 | 0.9995 | 0.0214 | 0.9978 |
| P14 | 9.45 | 0.9995 | 0.0204 | 0.9976 |
| P15 | 10.4 | 0.9995 | 0.0214 | 0.9978 |
| P17 | 9.3 | 0.9997 | 0.0199 | 0.9976 |
| **P21** | **9.2** | **0.9948** | **0.0199** | **0.9976** |
| P24 | 11.55 | 0.9995 | 0.0224 | 0.9980 |
| P25 | 10.95 | 0.9995 | 0.0219 | 0.9979 |
| P27 | 8.2 | 0.9995 | 0.0189 | 0.9973 |

**Table 4.**
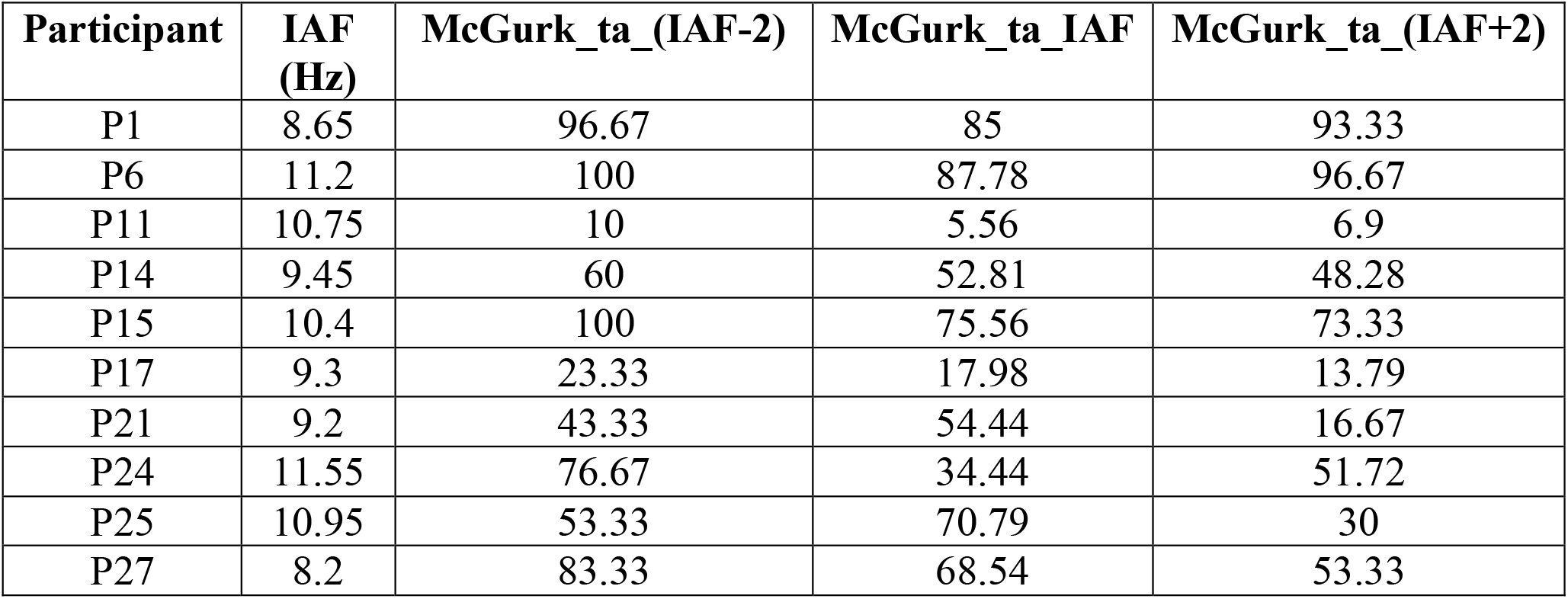
(% /ta/ responses)

| Participant | IAF (Hz) | McGurk_ta_(IAF-2) | McGurk_ta_IAF | McGurk_ta_(IAF+2) |
| --- | --- | --- | --- | --- |
| P1 | 8.65 | 96.67 | 85 | 93.33 |
| P6 | 11.2 | 100 | 87.78 | 96.67 |
| P11 | 10.75 | 10 | 5.56 | 6.9 |
| P14 | 9.45 | 60 | 52.81 | 48.28 |
| P15 | 10.4 | 100 | 75.56 | 73.33 |
| P17 | 9.3 | 23.33 | 17.98 | 13.79 |
| P21 | 9.2 | 43.33 | 54.44 | 16.67 |
| P24 | 11.55 | 76.67 | 34.44 | 51.72 |
| P25 | 10.95 | 53.33 | 70.79 | 30 |
| P27 | 8.2 | 83.33 | 68.54 | 53.33 |

Overall, the derived equations provide the loci of the Hopf, saddle node bifurcations, and the homoclinic trajectory as functions of the order parameter, coupling strength, neurophysiological delay, and detuning frequency. These expressions show the limits of different states in the A-V-AV network. They also reveal how tACS frequency disrupts synchronization. In the context of McGurk susceptibility, variation of tACS frequency relative to these dynamical boundaries may lead to enhanced or reduced phase synchronization, which corresponds to increased or decreased probabilities of illusory /ta/ responses, respectively. The participant-specific location of these dynamical boundaries can therefore provide a mechanistic explanation for the observed variability and asymmetry in McGurk responses around the individual alpha frequency.

## 3 Results

### 3.1 Behavioural outcomes: High congruent accuracy and baseline illusion susceptibility

During the no-stimulation condition, we observed that the Percentage Perceptual Response (PPR) for congruent AV-stimulus syllables /ta/ and /ka/ were 96.07% (SD = 8.322) and 97.21% (SD = 1.856), respectively, while for /pa/ was 85.2% (SD = 11.289). However, during tACS conditions (both at IAF and offsets: IAF+2 and IAF-2), we observed PPR were 96.74% (SD = 4.085) (/ta/), 97.84% (SD = 2.055) (/ka/), and 88.52% (SD = 12.503) (/pa/) (Table S1, Supplementary Material 3), averaged across participants. A Friedman test confirmed that PPR differed significantly across the five perceptual response categories (Mg_pa_ka, Mg_ta, ka_ka, pa_pa, ta_ta) within each stimulation condition (wo_tACS: χ^2^ = 81.101, W = 0.751; IAF: χ^2^ = 76.949, W = 0.712; IAF-2: χ^2^ = 61.916, W = 0.573; IAF+2: χ^2^ = 64.710, W = 0.599; all df = 4, all p_adj<0.001; Table S4, Supplementary Material 3) followed by a post-hoc pairwise Wilcoxon signed rank test for pairwise comparisons of PPR corresponding to different stimulus. PPR for congruent /pa/ were significantly lower than both congruent /ta/ and /ka/ (vs. /ta/: wo_tACS V = 32.0, p_adj = 0.0004; IAF V = 57.0, p_adj = 0.0020; IAF-2 V = 47.0, p_adj = 0.0140; IAF+2 V = 28.0, p_adj = 0.0060; vs. /ka/: wo_tACS V = 348.0, p_adj < 0.0001; IAF V = 305.0, p_adj = 0.0002; IAF-2 V = 252.0, p_adj = 0.0009; IAF+2 V = 170.0, p_adj = 0.0040;; see Table S5, Supplementary Materials 3) in the baseline (wo_tACS) and all tACS conditions (IAF and offsets), with no statistical difference between congruent /ta/ and /ka/ (wo_tACS: V = 132.5, p_adj = 0.627; IAF: V = 144.0, p_adj = 0.578; IAF-2: V = 95.0, p_adj = 0.168; IAF+2: V = 79.0, p_adj = 0.584) (**Figure 4**). Overall for congruent conditions, the difference in PPR that we observed for no stimulation condition aligns with previous literature (Kumar et al., 2020).

**Figure 4:**
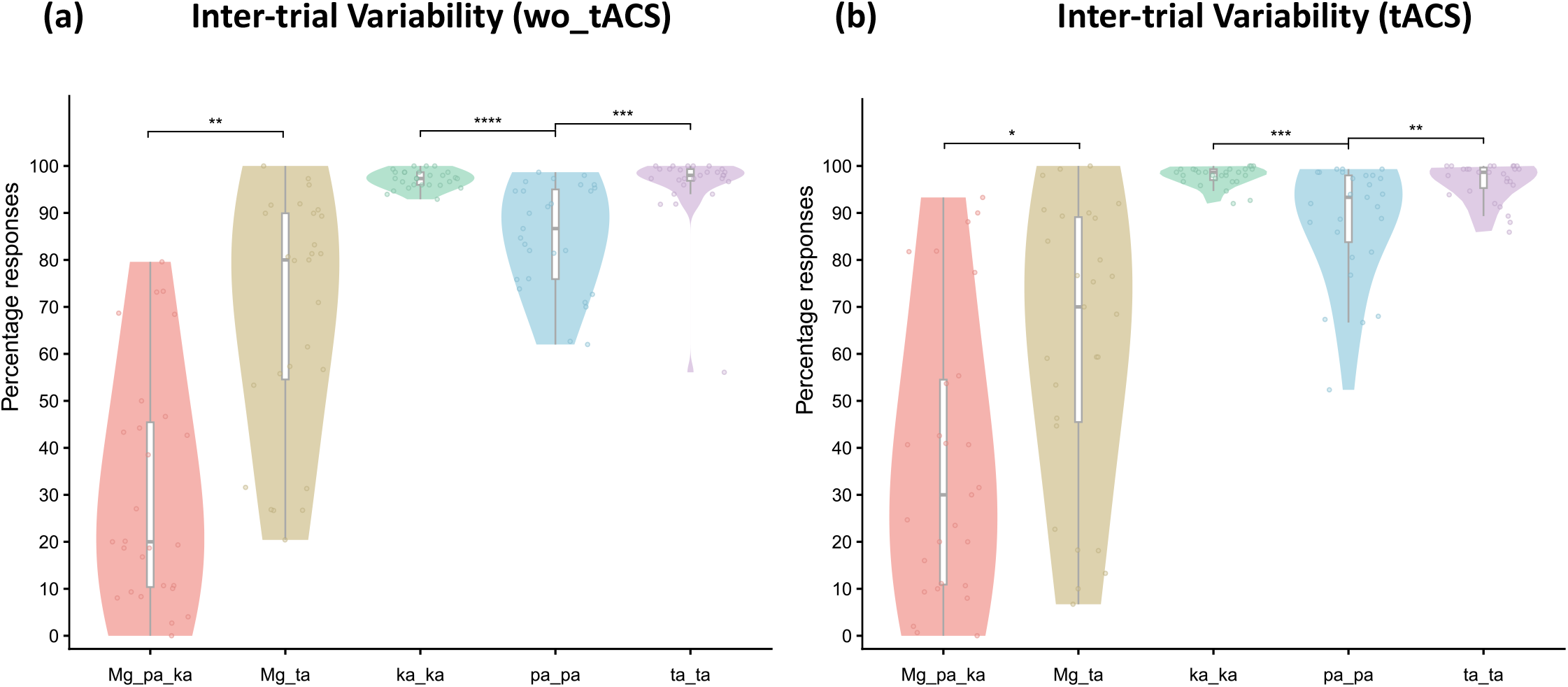
Inter-trial variability for all stimulus conditions. Violin plots illustrating the percentage of illusory (/ta/) and unisensory (/pa/) percepts during the presentation of incongruent McGurk stimuli, alongside the percentage of accurate perceptual responses for congruent audiovisual stimuli (/pa/, /ta/, and /ka/). (a) Data are averaged across all participants for the baseline control condition (wo_tACS) and (b) All tACS conditions (IAF and offsets). Y-axis represents percentage responses and X-axis denotes congruent and incongruent stimulus. FDR-corrected p-values from all ten pairwise comparisons per condition. For separate plots of IAF and offsets refer to Figure S4, Supplemntary Material 2.

Additionally, the difference between illusory /ta/ and non-illusory /pa_ka/ during incongruent McGurk trials was significant without tACS (V = 324, p_adj = 0.005) and tACS at IAF-2 (V = 287, p_adj = 0.038) but not under tACS at IAF (V = 248, p_adj = 0.067) or IAF+2 Hz (V = 275, p_adj = 0.053 (**Figure 5**). The median of the PPR of McGurk /ta/ i.e., illusory percept, without tACS was 80.0% (SD = 25.914), with tACS at IAF was reported to be 70.0% (SD = 29.653), at IAF+2 at 73.33 (SD = 34.062) and IAF-2 at 76.67 (SD = 33.713) indicating a shift in the illusory perception among the participants (see Table S2, Supplementary Material 3), from more illusory perception in the case of no stimulation to lower illusion perceived upon application of neurostimulation, lowest being at IAF (**Figure 6**). Furthermore, Friedman’s test was performed on the percentage response of both congruent and incongruent stimuli with stimulation as the independent variable. A numerical decrease in illusory response was observed across active stimulation conditions relative to baseline; however, this did not reach significance in the omnibus test (χ^2^ = 3.836, p_adj = 0.360, df = 3, Kendall_W = 0.047), and post-hoc pairwise Wilcoxon signed rank tests (FDR corrected) similarly revealed no significant pairwise differences (p_adj) = 0.226, 0.722, 0.449) for IAF, IAF-2 and IAF+2 respectively (refer Table S7 & S8, Supplementary Material 3). Additionally, there was no significant difference in the percentage response of congruent stimuli across the stimulation conditions except for congruent /pa/, which interestingly showed a significant increase in PPR for tACS at IAF (V = 66.0, p_adj = 0.013) condition as compared to without stimulation. No such significant difference was observed for tACS at offsets (See Figure S5, Supplementary Material 2; full statistical output in Table S7-S8, Supplementary Material 3).

**Figure 5:**
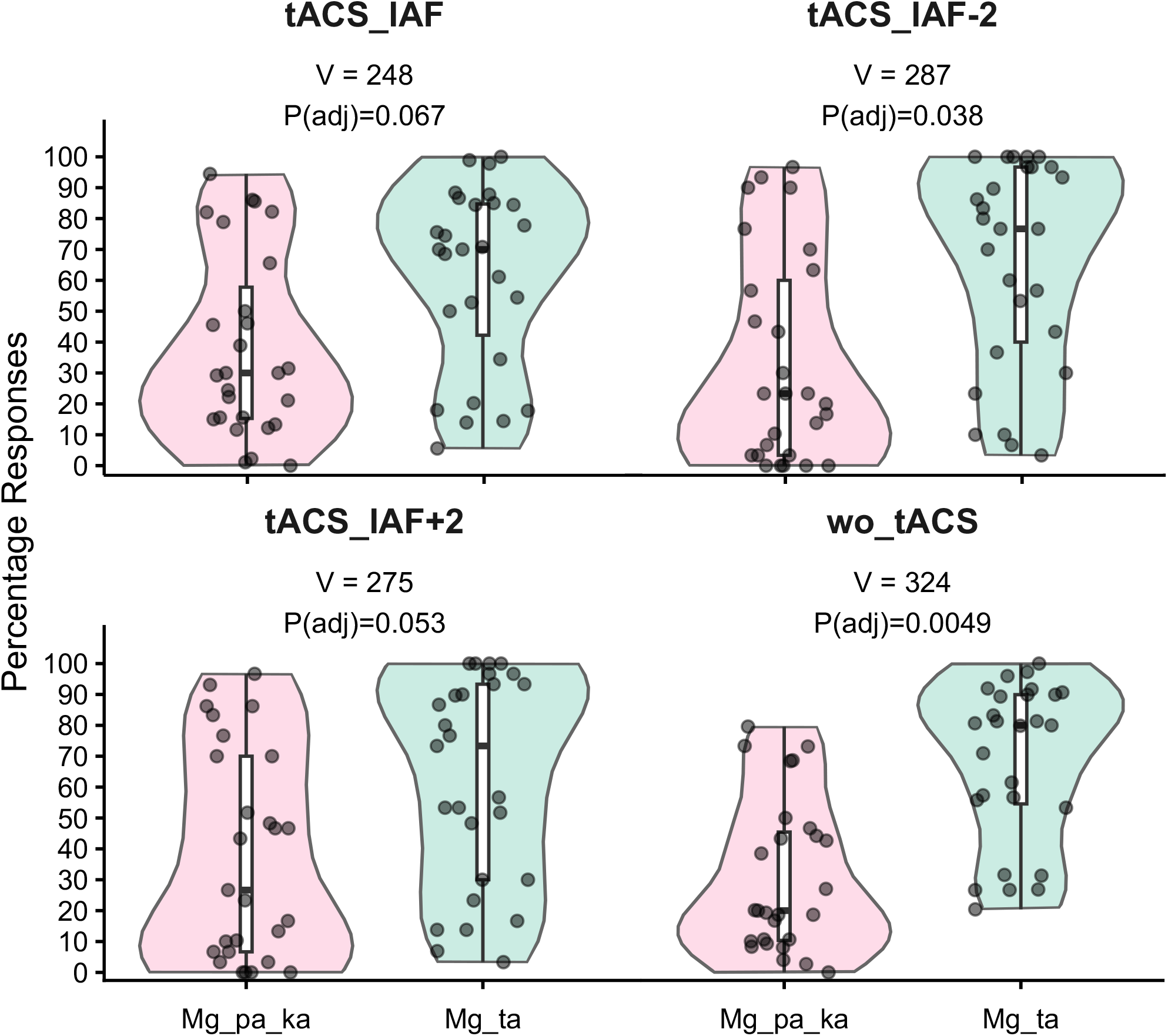
Inter-trial variability for incongruent stimuli. Distributions of illusory (/ta/) versus non-illusory (/pa/ or /ka/) perceptual responses during the presentation of the incongruent McGurk stimulus. Data are contrasted across the baseline control condition (wo_tACS) and all active stimulation conditions (tACS at IAF, IAF-2 Hz, and IAF+2 Hz). Y-axis represents percentage responses and X-axis denotes non-illusory (Mg_pa_ka) responses and illusory (Mg_ta) responses. FDR-corrected p-values across the four focal comparisons. Refer to Table S6, Supplementary Material 3, for full Wilcoxon test statisitcs.

**Figure 6:**
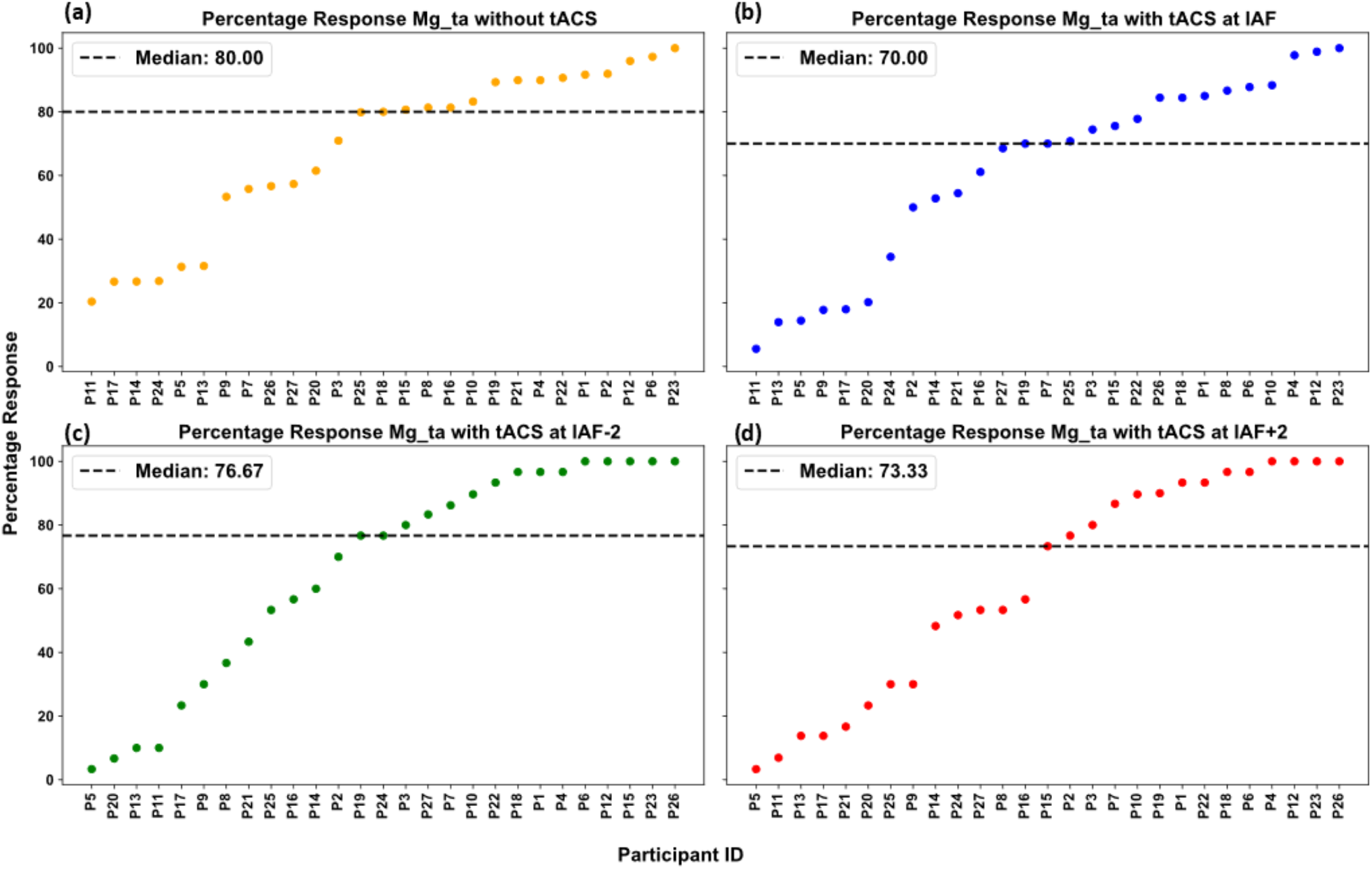
Inter-individual variability in McGurk illusion susceptibility. The propensity for the McGurk effect across all 27 participants is expressed as the percentage of illusory /ta/ percepts (Mg_ta) during the presentation of incongruent audiovisual stimuli. Responses are shown across the baseline condition (without tACS) and active stimulation conditions (tACS at IAF, IAF-2 Hz, and IAF+2 Hz). The y-axis represents the percentage of perceptual responses, and the x-axis displays participant identifiers sorted in ascending order of their baseline illusory response rate.

Despite the non-significant omnibus Friedman result for illusory responses, the Bayesian GLMM analysis of the same response proportions provided high evidence for a stimulation effect on the McGurk illusion (Section 3.2). This apparent discrepancy likely reflects the substantially lower statistical power of the omnibus non-parametric test relative to the Bayesian GLMM. The Friedman test operates on condition-level aggregated percentages across trials within each participant, treating the four conditions as equally weighted, non-independent ranks, whereas the ordered-beta GLMM models trial-count-weighted proportions directly and explicitly weights each condition according to its actual trial count (5 baseline blocks vs. 3 blocks for IAF and 1 block each for the offset conditions) via participant-level random effects. The GLMM’s use of the full trial-level dataset, rather than session-averaged ranks, is therefore better powered to detect the same underlying effect.

### 3.2 Trial-weighted Bayesian GLMM reveals frequency-general suppression of multisensory integration

To formally characterise this effect and estimate whether tACS selectively influenced the incongruent trial responses rather than inducing a generalised perceptual deficit, the same trial-weighted data were modeled using, a Bayesian Ordered Beta Regression model evaluated the interaction between Stimulation Condition (baseline and different tACS offset frequencies) and Stimulus Type (both congruent and incongruent AV stimuli). The model was specified with wo_tACS (baseline stimulation) and the congruent /ka/ stimulus serving as the baseline intercepts. For the incongruent McGurk stimulus, the illusory (Mg_ta) responses recorded by participants were analysed.

The model revealed evidence for the main effects of both Stimulation Condition and Stimulus Type. Importantly, the analysis yielded high evidence for an interaction between stimulation condition and the processing of the illusory McGurk stimulus (Full parameter estimates, 95% HDI/ROPE decisions, and BF values for all the main effects and interactions are reported in Tables S9-S11, respectively; also see Figure S6 for the HDI/ROPE plots). Specifically, high evidence was found for the interaction between IAF stimulation and the McGurk stimulus log_10_(BF_10_) = 12.648, as well as between IAF-2 and the McGurk stimulus log_10_(BF_10_) = 10.948. The interaction for the IAF+2 condition and the McGurk stimulus also demonstrated high evidence log_10_(BF_10_) = 17.073, see **Figure 7(a)**. Together, these three interactions indicate that each active stimulation frequency selectively modulated the illusory McGurk percept, rather than producing a uniform shift in responses across all stimulus types, supporting a frequency-specific effect of tACS on multisensory integration.

**Figure 7:**
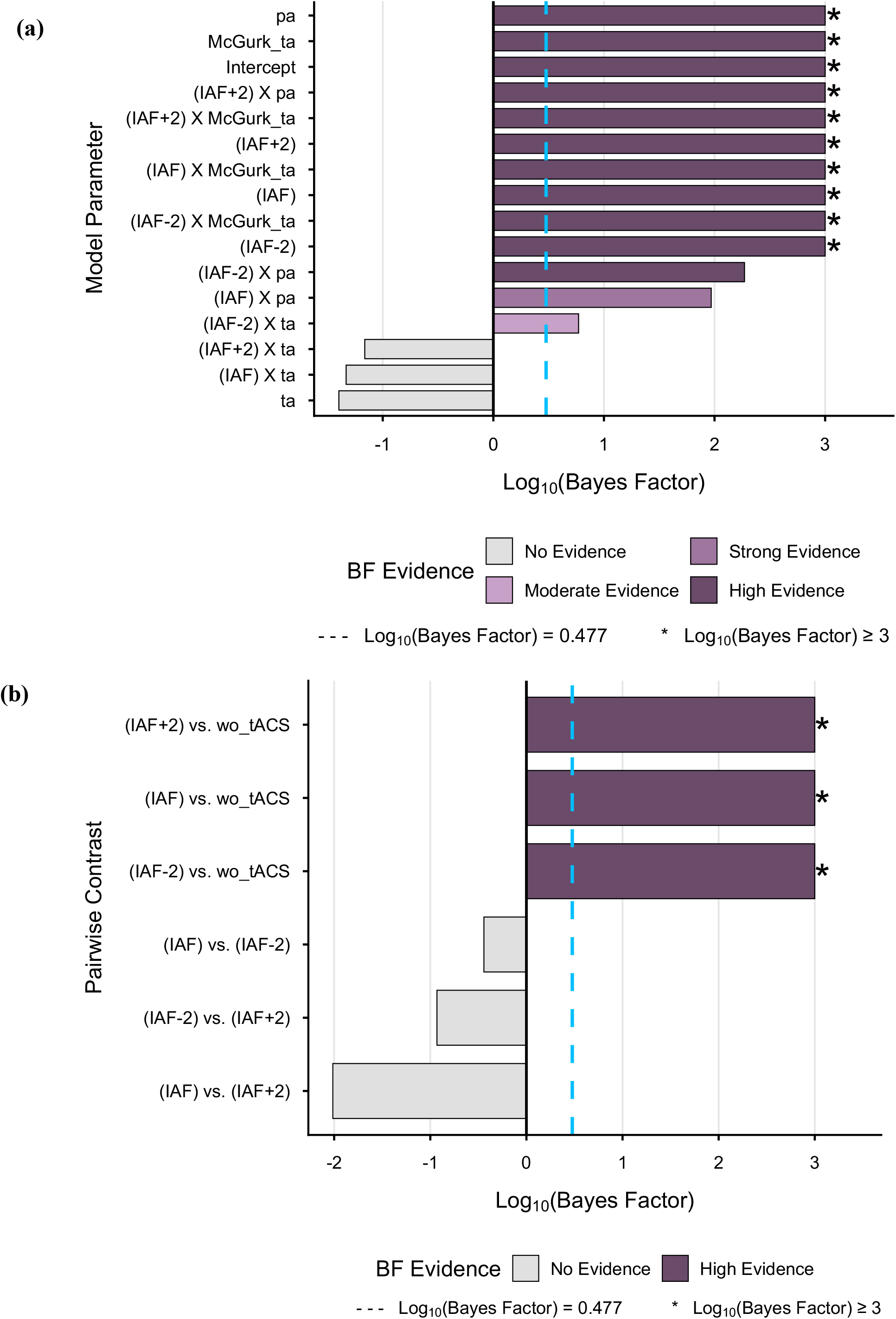
Bayesian strength of evidence for fixed-effect model parameters and stimulus-specific pairwise comparisons. (a) Horizontal bar chart displaying the Log_10_-transformed Bayes Factors (BF_10_) for the conditional main effects and interactions derived from the Generalised Linear Mixed Model (GLMM). The analysis yields high evidence for the main effects of stimulation and stimulus type, along with their crucial interaction during the processing of the incongruent McGurk stimulus. (b) Log_10_-transformed Bayes Factors for post-hoc pairwise comparisons isolating the illusory McGurk response across the various stimulation parameters. The plot illustrates high evidence that all active transcranial alternating current stimulation (tACS) conditions (IAF, IAF-2, and IAF+2) significantly deviated from the no-stimulation baseline (wo_tACS). Conversely, the Bayes Factors for comparisons directly contrasting the active tACS frequencies do not corroborate a difference, even though HDI- and p-value-based criteria indicated credible/significant differences for two of the three comparisons (IAF vs. IAF-2; IAF-2 vs. IAF+2) a divergence between inferential criteria discussed further in Section 4.

Post-hoc hypothesis testing (via *hypothesis()*) isolating the illusory McGurk response revealed that all active tACS conditions deviated from the baseline, which we observed as an reduction in the log-odds of perceiving the illusion across participants. tACS stimulation at IAF resulted in a significant reduction in illusory perception compared to baseline with high evidence (Estimate = -0.341, 95% CI [-0.377, -0.306], estimated error = 0.018, log_10_(BF_10_) = 16.048, p < 0.001). A significant deviation with high evidence was similarly observed for IAF-2 (Estimate = -0.261, 95% CI [-0.316, -0.206], estimated error = 0.028, log_10_(BF_10_) = 138.626, p < 0.001). Stimulation at IAF+2 also caused a significant reduction (Estimate = -0.337, 95% CI [-0.390, -0.283], estimated error = 0.027, log_10_(BF_10_) > 192.7, p < 0.001). Complete post-hoc contrasts for all pairwise comparisons among stimulation conditions are provided in Table S12, Supplementary Materials 3.

Furthermore, comparisons between IAF and the offset frequencies for the McGurk stimulus revealed modest, frequency-dependent asymmetry in the magnitude of multisensory modulation (Figure 7(b)), though the strength of support for this asymmetry differed across our three pre-specified inferential criteria-the p values, HDI and Bayes factor-based conclusions (interpreted further in Section 4). The effect of IAF stimulation differed nominally from the slower IAF-2 condition, with the 95% HDI excluding zero and the FDR-corrected p-value remaining significant (Estimate = -0.08, 95% CI [-0.137, -0.022], estimated error = 0.029, p(adj) = 0.009) however, the corresponding Bayes Factor did not corroborate this effect (log_10_(BF_10_) = -0.442, favouring the null). In contrast, comparing IAF to the faster IAF+2 condition yielded an estimate centered at zero (Estimate = -0.004, 95% CI [-0.059, 0.051], estimated error = 0.028, p(adj) = 0.884, log_10_(BF_10_) = -2.014), with the HDI, p-value, and Bayes Factor in full agreement that no difference exists. A comparison between the offset (IAF-2 vs IAF+2) conditions showed a similar pattern to the IAF vs IAF-2 contrast: a nominal, HDI and significant p(adj) supported difference between IAF-2 and IAF+2 (Estimate = 0.076, 95% CI [0.006, 0.147], estimated error = 0.036, p(adj) = 0.039), again without corroboration from the corresponding Bayes Factor (log_10_(BF_10_) = -0.931).Finally, an examination of the model’s interaction parameters confirmed that the modulatory effects of tACS on the McGurk illusion diverged significantly from baseline congruent perception. As the model utilised the congruent /ka/ stimulus as the comparative baseline, the strong effect associated with the condition-by-stimulus interaction terms decisive by Bayesian criteria for all three tACS conditions (IAF: Estimate = -0.511, log_10_(BF_10_) = 12.648; IAF-2: Estimate = -0.713, log_10_(BF_10_) = 10.948; IAF+2: Estimate = -0.962, log_10_(BF_10_) = 17.073) demonstrate that the tACS-induced disruption of the illusory percept is statistically distinct from the stimulation’s minimal impact on the congruent /ka/ stimulus (see Figure S7, Supplementary Material 2). This was directly confirmed by interaction contrasts testing whether the tACS-related shift in McGurk_ta responses differed from the corresponding shift in ka responses: the difference was credible for IAF (Estimate = 0.510, 95% HPD [0.450, 0.571], p < .001, log_10_(BF_10_) = 14.882), IAF-2 (Estimate = 0.714, 95% HPD [0.600, 0.825], p < .001, log_10_(BF_10_) > 192.7), and IAF+2 (Estimate = 0.963, 95% HPD [0.837, 1.082], p < .001, log_10_(BF_10_) = 17.326) relative to baseline stimulation (See Table S13, Supplementary Material 3 for all remaining stimulus pairs and Figure S8, Supplementary Material 2 for the corresponding plot). These results are consistent with a selective effect on incongruent trial responses relative to congruent speech (see Section 5-Limitations for a discussion of alternative accounts and design-related caveats).

### 3.3 A simple three-node interacting Kuramoto oscillator model explains the pattern of participants’ McGurk /ta/ responses

A three-node Kuramoto model comprising STG, occipital, and STS was employed to simulate how participant-specific synchronisation or McGurk /ta/ responses change as stimulation frequency varies. This three-node model is perturbed by applying tACS to the occipital region of the brain. Synchronisation is measured by the order parameter, which ranges from 0 to 1 and can be related to the proportion of McGurk /ta/ responses for each participant. We have simulated eq. (18) to check the variation of the order parameter across the saddle node curve, and participants 1 and 4 are randomly chosen out of 27 participants. Simulation results are presented in Figure 8, where three different combinations of M and conduction delay have been considered for eq. (18) for both participant 1 and participant 4. From Figures 8(a) and 8(b), we see that the order parameter (R) is close to zero when the tACS frequency is near the IAF for both participants. When tACS frequency is (IAF+2) i.e. Ω ≈ 12.57 *rad*/*sec* and when tACS frequency is (IAF-2) i.e. Ω ≈ −12.57 *rad*/*sec*, the value of the order parameter increases from zero. For participant 1 in Figure 8(a), with M=10 and *τ* = 5 *ms*, the R value at Ω ≈ 12.57 *rad*/*sec* is 0.7582, which is higher than the corresponding R value of 0.6868 at Ω ≈ −12.57 *rad*/*sec*. With M=20 and *τ* = 2.5 *ms*, the R value in participant 1 at Ω ≈ 12.57 *rad*/*sec* is 0.3881, which is higher than the corresponding R value of 0.3076 at Ω ≈ −12.57 *rad*/*sec*. With M=5 and *τ* = 10 *ms*, the R value in participant 1 at Ω ≈ 12.57 *rad*/*sec* is 0.9484, which is higher than the corresponding R value of 0.9237 at Ω ≈ −12.57 *rad*/*sec*. Similarly, for participant 4 in Figure 8(b), with M=10 and *τ* = 5 *ms*, the R value at Ω ≈ 12.57 *rad*/*sec* is 0.7955, which is higher than the corresponding R value of 0.7503 at Ω ≈ −12.57 *rad*/*sec*. With M=20 and *τ* = 2.5 *ms*, the R value in participant 4 at Ω ≈ 12.57 *rad*/*sec* is 0.4393, which is higher than the corresponding R value of 0.3774 at Ω ≈ −12.57 *rad*/*sec*. With M=5 and *τ* = 10 *ms*, the R value in participant 4 at Ω ≈ 12.57 *rad*/*sec* is 0.9596, which is higher than the corresponding R value of 0.9461 at Ω ≈ −12.57 *rad*/*sec*.

**Figure 8:**
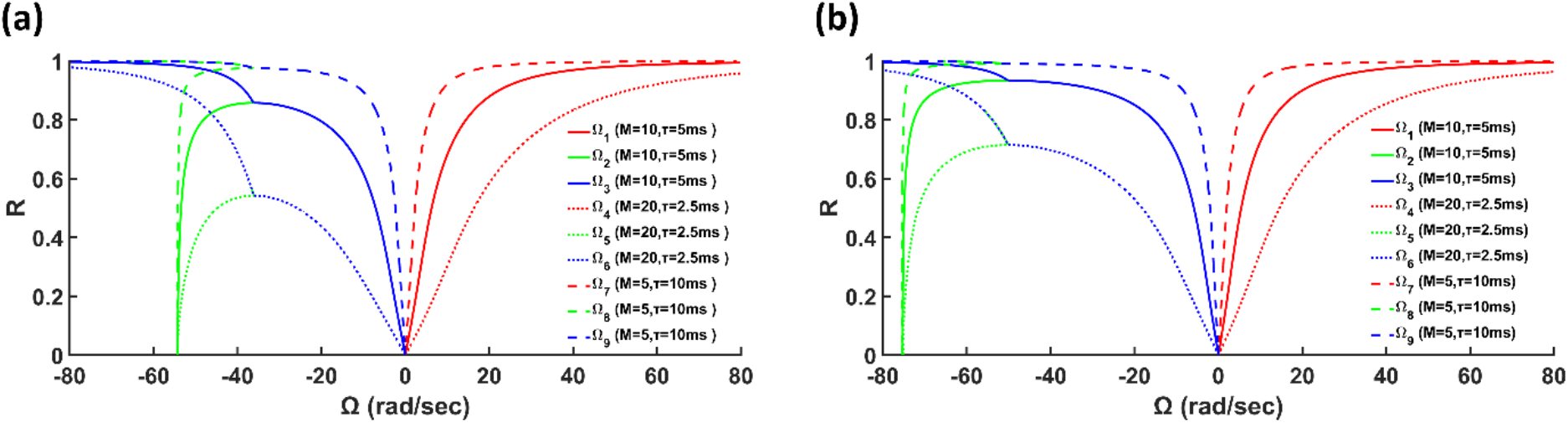
(a) Effect of varying Ω on the order parameter or illusory responses for participant 1 with IAF= 8.65 *Hz*. Red, green and blue colours represent three different branches of the saddle-node curve. Solid line indicates the variation of order parameter for M= 10, *τ* = 5*ms*, dotted line indicates the variation of order parameter for *M* = 20, *τ* = 2.5*ms*, dashed line indicates the variation of order parameter for *M* = 5, *τ* = 10*ms*. (b) Effect of varying Ω on the order parameter or illusory responses for participant 4 with IAF= 12 *Hz*. Red, green and blue colours represent three different branches of the saddle-node curve. Solid line indicates the variation of order parameter for M= 10, *τ* = 5*ms*, dotted line indicates the variation of order parameter for *M* = 20, *τ* = 2.5*ms*, dashed line indicates the variation of order parameter for *M* = 5, *τ* = 10*ms*.

The order parameter plots along the homoclinic trajectory are also presented for participants 1 and 4 under different coupling strengths and time delays. We have simulated equation (20) in (Ω,t) plane for 7 minutes since the experiment with tACS was conducted for 7 minutes for each participant. It can be observed from Figure 9(a) and Figure 9(b) that the variation of order parameter along the homoclinic trajectory does not affect Ω = 0, Ω ≈ 12.57 *rad*/*sec and* Ω ≈ −12.57 *rad*/*sec* for the participants. In Figure 10(a), we have simulated eq. (17) for Hopf curve, eq. (18) for three branches of saddle-node curve and eq. (20) for homoclinic manifold for participant 1 at *M* = 10 *and τ* = 5 *ms*. Similarly, in Figure 10(b), we have simulated eq. (17) for Hopf curve, eq. (18) for three branches of saddle-node curve and eq. (20) for homoclinic manifold for participant 4 at *M* = 20 *and τ* = 2.5 *ms*.

**Figure 9:**
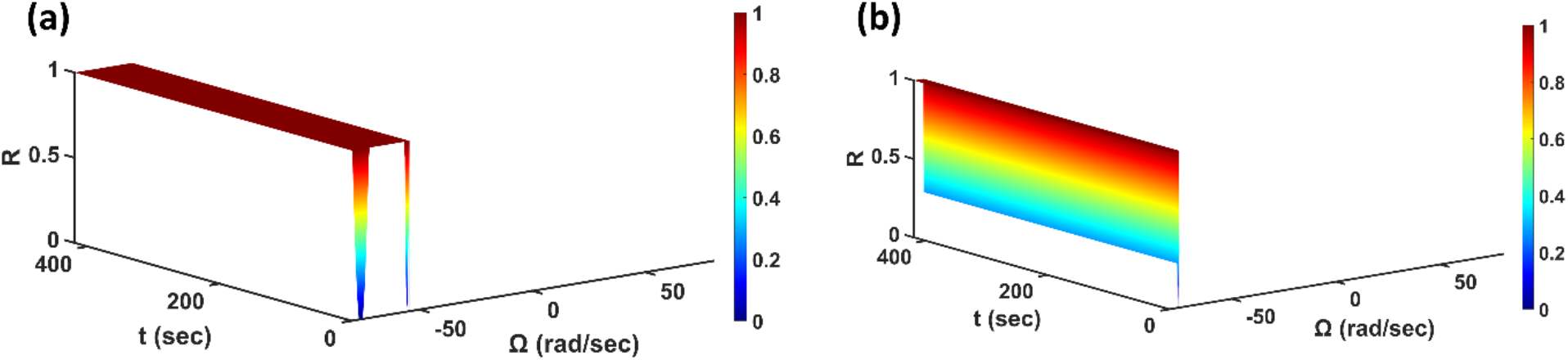
(a) 3D surface plot illustrating the variation of the order parameter or illusory responses along the homoclinic trajectory in the (Ω, *t*) plane for participant 1 with IAF= 8.65 *Hz for M* = 10 *and τ* = 5 *ms*. Colourbar indicates different values of the order parameter. Variation of the order parameter does not affect Ω = 0, Ω ≈ 12.57 *rad*/*sec and* Ω ≈ −12.57 *rad*/*sec*. (b) 3D surface plot illustrating the variation of the order parameter or illusory responses along the homoclinic trajectory in the (Ω, *t*) plane for participant 4 with IAF= 12 *Hz for M* = 20 *and τ* = 2.5 *ms*. Colourbar indicates different values of the order parameter. Variation of the order parameter does not affect Ω = 0, Ω ≈ 12.57 *rad*/*sec and* Ω ≈ −12.57 *rad*/*sec*.

**Figure 10:**
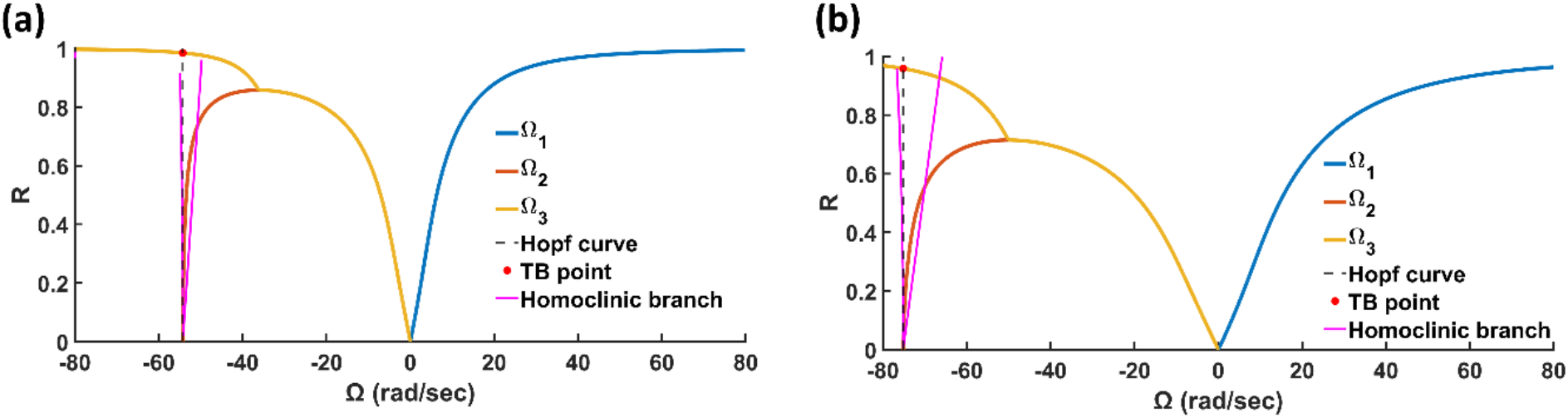
(a) Effect of varying Ω on the order parameter or illusory responses for participant 1 with IAF= 8.65 *Hz for M* = 10 *and τ* = 5 *ms*. The red, yellow and blue solid lines represent three different branches of the saddle-node curve. Dashed line indicates Hopf curve for participant 1 which intersects top yellow saddle-node branch at (Ω_*TB*_, *R*_*TB*_) ≈ (−54.239, 0.985). Pink solid line indicates the homoclinic trajectory of order parameter or illusory responses for participant 1. For homoclinic trajectory, (20) has been simulated for participant 1 for *t* = 3 minutes. (b) Effect of varying Ω on the order parameter or illusory responses for participant 4 with IAF= 12 *Hz for M* = 20 *and τ* = 2.5 *ms*. The red, yellow and blue solid lines represent three different branches of the saddle-node curve. Dashed line indicates Hopf curve for participant 4 which intersects top yellow saddle-node branch at (Ω_*TB*_, *R*_*TB*_) ≈ (−75.178, 0.958). Pink solid line indicates the homoclinic trajectory of order parameter or illusory responses for participant 4. For homoclinic trajectory, eq. (20) has been simulated for participant 4 for *t* = 3 minutes.

Ten participants’ percentage McGurk /ta/ responses were higher at (IAF-2) than at (IAF+2) (see Table 4 in Supplementary Material 4). Out of these ten participants, participants 1 and 6 have been randomly chosen, and we have invoked the effective coupling strategy to simulate the locus of the saddle-node bifurcation. From Figure 11(a) and Figure 11(b), we can see that for both participant 1 and participant 6, the order parameter or percentage of illusory responses at Ω ≈ −12.57 *rad*/*sec* is greater than at Ω ≈ 12.57 *rad*/*sec* as well as the finite value of the order parameter magnitude exists at Ω = 0 (see also Table 3 in Supplementary Material 4) which aligns with the pattern of the empirical observation. However, in earlier case, without effective coupling, order parameter value at Ω ≈ 12.57 *rad*/*sec* is always greater than at Ω ≈ −12.57 *rad*/*sec* as well as the magnitude of the order parameter is undefined at Ω = 0 for 13 out of 17 participants (see also Table 1 in Supplementary Material 4).

**Figure 11:**
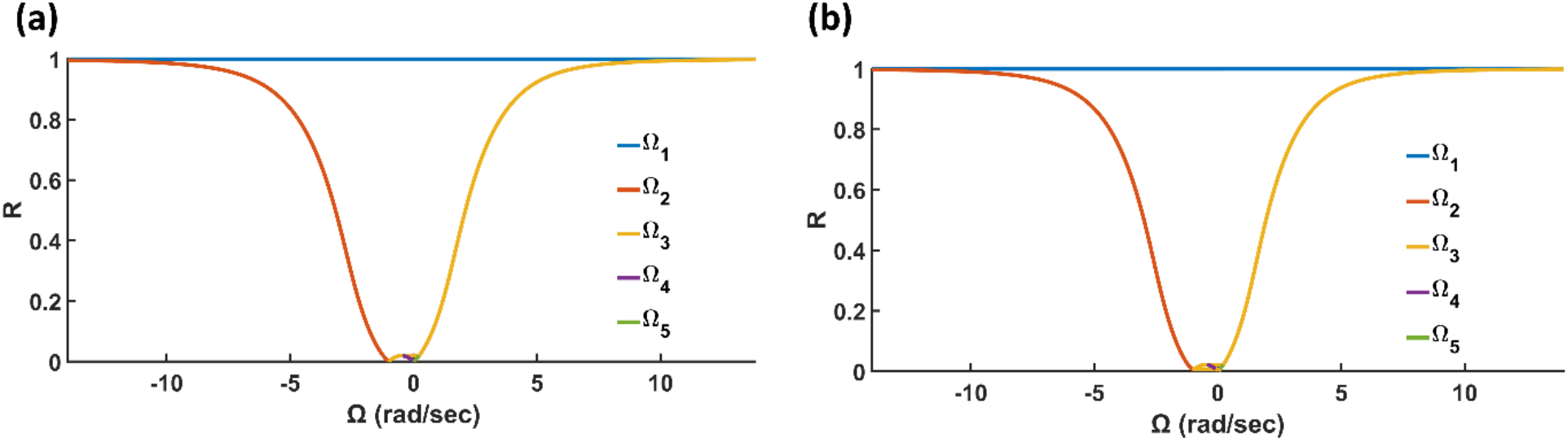
(a) Effect of varying Ω on the order parameter or illusory responses for participant 1 with IAF= 8.65 *Hz for M* = 10 *and τ* = 5 *ms*. The red, yellow, blue, violet and green solid lines represent five different branches of the saddle-node curve with effective coupling for *β* = 1. At Ω ≈ 12.57 *Hz*, R is 0.9974 whereas at Ω ≈ −12.57 *Hz*, R is 0.9995. At Ω = 0, R is 0.0194. Homoclinic trajectory is not shown. (b) Effect of varying Ω on the order parameter or illusory responses for participant 6 with IAF= 11.2 *Hz for M* = 10 *and τ* = 5 *ms*. The red, yellow, blue, violet and green solid lines represent five different branches of the saddle-node curve with effective coupling for *β* = 1. At Ω ≈ 12.57 *Hz*, R is 0.9980 whereas at Ω ≈ −12.57 *Hz*, R is 0.9997. At Ω = 0, R is 0.0219. Homoclinic trajectory is not shown.

Finally, we correlated model-predicted order-parameter asymmetry (R values at IAF-2 minus IAF+2) vs. behavioural asymmetry (the proportion of McGurk /ta/ responses at IAF−2 minus IAF+2) across 27 participants. We utilised interpolation/extrapolation techniques (see the details in Supplementary Material 4) to obtain plausible values for participant-specific M and *τ* (see Table 1 in Supplementary Material 4). These participant-specific M and *τ* values are put in the equation (18) to get R values at (IAF-2) and (IAF+2). For 10 participants whose % /ta/ responses at (IAF-2) are greater than at (IAF+2), we put eq. (21) into eq. (18) to obtain the locus of the quintic saddle-node curve. Instead of using the interpolation/extrapolation technique, we have kept the same M, *τ*, and *β* values for these 10 participants since variation of M, *τ*, and *β* values provide almost similar R values at (IAF-2), IAF, and (IAF+2) obtained from this quintic saddle-node curve. Without tACS, the case is excluded from the correlation analysis (see the details in Supplementary Material 4). The regression line and scatter plot for all 27 participants are depicted in Figure 12. Though directionality is present (for 17 participants, % /ta/ responses at (IAF+2) are higher than at (IAF-2), for 10 participants it is reverse), Figure 12 indicates a statistically significant, positive Spearman correlation (*ρ* = 0.8061, *p* ≪ 0.00001) between model-predicted order-parameter asymmetry (R values at IAF-2 minus IAF+2) and behavioural asymmetry (the proportion of McGurk /ta/ responses at IAF−2 minus IAF+2).

**Figure 12:**
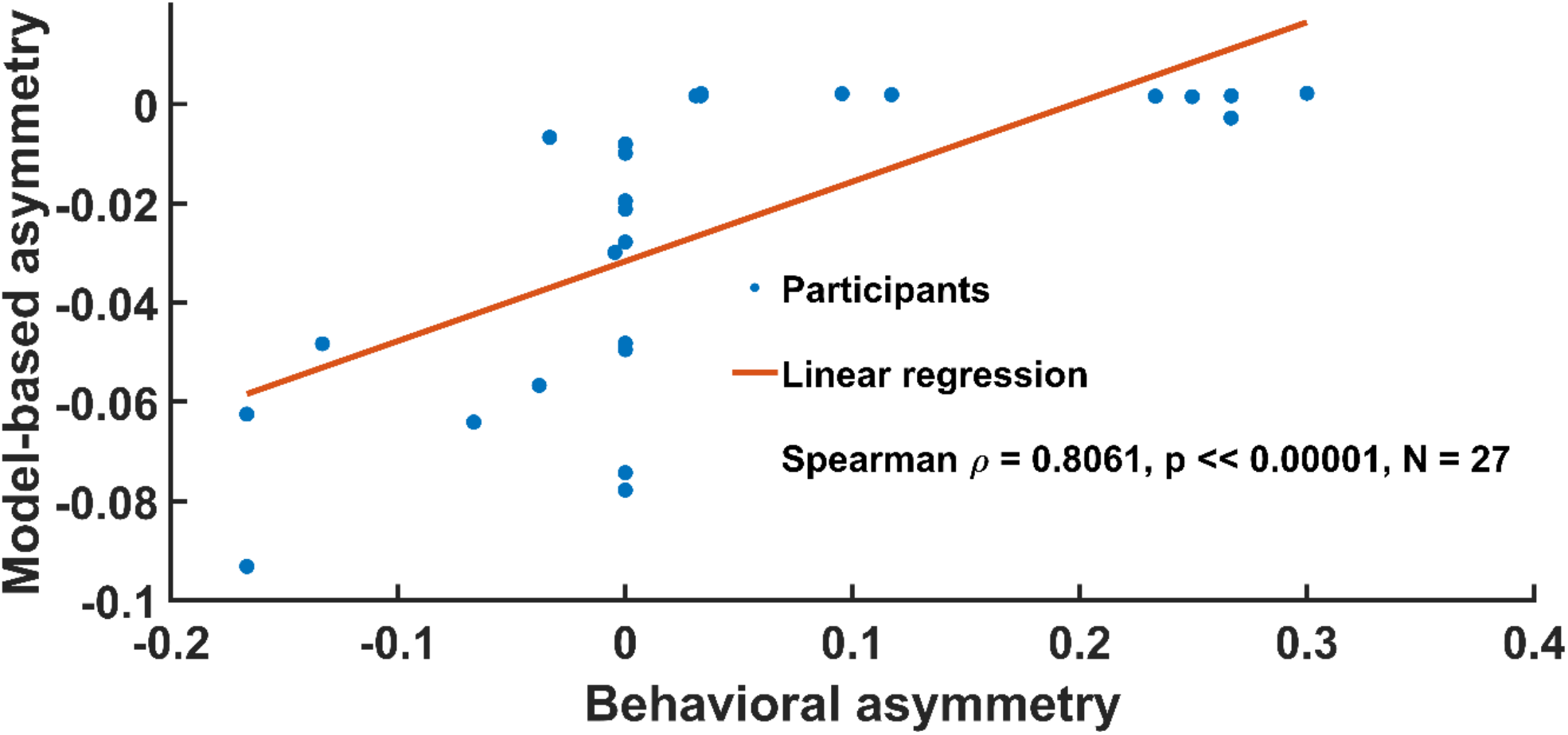
The regression line and scatter plot between Kuramoto model-predicted order-parameter asymmetry (R values at IAF-2 minus IAF+2) vs. behavioural asymmetry (the proportion of McGurk /ta/ responses at IAF−2 minus IAF+2) across all the 27 participants (see Table 5 in supplementary material 4 for these correlations and regression statistics). Blue circles and red lines indicate the participants and the linear fit, respectively (ρ=0.8061, p≪0.00001, *N* = 27).

## 4 Discussion

The present study investigated the causal role of occipital alpha-band oscillations in multisensory speech processing by applying individualised transcranial alternating current stimulation (tACS). A central challenge in multisensory speech perception processing is maintaining the balance between sensory integration (binding congruent, synchronous cues) and sensory segregation (parsing conflicting or asynchronous cues into distinct perceptual events) (Rosenblum, 2019). A substantial body of work now converges on the idea that the speed of alpha oscillations, sets the grain of this temporal parsing. Individuals with faster IAF tend to show finer temporal resolution and more accurate segregation of closely spaced events, while slower IAF is associated with a bias toward integration (Samaha & Postle, 2015; reviewed in Samaha & Romei, 2024), although the relationship has not always replicated (Buergers & Noppeney, 2022; London et al., 2022). Causal manipulation work has reinforced this. Entraining visual cortex (on V5) at 10 Hz via tACS shifts observers toward integrating two successive flashes into one percept relative to sham stimulation (Battaglini et al., 2020), and more recent bifocal tACS work targeting extra-striate visual areas found that stimulating above IAF (IAF+2) improved temporal segregation accuracy relative to stimulating below IAF (IAF-2), broadly consistent with a “faster alpha, finer resolution” account (Santoni et al., 2025). This general relationship between alpha speed and temporal/perceptual precision has since been replicated using a range of paradigms and populations (Mierau et al., 2017; Wutz et al., 2018; Cooke et al., 2019; Migliorati et al., 2020; Noguchi, 2022; Ronconi et al., 2022), and extends specifically to the temporal window over which multisensory cues are bound. Cecere and others (2015) showed that individual differences in resting IAF predict the width of the audiovisual temporal binding window in a simple flash-beep illusion, and Venskus and Hughes (2021)further dissociated this relationship from time perception more broadly, tying IAF specifically to the multisensory integration window. While causal evidence links intrinsic alpha frequency to the temporal window of multisensory binding in simple visual and audiovisual illusions (Cecere et al., 2015), and correlational work extends this relationship to the temporal resolution of perception (Samaha & Postle, 2015), however, direct causal evidence within the more complex, ecologically valid domain of audiovisual speech has remained limited. By applying tACS stimulation at each participant’s occipital IAF and offsets (IAF2, IAF-2), we observed a significant reduction in propensity to McGurk perception (see Figure 7(b)). This effect suggests that exogenous modulation of occipital alpha reduces susceptibility to the McGurk illusion. Consistent with this finding, an examination of the model’s interaction parameters showed that the effect of tACS on the incongruent McGurk illusion was statistically distinct from its effect on congruent speech processing. The significant reduction in illusory perception across IAF, IAF-2 and IAF+2 is not readily explained by a mechanism that simply degrades visual-stream fidelity, since such a mechanism should scale primarily with current dose rather than with the specific relation between stimulation frequency and endogenous alpha. Occipital tACS selectively reduced responses to the audio visually incongruent McGurk stimulus while leaving congruent speech largely unaffected. Two accounts could explain this selectivity. Alpha activity is proposed to gate information flow through pulsed inhibition and to time cortical activity (Jensen & Mazaheri, 2010; Klimesch, 2012) and ongoing oscillatory state and STS connectivity influence the occurrence of McGurk fusion (Keil et al., 2012). Another possibility is that tACS reduced the weight given to the visual stream, shifting responses toward the auditory percept, since STS connectivity with sensory regions varies with cue reliability (Nath & Beauchamp, 2011). The present design cannot separate these accounts, and testing them is a direction for future work.

**Table 5.** (Correlation and Regression statistics)

|  |  |
| --- | --- |
| Pearson $r$ | 0.6741 |
| 95% CI | [0.3953, 0.8391] |
| p-value | 0.000116 |
| Spearman $\rho$ | 0.8061 |
| 95% bootstrap CI | [0.6154, 0.8992] |
| Spearman p-value | 0.000000 |
| Slope | 0.1606 |
| Intercept | -0.0317 |
| Regression $R^2$ | 0.4544 |

The primary finding of this study is the robust, frequency-general reduction of the McGurk effect across all active tACS conditions relative to baseline (see, Figure 7(b)), linking alpha oscillatory activity to the temporal window of integration (TWI). Prior work also links occipital alpha frequency to temporal integration windows. Individual alpha frequency (IAF) is negatively associated with the width of the audiovisual temporal binding window (Cecere et al., 2015; Venskus & Hughes, 2021) and with the temporal resolution of visual perception (Samaha & Postle, 2015). This reduction was of comparable magnitude across IAF, IAF-2, and IAF+2: pairwise comparisons between these three active frequencies yielded 95% credible intervals excluding zero (except for comparison between IAF and IAF+2), accompanied by FDR-corrected p-values reaching conventional significance (Figure 7b), together nominally consistent with a small offset-dependent asymmetry but the corresponding Bayes Factors did not corroborate either effect (log_10_(BF_10_) = -0.442 (for IAF vs IAF-2) and -0.931 (for IAF-2 vs IAF+2), respectively), instead anecdotal-to-moderately favouring the null (Figure 7b). This divergence is expected within a single Bayesian model, because interval-based and Bayes-factor-based conclusions answer different questions, and the Bayes factor also depends on the prior width for small effects (Kruschke, 2011). Given this, we treat the offset asymmetry as an open question rather than a confirmed effect, a caveat that concerns only the fine-grained comparison between active frequencies, and does not bear on the primary, decisively supported result that sall three active frequencies robustly reduced illusory perception relative to baseline. Under either reading, the overall pattern does not match the strict pacemaker framework (Samaha & Postle, 2015), in which predicts a bidirectional effect relative to baseline.A faster rhythm (IAF+2) should narrow the temporal window of integration and reduce illusion susceptibility, while a slower rhythm (IAF-2) should widen the window and increase it. We did not observe this. All three active frequencies reduced the illusion relative to baseline, with IAF−2 showing a slightly smaller reduction than IAF+2. The direction of this small asymmetry is consistent with the faster-alpha, finer-resolution account, and with the IAF+2 > IAF−2 ordering Santoni et al., 2025 reported for two-flash segregation., although, as noted above, it was not corroborated by the Bayes factors. Importantly, such an ordering is a difference between offsets, whereas a pacemaker account also predicts a change of sign relative to baseline, which we did not find. The dominant feature of our data is therefore the broadly similar reduction across all three active frequencies. This suggests that exogenous entrainment may disrupt the McGurk illusion by overriding the natural pacing of endogenous alpha, rather than through frequency-specific widening or narrowing of the temporal window alone.It further indicates that the mechanism engaged by occipital tACS in the McGurk paradigm may differ, at least in degree, from that operating in simple flash-fusion illusions, where bidirectional (Cecere et al., 2015), frequency-specific effects have been reported (Battaglini et al., 2020).

These behavioural findings establish that alpha entrainment reliably shapes multisensory binding, with a robust reduction in illusory perception observed across all three active stimulation frequencies relative to baseline. At the population level, this reduction was significantly smaller at IAF-2 than at IAF and IAF+2 on the 95% HDI and the FDR-corrected, posterior-derived p-value, though this asymmetry was not corroborated by the corresponding Bayes Factors (Section 3.1). Considerable heterogeneity was apparent at the individual-participant level, however, with the majority of participants showing a numerically larger reduction at IAF+2 than at IAF-2, and a substantial minority (10 of 27) showing the reverse pattern (See Figure S3, Supplementary Material 2). To characterize the synchronisation dynamics that could give rise to this frequency-general suppression of the illusion, and to the participant-specific heterogeneity observed within it, we turn to a Kuramoto model of coupled neural oscillators. Within this Kuramoto framework, the order parameter provides a phenomenological measure of the degree of phase synchronisation induced by tACS and therefore offers a simple mechanistic link between stimulation frequency and multisensory perceptual outcome. In the context of the occipital-STG-STS network, tACS-induced modulation of alpha activity in the occipital region can alter the temporal coordination between occipital and STG regions which are subsequently integrated within the STS region. Hence, shifts in the model’s synchronisation can be interpreted as a change in the timing of sensory signals reaching the STS region. This shift can alter how likely a person is to experience the McGurk illusion (Keil et al., 2012; Nath & Beauchamp, 2012; Kumar et al., 2020).

Furthermore, the underlying nonlinear dynamics further indicate that the response to stimulation is not necessarily a simple function of the difference between tACS frequency and IAF. The saddle node and Hopf structures of the parsimonious model define distinct synchronisation regimes that provide a possible explanation for the strong dependence of the order parameter on stimulation frequency. Near the IAF, synchronisation can be reduced when dynamics approach a weakly synchronised regime, whereas stimulation away from this region can move the system towards stronger phase locking. This provides a mechanistic interpretation of the predominant pattern, in which (IAF+2) produces stronger synchronisation than (IAF-2) across most participants. However, these synchronisation-desynchronisation phenomena were explained by changing the coupling strength in the finite-dimensional Kuramoto model (Maistrenko et al., 2004; Maistrenko et al., 2005).

Importantly, however, the present Kuramoto formulation assumes identical oscillators and ignores heterogeneity in intrinsic frequencies or conduction delays. As a consequence, this model alone cannot explain why some participants show a reverse pattern, where synchronisation at (IAF-2) exceeds that at (IAF+2) or remains finite near IAF. Although the dynamical properties of the model suggest that alternative dynamical regimes such as homoclinic transition may contribute to such variability, these possibilities should be regarded as hypotheses rather than direct mechanistic predictions of the present model. More detailed models incorporating heterogeneous intrinsic frequencies or delays (Lee et al., 2009; Skardal, 2018) between occipital, STG, and STS regions will be required to determine how individual differences in network dynamics give rise to these distinct tACS response profiles. However, this issue has been addressed here by imposing an effective coupling criterion within the Kuramoto model framework with identical oscillators.

## 6 Limitations

Two considerations are important when interpreting the baseline-vs-active tACS comparisons reported above. First, the two offset conditions (IAF+2, IAF−2) served as frequency-specific active controls rather than a sham condition: because real current was delivered in all three active conditions, nonspecific sensory and expectation confounds (skin sensation, phosphenes, arousal, awareness of being stimulated) that a true sham could not fully rule out were held constant, allowing frequency-specificity to be assessed cleanly among IAF, IAF+2, and IAF−2. Second, however, the wo_tACS baseline was not counterbalanced against the active conditions: for logistical reasons, IAF had to be measured before the Day 2 stimulation frequencies could be set, so Day 1 always consisted of the no-stimulation session and Day 2 always consisted of the three active conditions. Because baseline was therefore always the first, shorter (40–50 min vs. ∼2.5 hr) exposure to the task, differences between wo_tACS and any active tACS condition – including the reduction in illusory /ta/ percepts reported above – cannot be fully dissociated from generic practice, fatigue, or session-order effects. This confound does not affect the frequency-specificity comparisons among the three active conditions, which were all obtained within the same Day 2 session and order structure. A limitation of the present design is the restricted range of stimulation frequencies (IAF±2 Hz); broader offsets and independent psychophysical estimates of the temporal binding window in the same participants would be required to distinguish a TBW-based pacemaker mechanism from a frequency-nondirectional entrainment-override mechanism. An alternative interpretation, that occipital tACS selectively degrades visual encoding of the articulator movements and thereby biases responses toward the auditorily-correct /pa/, cannot be fully excluded by the present design; the observed simultaneous decrease in McGurk /ta/ responses and increase in congruent /pa/ accuracy under IAF stimulation is qualitatively consistent with such an account. Relatedly, because analyses were proportion-based without explicit criterion modelling and response mappings were fixed, a response-bias account in which stimulation shifts participants’ criterion for reporting “/pa/” rather than altering the underlying percept cannot be fully excluded (David M. Green, 1967; Mallick et al., 2015). This concern is compounded by the near-ceiling accuracy already observed on congruent /ka/ trials at baseline (Section 3.1), which leaves limited room to detect any comparable bias-driven change on congruent stimuli and makes the apparent selectivity of the tACS effect for incongruent trials difficult to fully dissociate from a general response-bias account. Finally, the current statistical specification aggregates across blocks and includes only participant random intercepts; residual day/block and participant-specific slope effects may therefore partially contribute to the Condition × Stimulus estimates reported above. For 10 participants in whom the % McGurk /ta/ responses at (IAF-2) are higher than at (IAF+2), the R values at IAF are close to zero, whereas R values are close to 1 at both (IAF-2) and (IAF+2) (see Table 4 in Supplementary Material 4; see also Figure 11(a) and Figure 11(b)). This nearly U-shaped pattern of R values indicates that the Kuramoto model behaves closely in robust attractor/saturation mode. To capture participant-specific variability in McGurk /ta/ responses, rather than tuning M, *τ*, and *β* values, we need to introduce an additional parameter into the existing Kuramoto framework with a plausible, legitimate biological interpretation and then tune it. Even though the Kuramoto model broadly reproduced the predominant behavioural pattern, stimulation at (IAF+2) produced stronger synchronisation than stimulation at (IAF-2) in the majority of participants (17/27). Because this match rate did not reach statistical significance (p = 0.12), it indicates qualitative alignment rather than a statistically validated prediction.

## 6 Conclusions and future directions

Despite these limitations this study provides causal evidence that the frequency and synchronisation dynamics of occipital alpha oscillations play an important role in shaping multisensory speech perception. By applying individualised tACS, we successfully modulated the rate of illusory cross-modal binding without affecting general unisensory comprehension, highlighting the close coupling between neural oscillatory speed and sensory integration. We further implemented Kuramoto oscillator model that suggested that the individual variability in order parameter asymmetry observed across participants may arise from proximity to a Takens–Bogdanov bifurcation, offering a candidate mechanistic account for why some participants deviate from the expected IAF+2/IAF−2 pattern. While the behavioural and statistical evidence presented here provides direct causal support for the role of occipital alpha oscillations in multisensory speech perception, the specific network-level mechanisms proposed to underlie this effect are derived from computational modelling rather than direct neural measurement. In the Kuramoto framework, we used the tACS frequency as the sole control parameter to study how the system’s behaviour changes. In the future, we shall extend this framework by invoking heterogeneity in intrinsic oscillator frequencies as well as inter-oscillator time delays (Omel’chenko et al., 2008), which will break the underlying symmetry and enable the exploration of substantially richer dynamical behaviours. Also, we shall introduce additional bifurcation parameters, such as coupling strength and conduction delay, into the system that may unfold more complex synchronisation regimes and provide a more accurate explanation of the empirical observations. Moreover, the present model does not account for higher-order interactions that can emerge in finite-size oscillator populations (Clusella & Politi 2020; Costa et al. 2024). Therefore, a natural extension would be to study higher-order synchronisation using the Kuramoto–Daido order parameter that might help us to understand these collective effects to unravel the deeper insight into the experimentally observed phenomena. Since multisensory speech perception involves multiple brain regions rather than the STG-STS-occipital network, we can extend our three-node network model to the virtual brain in the future to gain deeper insights.

Beyond refining the computational model itself, future investigations should also pivot toward leveraging simultaneously acquired EEG datasets to extract precise source-level functional connectivity. Utilising advanced spatial filtering techniques will allow researchers to mathematically quantify how targeted tACS over the occipital cortex modulates phase coherence and directional information flow between the visual cortex and the superior temporal sulcus (STS). Additionally, the development of large-scale computational models of these specific oscillatory networks will be critical. Simulating these temporal dynamics in silico will enable the prediction of individualised responses to tACS, facilitating the optimisation of stimulation parameters (such as current density, exact phase angles, and tailored offset frequencies) and supporting the eventual translation of these findings toward clinical applications.

## Conflict of Interest

The authors declare that the research was conducted in the absence of any commercial or financial relationships that could be construed as a potential conflict of interest.

## Author Contributions

SD: [Conceptualisation, Data curation, Formal analysis, Investigation, Methodology, Software, Visualization, Writing – original draft, Writing – review & editing]. DB: [Conceptualisation, Formal analysis, Methodology – computational modelling, Software, Visualisation, Writing - original draft, Writing – review & editing]. VAVS: [Conceptualisation, Formal analysis, Methodology, Software, Supervision, Writing – review & editing]. VM: [Formal analysis, Supervision, Writing – review & editing]. DR: [Conceptualisation, Funding Acquisition, Project Administration, Supervision, Writing – review & editing]. SD and DB equally contributed to this work. All authors contributed to the article and approved the submitted version.

## Funding

The project was supported by SERB Core Research Grant (CRG) CRG/2022/001049 extramural grant from the Department of Science and Technology, Ministry of Science and Technology, Govt. India.

## Acknowledgments

The authors acknowledge the generous support of IIT Jodhpur core funds and computational facility.

## Data and Code Availability

The data and codes for this study can be found on the Open Science Framework at https://osf.io/ej8wt/overview

## Abbreviations

EEG: electroencephalography
STS: superior temporal sulcus
PPR: percentage perceptual response
MCMC: Markov chain Monte Carlo
GLMM: generalised linear mixed model
BF: Bayes factor.

## Supplementary Material 1 - Derivations

### Derivation of eq. (10)

Let us start with Dean-Kawasaki formulation [1] which leads to Langevin equation for the evolution of oscillators’ density *ρ* that generalizes the Fokker-Planck equation

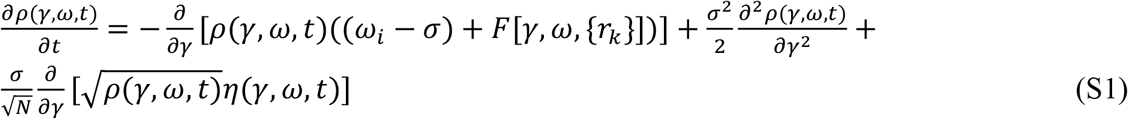

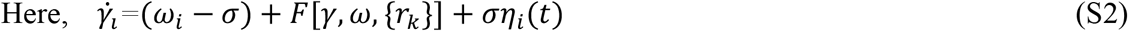

In our Kuramoto model, no stochastic term is there. Hence, *σ* = 0.

We can write, Kuramoto order parameter *r*(*t*) as following

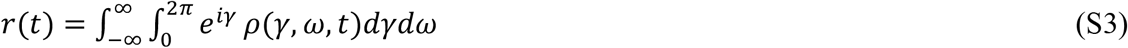

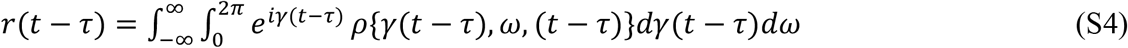

We can write, velocity *v*(*γ, ω, t*) on the circle as following

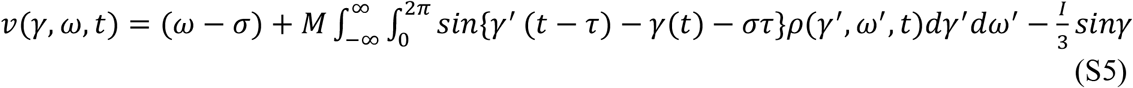

Therefore, from eq. (S1), eq. (S2), eq. (S4), and eq. (S5), we can write

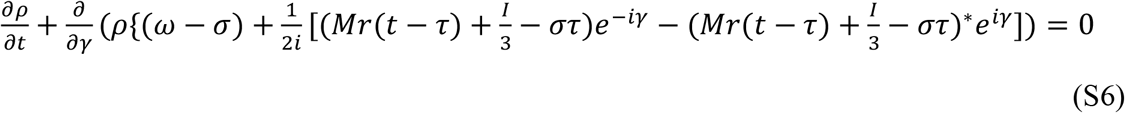

By expanding *ρ* as a Fourier series in *γ*, we can write,

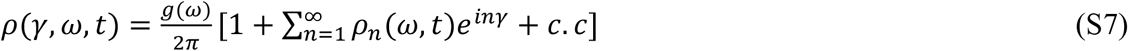

By applying Ott-Antonsen ansatz [2], we can write,

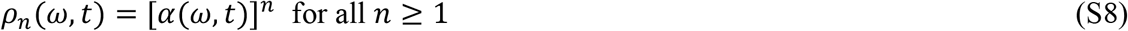

From eq. (S4), eq. (S7), and eq. (S8), we can write,

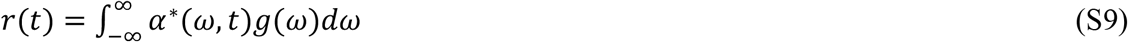

From eq. (S6), eq. (S7), and eq. (S8), we can write,

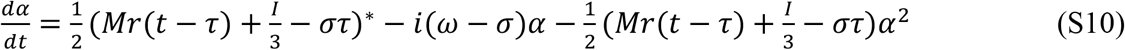

For identical oscillators, *g*(*ω*) = *δ*(*ω* − *ω*_0_)

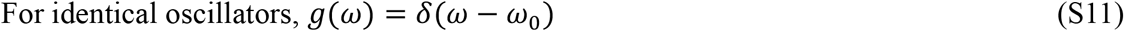

From eq. (S9) and eq. (S11), we can write,

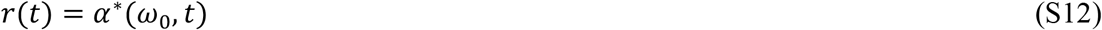

Then, from eq. (S10) and eq. (S12), we can write,

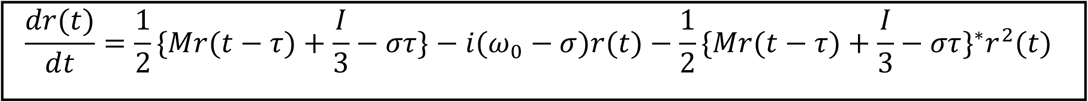

### Derivation of eq. (S19) and eq. (S20)

Since *τ* ≪ 1, we can write, 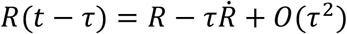

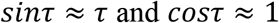

Let, 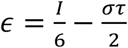

Then, we can write,

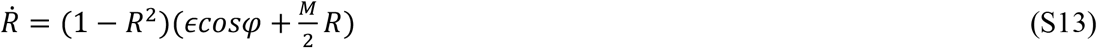

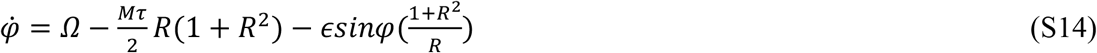

At steady state,

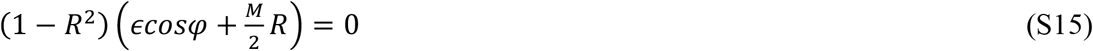

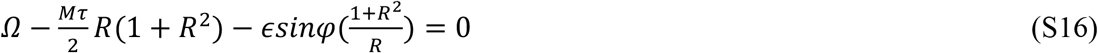

Let us define the perturbations,

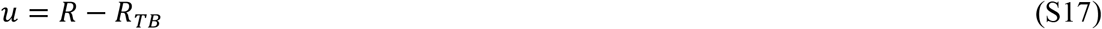

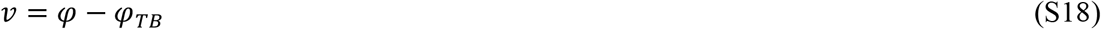

Substituting eq. (S15), eq. (S17), and eq. (S18) in eq. (S13), we can write,

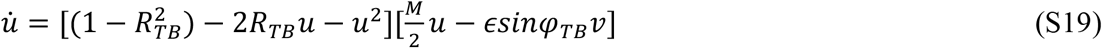

Which is as following

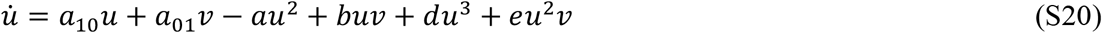

Where 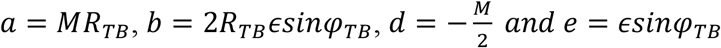

Similarly, substituting eq. (S16), eq. (S17), and eq. (S18) in eq. (S14), we can write,

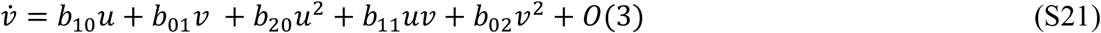

The linearized system is

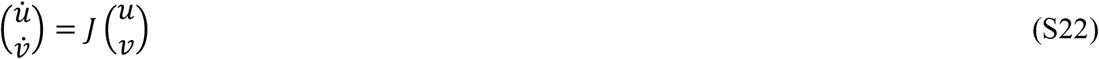

At TB, *tr J* = 0 and *det J* = 0

After linear transformation to Jordan form 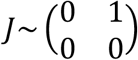 and introducing canonical coordinates

*x* = *u* and 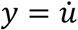, we can write,

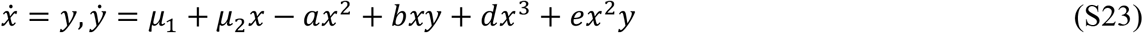

Where *μ*_1_ = Ω − Ω_*TB*_, *μ*_2_ = *σ* − *σ*_*TB*_

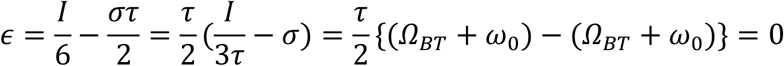

Therefore, *b* = 0 and *e* = 0

Hence, normal form of TB is following

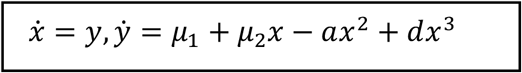

Now, we can write,

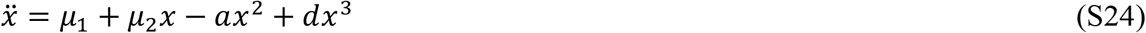

Multiplying by 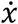 and integrating, we get,

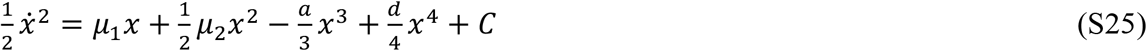

Thus, the Hamiltonian is

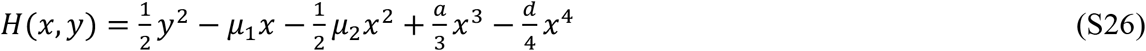

Homoclinic orbit exists when the energy level through the saddle closes.

At equilibria,

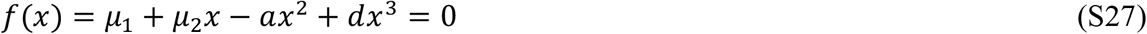

The saddle node curve is obtained from *f*^′^(*x*) = *μ*_2_ − 2*ax* + 3*dx*^2^ = 0

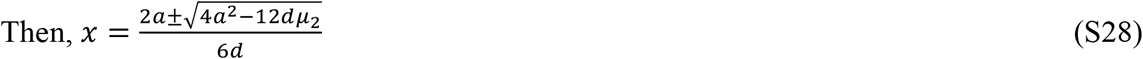

Choosing the branch that passes through *x* = 0 when *μ*_2_ = 0

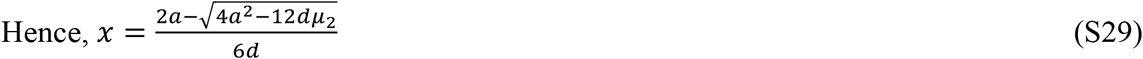

Expanding the square root for small *μ*_2,_

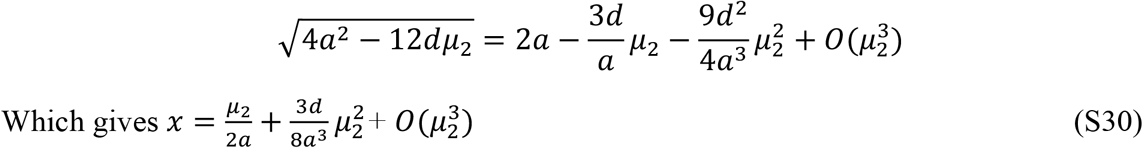

Substituting eq. (S30) into eq. (S27), we get,

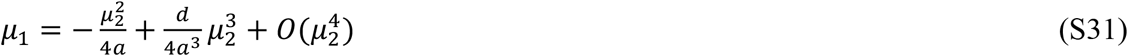

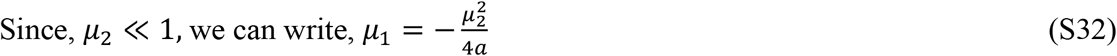

We know that the homoclinic bifurcation curve from TB satisfies the following asymptotic [3]

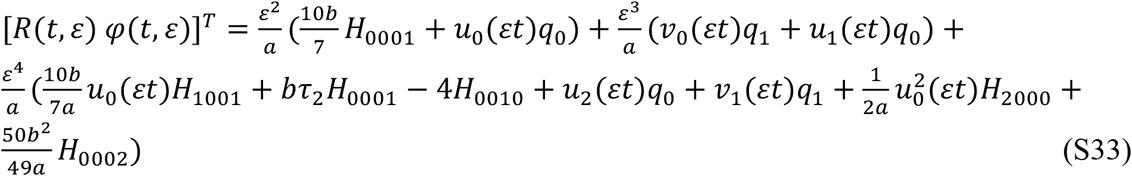

Here, 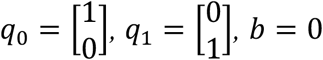

Since *b* = 0, the system falls into degenerate TB and homoclinic orbit will not emerge from TB point.

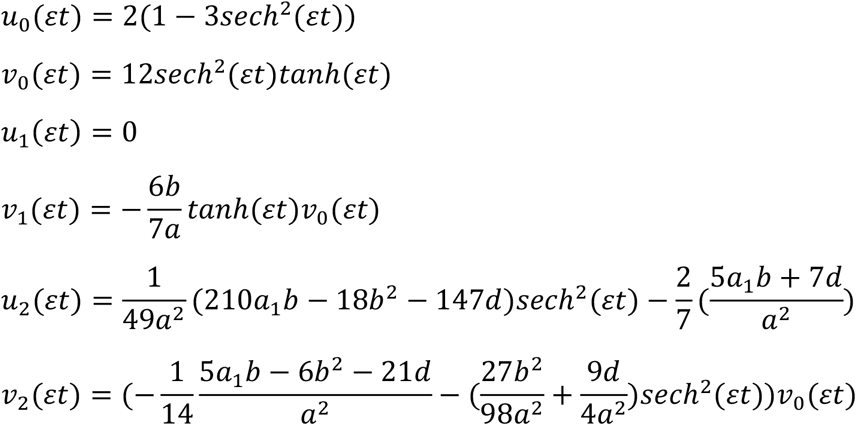

Hence, we can write,

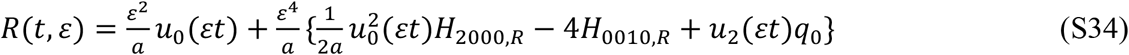

The standard TB scaling is following:

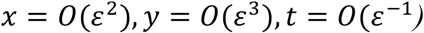

Therefore, we can write,

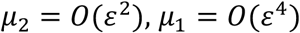

Finally,

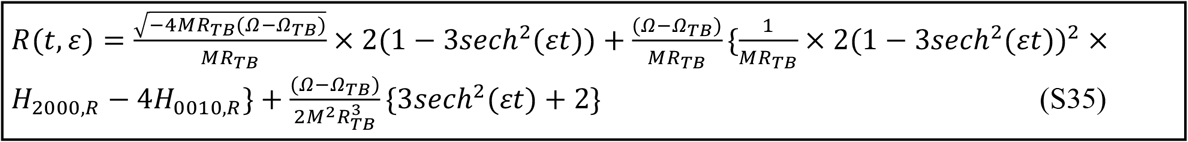

At *t* = 0, *sech*(*εt*) = 1

Hence,

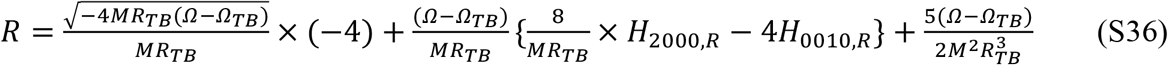

Here, *H*_2000_, *H*_0010_ are the normal form coefficients on *R*.

Now, we can compute *H*_2000,*R*_ from the following [4]:

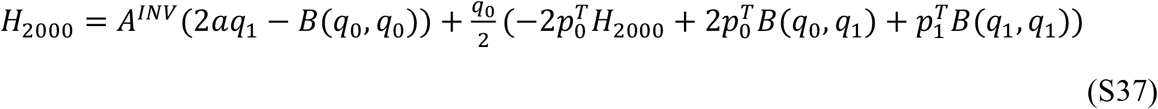

Here, 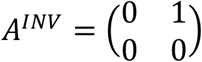

*p*_0_, *p*_1_ are the adjoint eigenvectors of (*A*^*INV*^)^*T*^

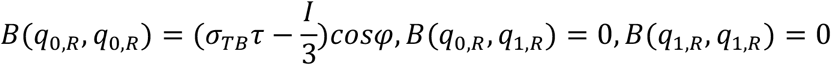

Putting these values in eq. (S33), we get,

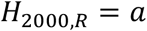

Now, we compute *H*_0010,*R*_ from the following two equations [4]

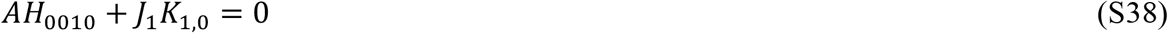

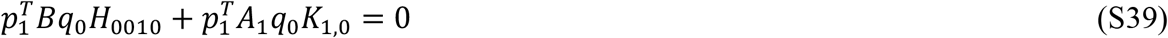

Here, 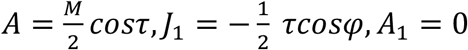

Putting these values, we get,

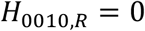

Now, substituting *H*_2000,*R*_ and *H*_0010,*R*_ in eq. (S32), we get,

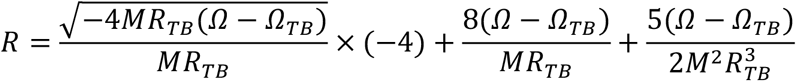

## Supplementary Material 2 - Plots

**Figure S1:**
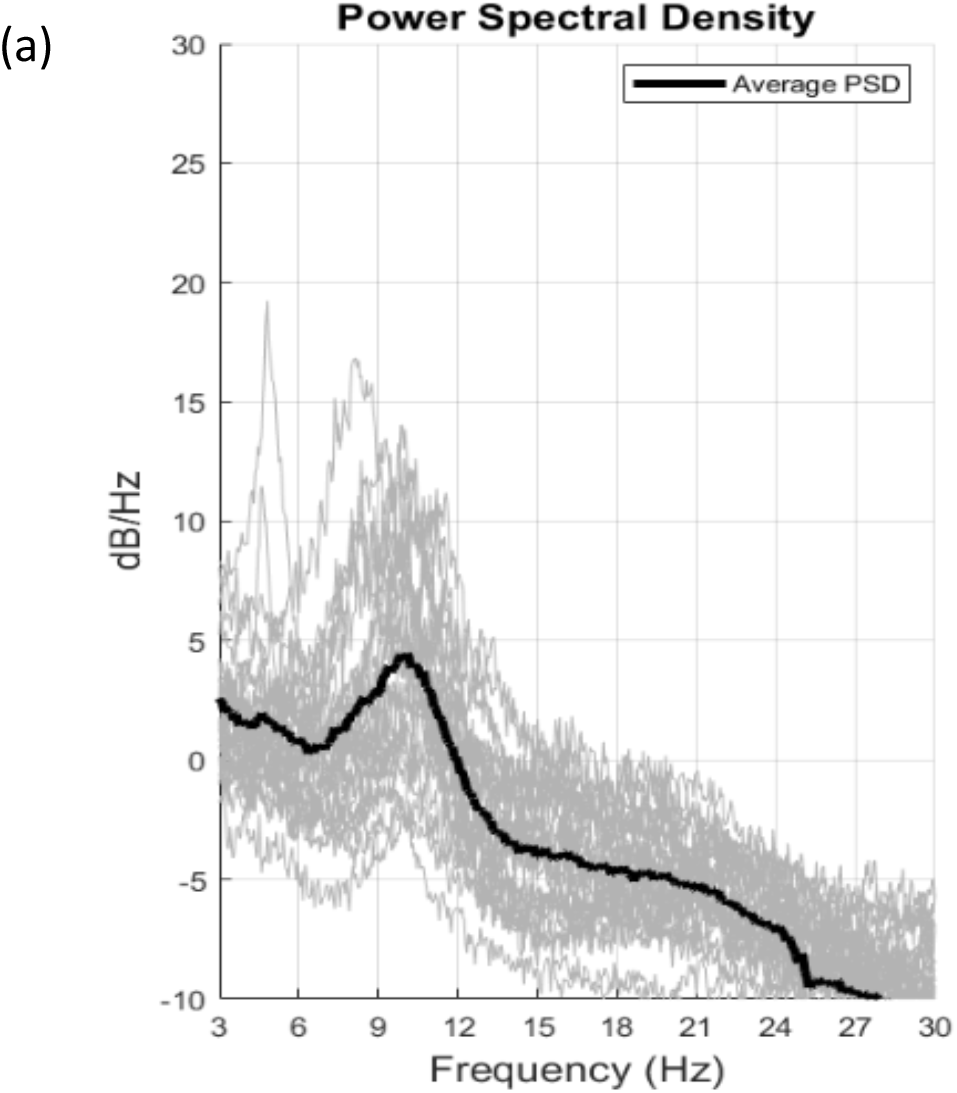
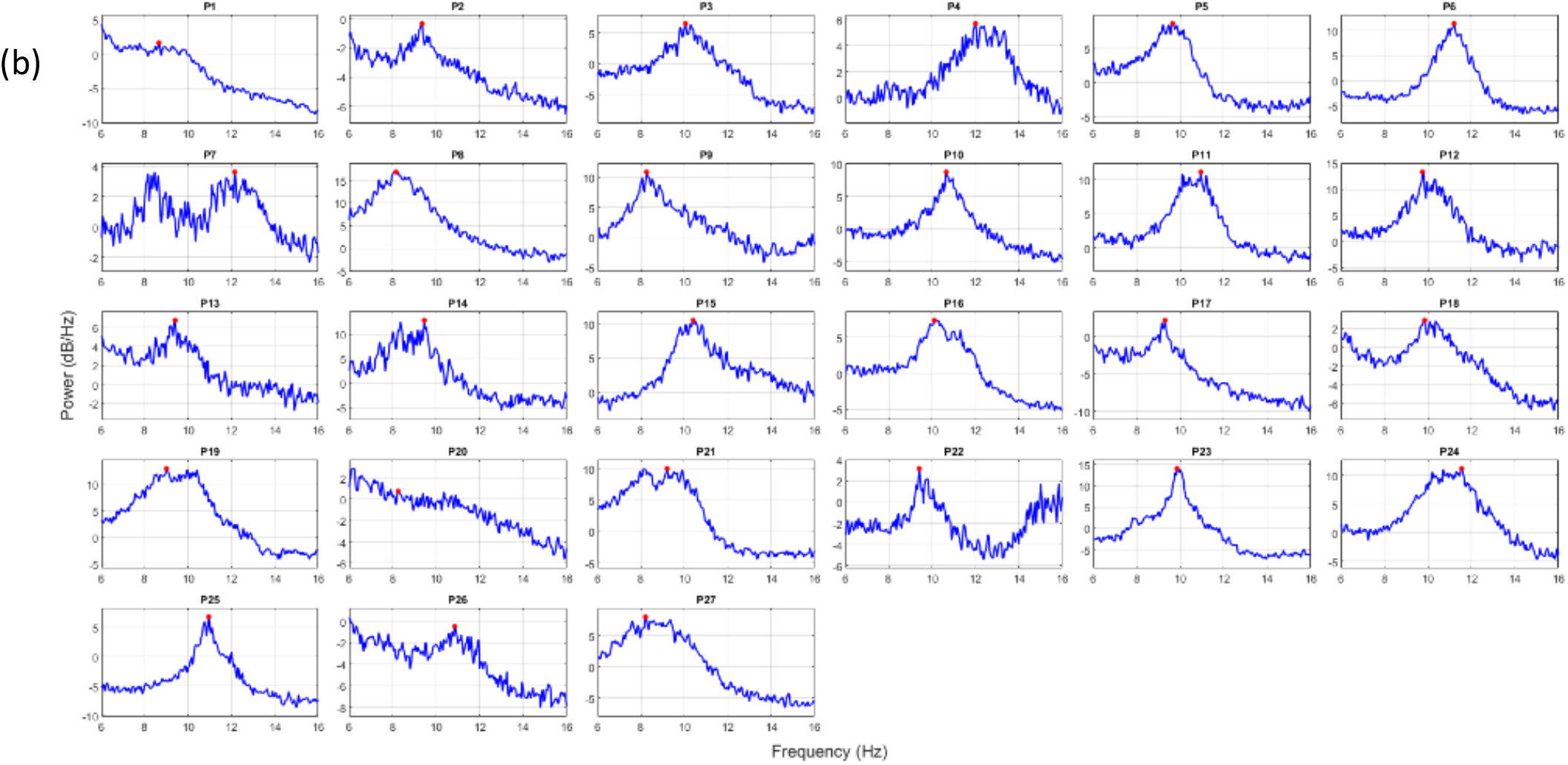
(a) Average IAF Power Spectral Density (PSD) denoted by black bold line. Light grey lines represent PSD for each of the 27 participants. (b) Each window shows the individual IAF PSD for each participant (IAF: 8.15–12.15Hz, mean: 9.9Hz). The Y-axis represents Power Spectral Density (in dB/Hz), and the X-axis represents Frequency in Hz.

**Figure S2:**
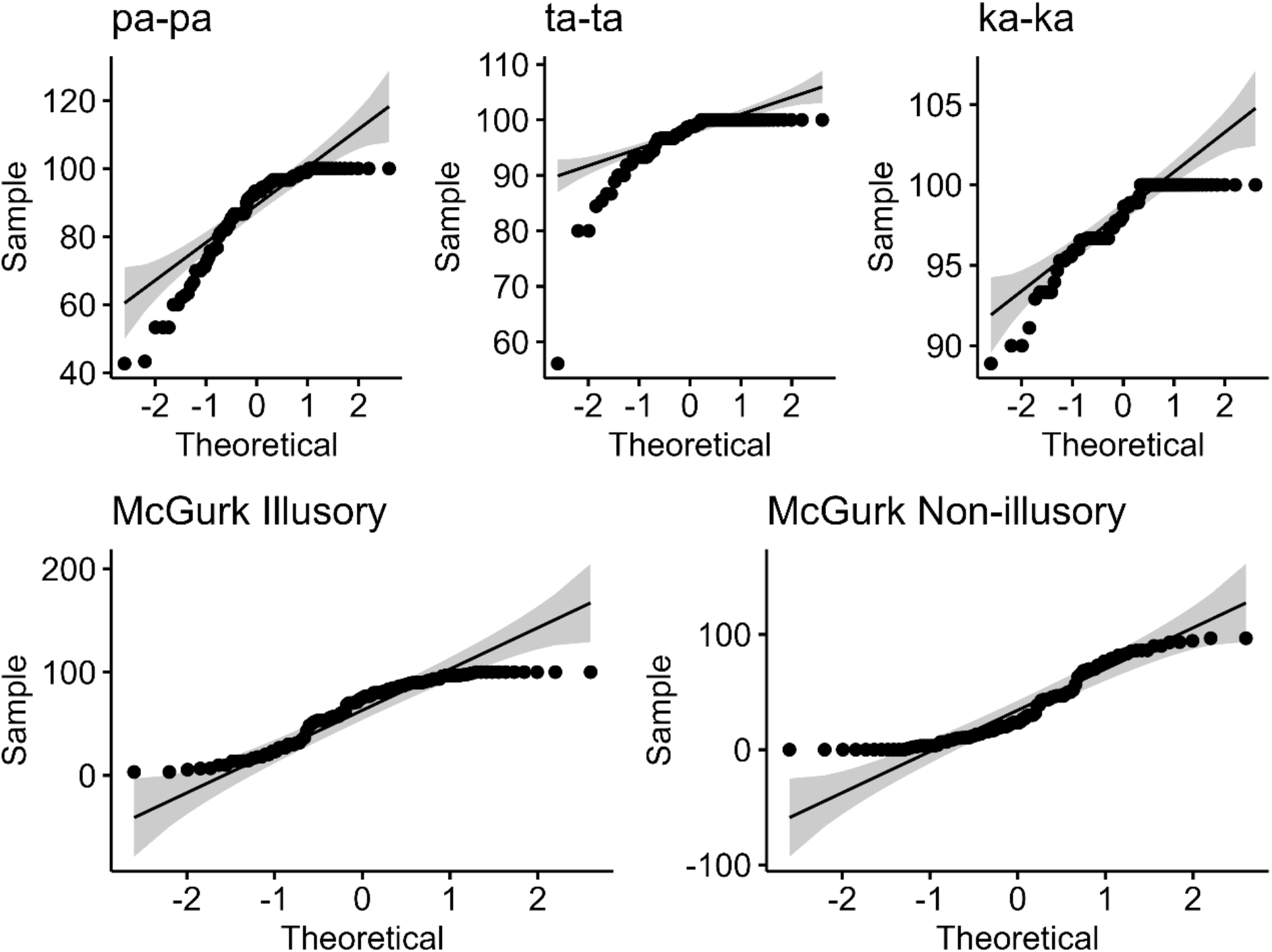
Q-Q plots assessing normality of the percentage perceptual response data for each stimulus condition (pa-pa, ta-ta, ka-ka, McGurk illusory, McGurk non-illusory). The X-axis (“Theoretical”) shows the theoretical quantiles expected under a standard normal distribution, and the Y-axis (“Sample”) shows the observed quantiles of the percentage perceptual response data. Points falling along the diagonal reference line indicate agreement with normality; the shaded region represents the 95% confidence band around this line. Deviations from the line, particularly the flattened plateaus near the upper quantiles in each panel, indicate departures from normality, consistent with ceiling effects in the response data.

**Figure S3:**
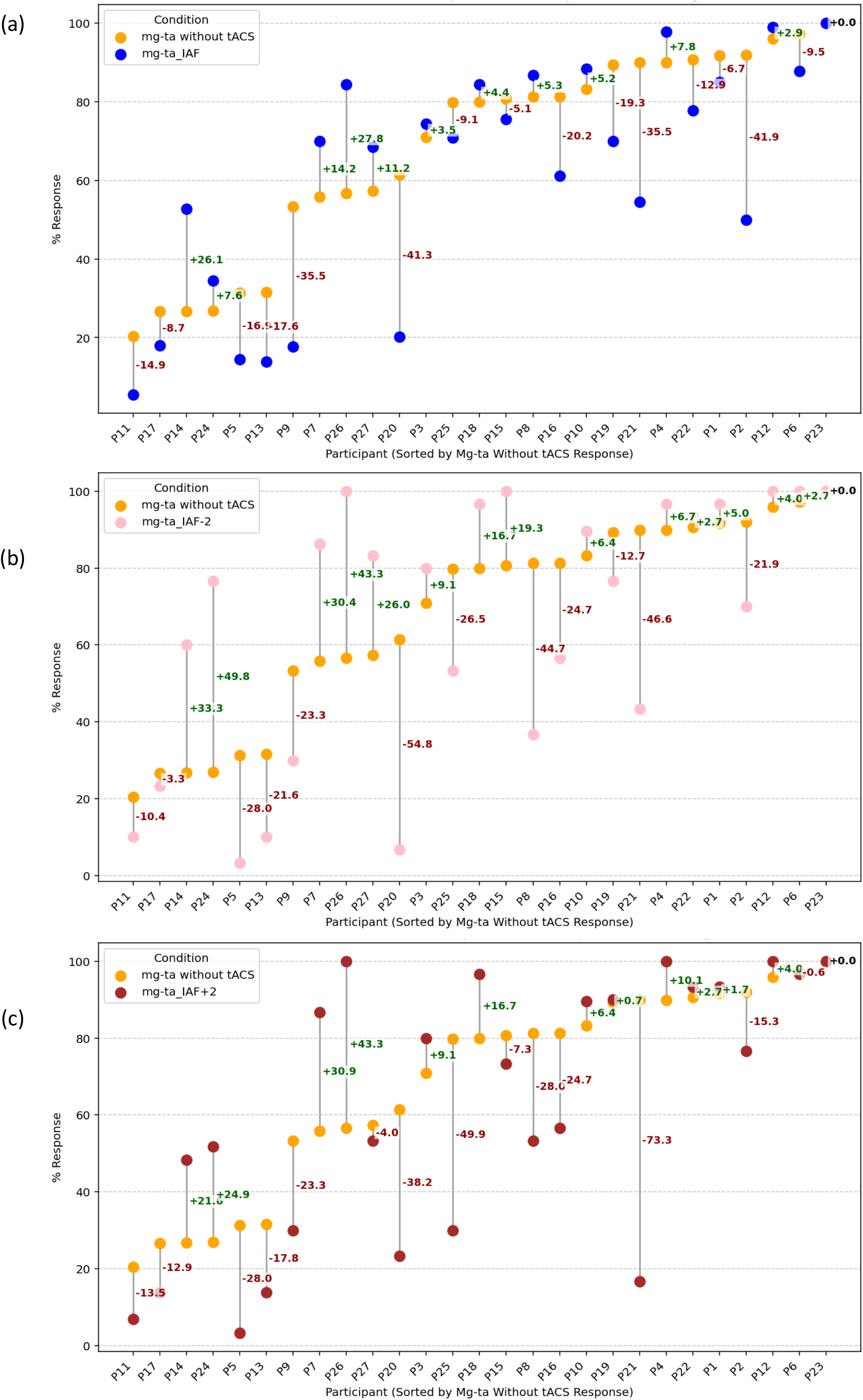
Individual-participant percentage of McGurk ‘ta’ responses (mg-ta) under transcranial alternating current stimulation (tACS) delivered at three frequencies relative to individual alpha frequency (IAF), compared against the no-stimulation baseline. **(A)** tACS at IAF, **(B)** tACS at IAF−2 Hz, **(C)** tACS at IAF+2 Hz. In each panel, orange markers show the %response without tACS and the second marker (blue, pink, or dark red in A, B, and C respectively) shows the %response under the corresponding tACS condition for the same participant; grey vertical lines connect paired values. Participants (x-axis, P1–P27) are ordered by ascending %response in the without-tACS condition, and this order is held constant across all three panels to facilitate visual comparison. Numeric labels denote the within-participant change in %response (tACS minus without tACS), colored green for an increase and dark red for a decrease under stimulation. N = 27 participants.

**Figure S4:**
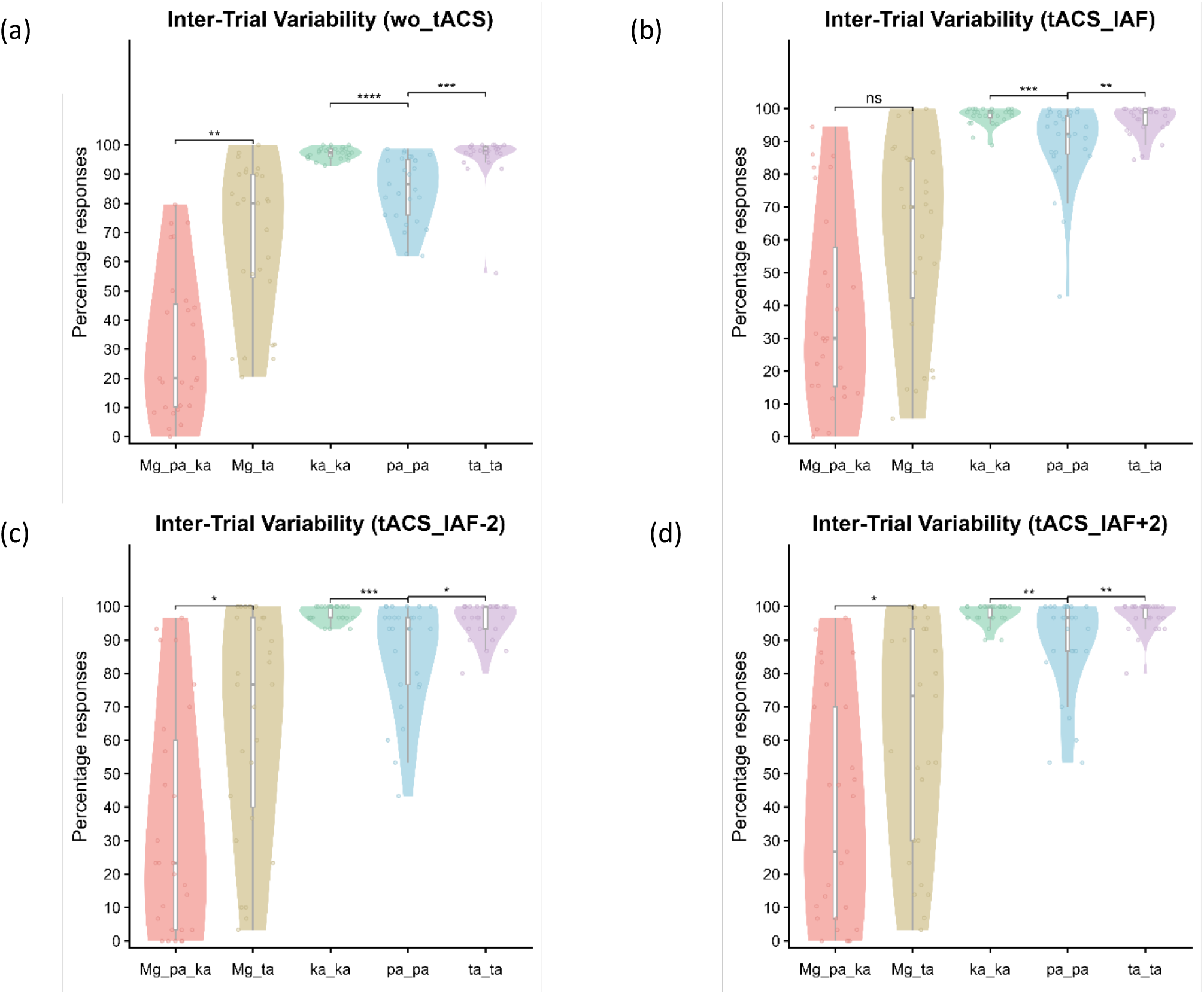
Inter-trial variability in percentage responses across all stimulation conditions. Violin plots (with overlaid boxplots and individual participant data points) showing the percentage of illusory (Mg_ta) and non-illusory/unisensory (Mg_pa_ka) percepts during incongruent McGurk trials, alongside the percentage of accurate perceptual responses for congruent audiovisual stimuli (ka_ka, pa_pa, ta_ta), across all four stimulation conditions: (a) baseline (wo_tACS), (b) tACS at individual alpha frequency (tACS_IAF), (c) tACS at IAF−2 Hz, and (d) tACS at IAF+2 Hz. Y-axis represents percentage responses; X-axis denotes response category. Horizontal brackets indicate results of post-hoc pairwise Wilcoxon signed-rank tests (FDR-corrected) for the Mg_pa_ka vs. Mg_ta, ka_ka vs. pa_pa, and pa_pa vs. ta_ta comparisons within each condition (ns = not significant, *p < 0.05, **p < 0.01, ***p < 0.001, ****p < 0.0001). Note: significance for the Mg_pa_ka vs. Mg_ta comparison shown here reflects FDR correction across all pairwise comparisons within each condition; the same contrast evaluated across stimulation conditions (Fig. 5) uses FDR correction within that comparison alone and yields a non-significant result at tACS_IAF+2 (p.adj = 0.053).

**Figure S5:**
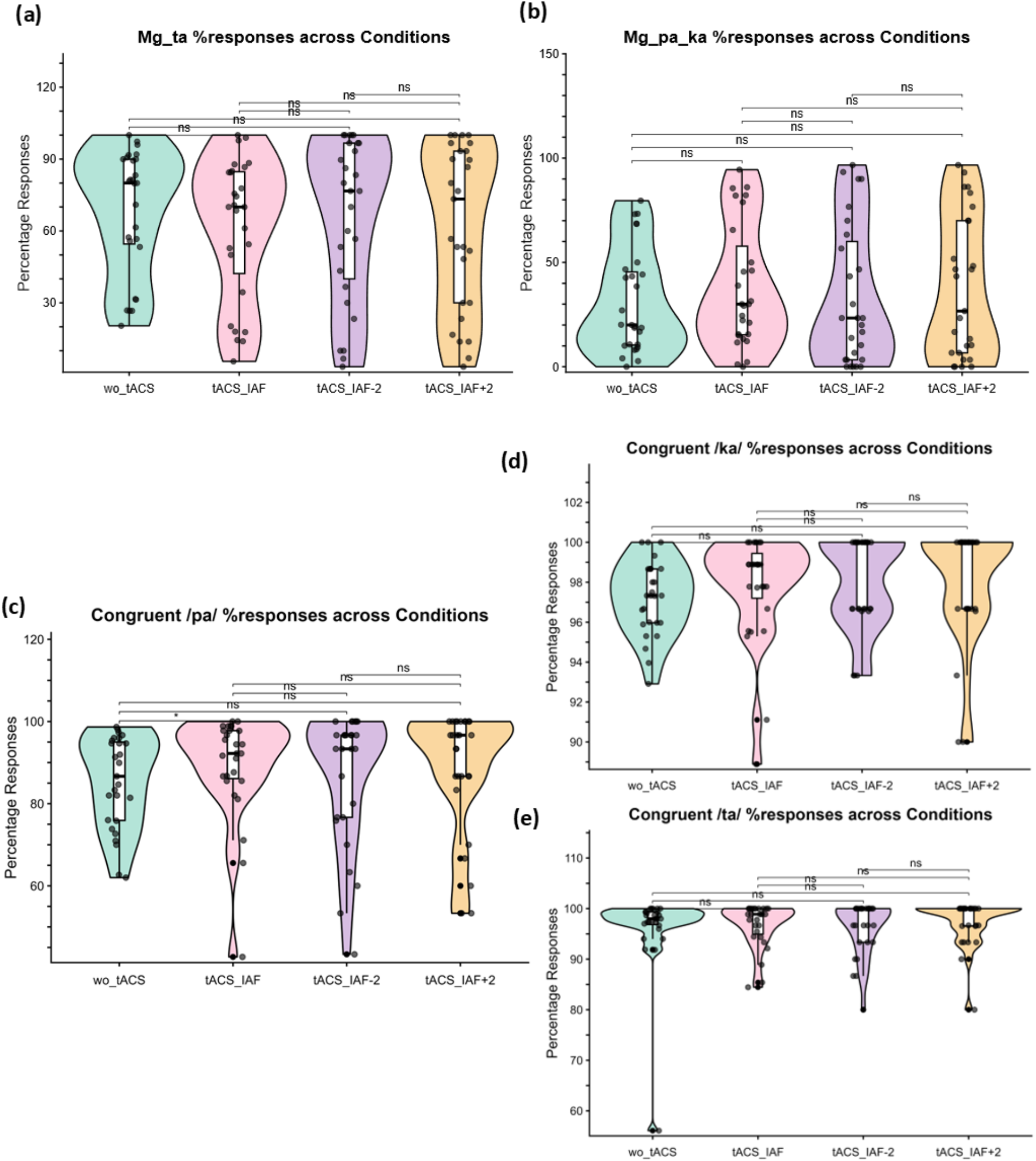
Percentage perceptual responses across tACS conditions for McGurk and congruent audiovisual stimuli. **(a)** Percentage of illusory McGurk ‘ta’ responses (Mg_ta). **(b)** Percentage of non-illusory McGurk ‘pa-ka’ responses (Mg_pa_ka). **(c)** Percentage of correct responses to the congruent /pa/ stimulus. **(d)** Percentage of correct responses to the congruent /ka/ stimulus. **(e)** Percentage of correct responses to the congruent /ta/ stimulus. In all panels, violin plots show the distribution of responses across participants for each condition (wo_tACS, tACS_IAF, tACS_IAF-2, tACS_IAF+2 Hz), with overlaid boxplots (median and interquartile range) and individual participant data points (black dots). Horizontal brackets indicate pairwise comparisons between conditions; ns = not significant, * = p < 0.05.

**Figure S6:**
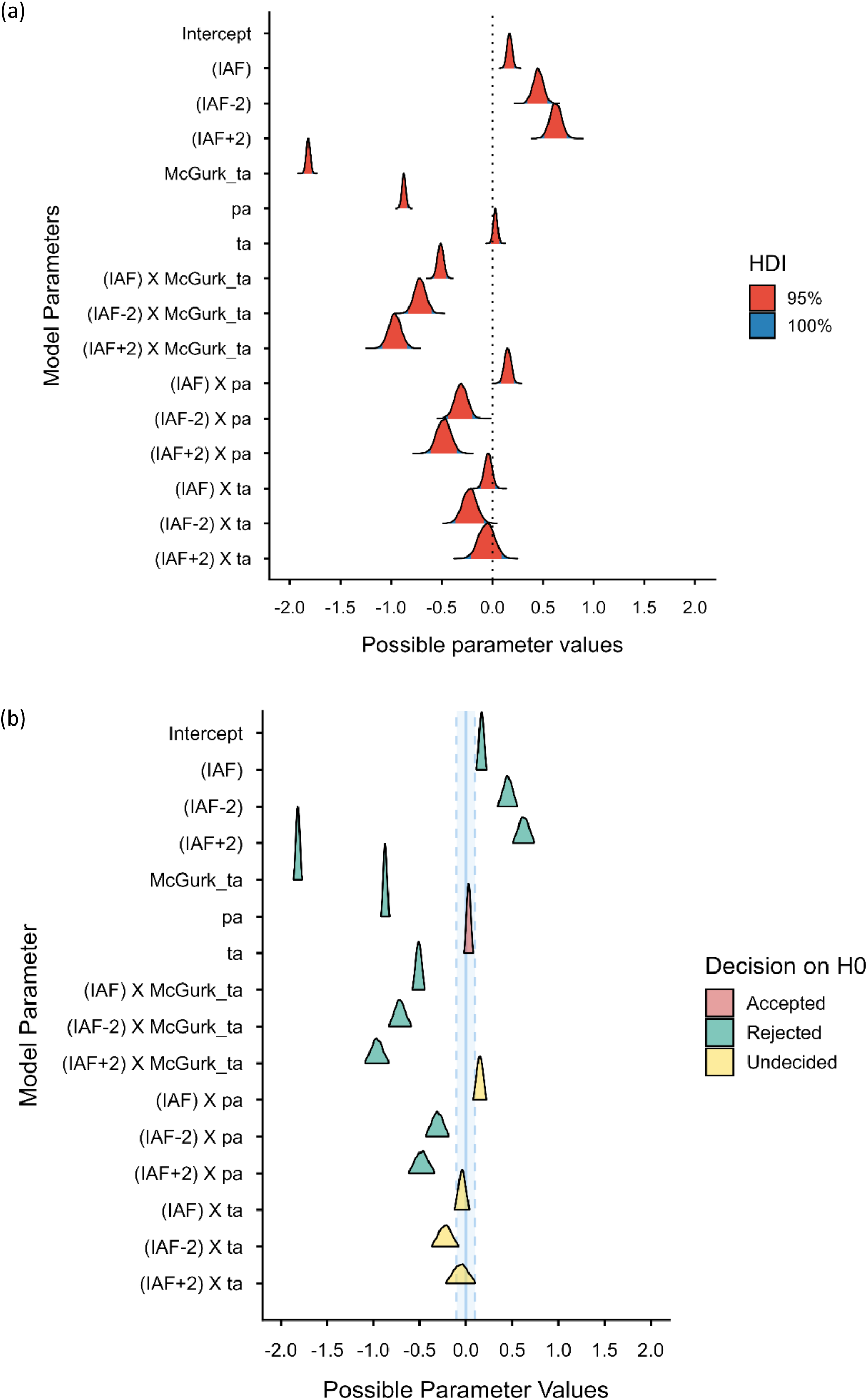
Posterior estimates of population-level parameters from the Bayesian ordered beta regression GLMM (Condition × Stimulus, random intercept for participant; reference level: wo_tACS, /ka/), on the logit scale. (a) Posterior distributions and 95% highest density intervals (HDI). A positive HDI (entirely right of zero) indicates a credible increase, and a negative HDI (entirely left of zero) a credible decrease, in response proportion relative to the reference condition; HDIs spanning zero indicate no credible effect. (b) Region of Practical Equivalence (ROPE) test for the same parameters (ROPE = [-0.1, 0.1]). Triangles mark the posterior median; colour indicates the decision on the null hypothesis rejected (teal), accepted (pink), or undecided (yellow). Parameter order matches across panels. N = 27 participants. Credible increases were observed for all three tACS conditions (IAF, IAF-2, IAF+2) relative to wo_tACS, whereas the McGurk illusory response showed a credible decrease relative to congruent /ka/ (Figure S6 (a)); tACS credibly reduced McGurk /ta/ responses in combination with IAF and IAF-2 stimulation (Figure S6 (b)).

**Figure S7:**
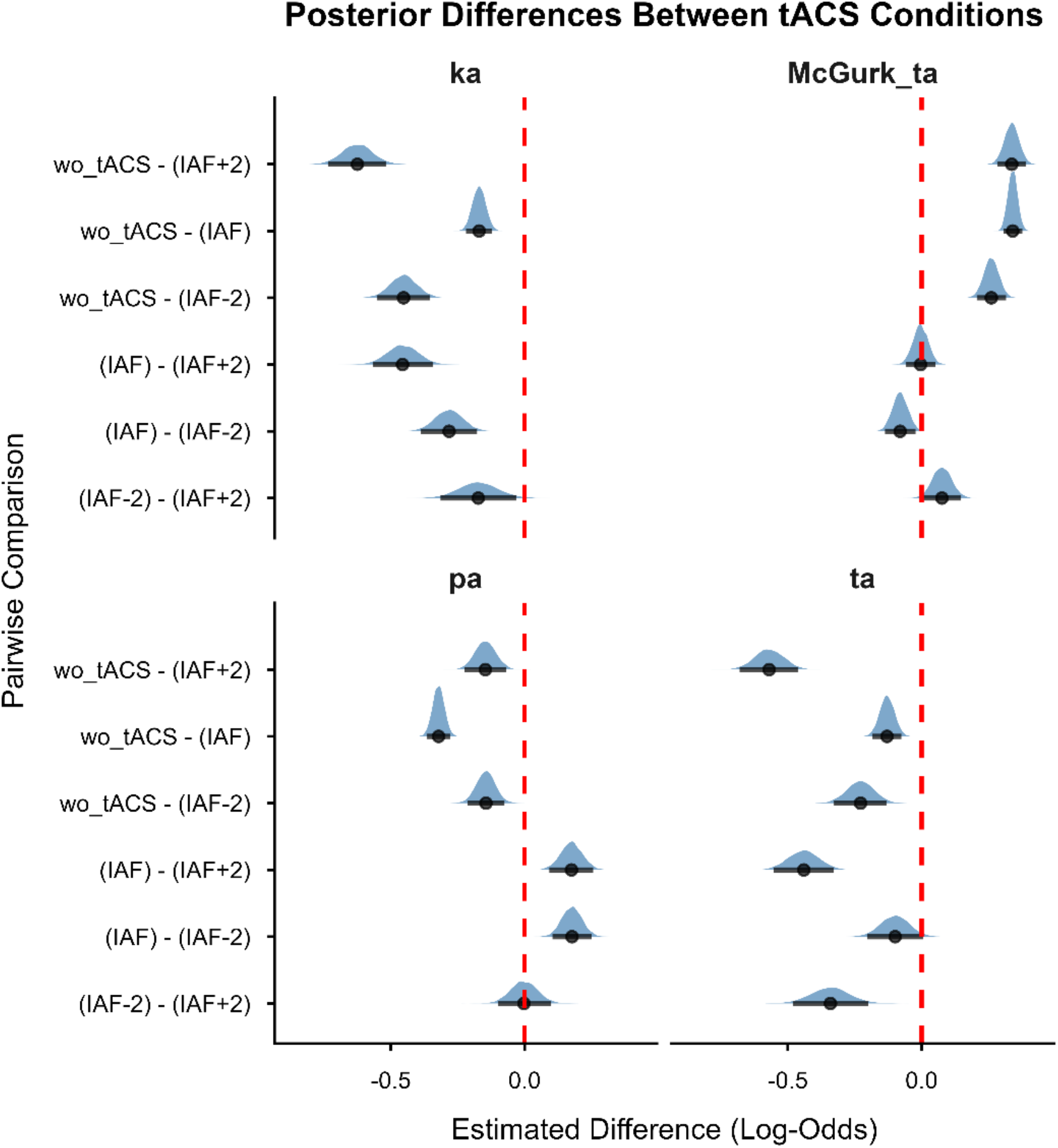
Post-hoc pairwise contrasts between tACS conditions from the Bayesian ordered beta regression GLMM, on the logit scale. Posterior differences between tACS conditions (wo_tACS, IAF, IAF−2, IAF+2) are estimated separately for each stimulus type (ka, McGurk_ta, pa, ta). Shaded densities show the full posterior distribution of each pairwise difference, with the median and 95% (HDI) marked. The dashed red line marks zero (no difference); a distribution lying entirely to one side of this line indicates a credible difference between the two tACS conditions being compared.

**Figure S8:**
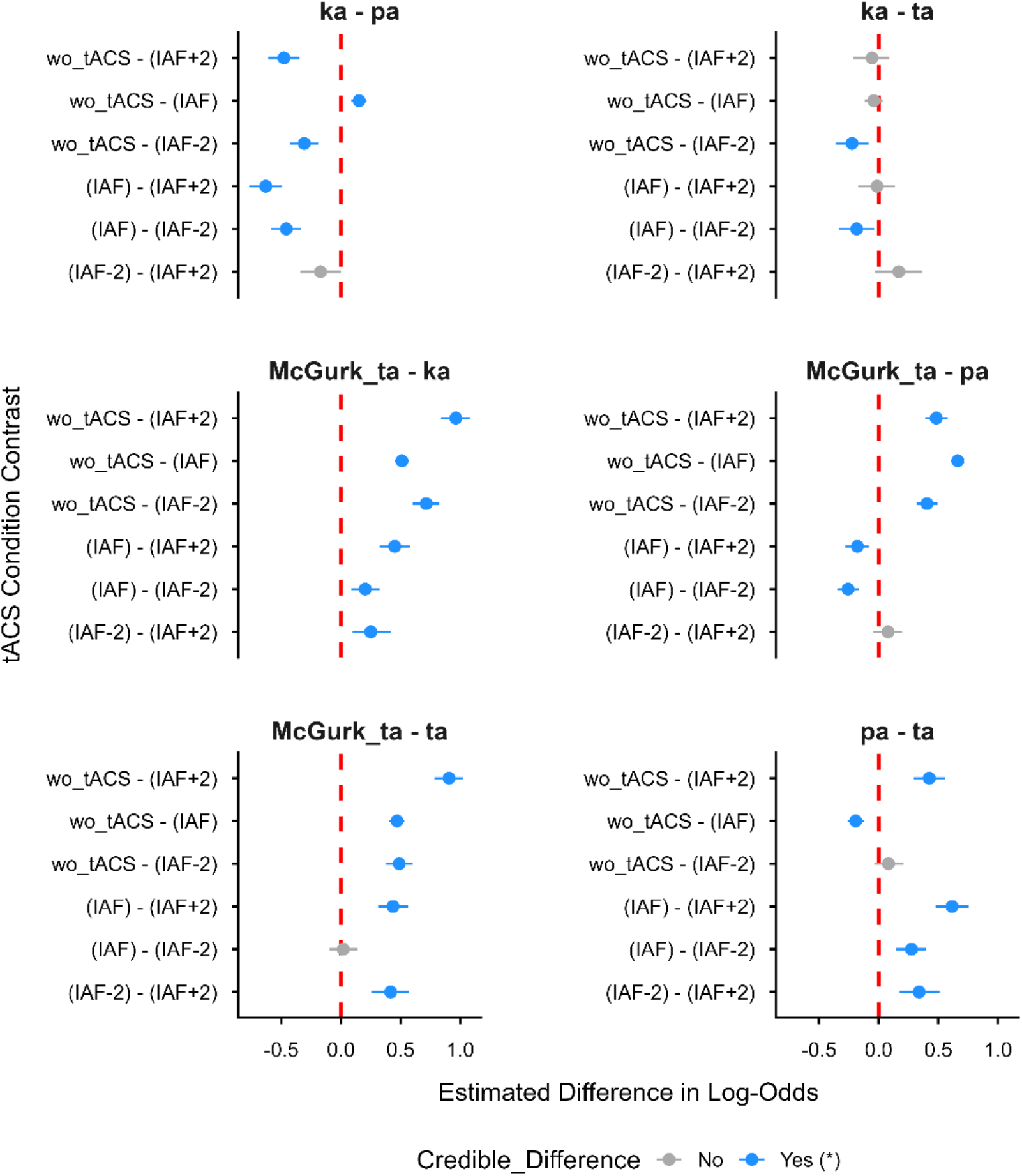
Interaction contrasts from the Bayesian ordered beta regression GLMM testing whether the effect of tACS condition differed between stimulus types, on the logit scale. Each panel shows a pairwise stimulus comparison (ka, McGurk_ta, pa, ta), and each row within a panel shows a pairwise tACS-condition comparison (wo_tACS, IAF, IAF−2, IAF+2). Points show the estimated interaction difference with 95% HDI; blue indicates a credible difference (interval excludes zero) and grey indicates no credible difference. The dashed red line marks zero (no difference).

## Supplementary Material 3 - Tables

**Table S1:** Descriptive statistics of perceptual responses at baseline and under generalized tACS.

| Variable | Median | IQR | Mean | SD | Coeff. of variation | Variance | Shapiro-Wilk | P-value of Shapiro-Wilk | Range | Min | Max |
| --- | --- | --- | --- | --- | --- | --- | --- | --- | --- | --- | --- |
| **pa** | 86.670 | 19.085 | 85.158 | 11.289 | 0.133 | 127.449 | 0.906 | 0.019 | 36.670 | 62.000 | 98.670 |
| **ta** | 98.000 | 2.490 | 96.070 | 8.322 | 0.087 | 69.259 | 0.418 | < .001 | 43.920 | 56.080 | 100.000 |
| **ka** | 97.330 | 2.695 | 97.212 | 1.856 | 0.019 | 3.444 | 0.964 | 0.459 | 7.090 | 92.910 | 100.000 |
| **Mg_pa_ka** | 20.000 | 35.075 | 30.853 | 24.835 | 0.805 | 616.755 | 0.887 | 0.007 | 79.590 | 0.000 | 79.590 |
| **Mg_ta** | 80.000 | 35.375 | 68.231 | 25.914 | 0.380 | 671.539 | 0.872 | 0.003 | 79.590 | 20.410 | 100.000 |
| **pa_tACS** | 93.330 | 14.195 | 88.516 | 12.503 | 0.141 | 156.316 | 0.815 | < .001 | 46.980 | 52.350 | 99.330 |
| **ta_tACS** | 98.650 | 4.370 | 96.737 | 4.085 | 0.042 | 16.686 | 0.790 | < .001 | 14.090 | 85.910 | 100.000 |
| **ka_tACS** | 98.670 | 2.335 | 97.837 | 2.055 | 0.021 | 4.223 | 0.830 | < .001 | 8.000 | 92.000 | 100.000 |
| **Mg_pa_ka_tACS** | 30.000 | 43.620 | 37.175 | 30.361 | 0.817 | 921.792 | 0.892 | 0.009 | 93.290 | 0.000 | 93.290 |
| **Mg_ta_tACS** | 70.000 | 43.620 | 62.603 | 30.243 | 0.483 | 914.666 | 0.897 | 0.012 | 93.290 | 6.710 | 100.000 |

The table presents the median, mean, standard deviation (SD), coefficient of variation, variance, and Shapiro-Wilk normality test results for percentage perceptual responses. Data are categorized by congruent syllables (/pa/, /ta/, /ka/) and incongruent McGurk stimuli yielding illusory (Mg\_ta) and non-illusory (Mg\_pa\_ka) percepts during baseline and generalized transcranial alternating current stimulation (tACS)

**Table S2.**
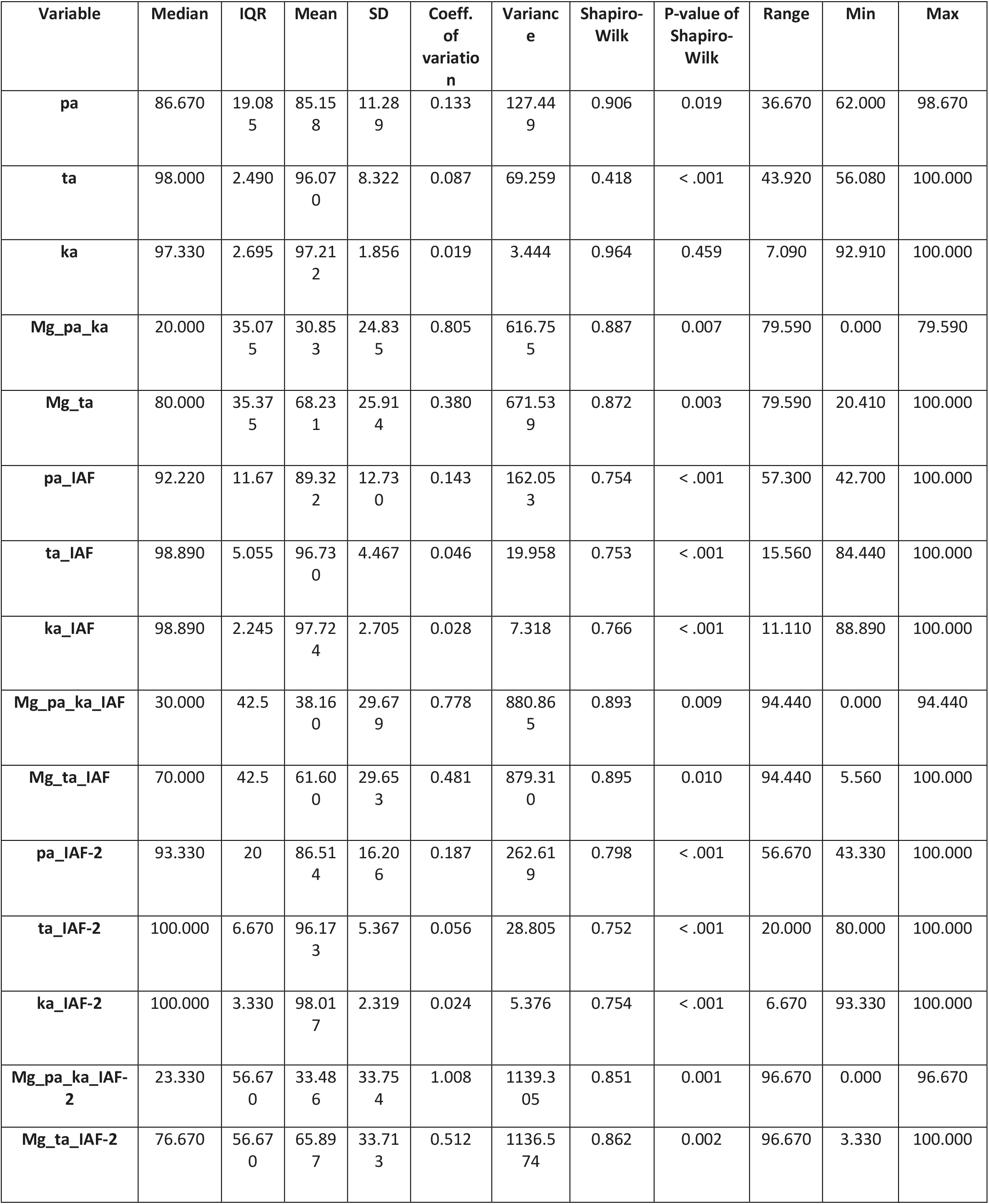
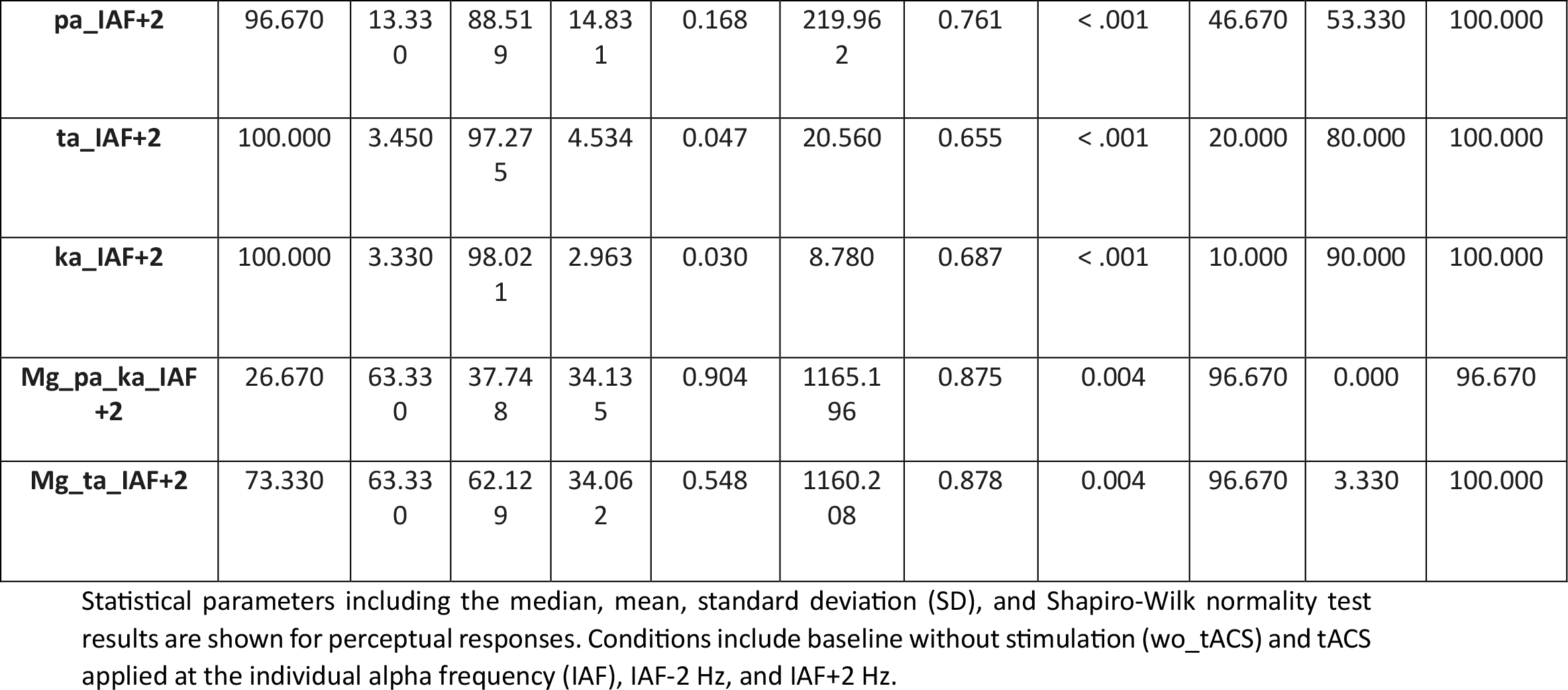
Descriptive statistics of perceptual responses across all specific tACS frequency conditions.

**Table S3.** Shapiro-Wilk normality test results for inter-trial variability.

| **Condition** | **response_type** | **variable** | **statistic** | **p** |
| --- | --- | --- | --- | --- |
| wo_tACS | Mg_pa_ka | value | 0.8868 | 0.0068 |
| wo_tACS | Mg_ta | value | 0.8717 | 0.0032 |
| wo_tACS | ka_ka | value | 0.9643 | 0.4593 |
| wo_tACS | pa_pa | value | 0.9063 | 0.0187 |
| wo_tACS | ta_ta | value | 0.4184 | <0.001 |
| tACS_IAF | Mg_pa_ka | value | 0.8931 | 0.0093 |
| tACS_IAF | Mg_ta | value | 0.8947 | 0.0101 |
| tACS_IAF | ka_ka | value | 0.7665 | <0.001 |
| tACS_IAF | pa_pa | value | 0.7539 | <0.001 |
| tACS_IAF | ta_ta | value | 0.7526 | <0.001 |
| tACS_IAF-2 | Mg_pa_ka | value | 0.8515 | 0.0012 |
| tACS_IAF-2 | Mg_ta | value | 0.8619 | 0.0020 |
| tACS_IAF-2 | ka_ka | value | 0.7538 | <0.001 |
| tACS_IAF-2 | pa_pa | value | 0.7978 | 0.0001 |
| tACS_IAF-2 | ta_ta | value | 0.7520 | <0.001 |
| tACS_IAF+2 | Mg_pa_ka | value | 0.8750 | 0.0038 |
| tACS_IAF+2 | Mg_ta | value | 0.8775 | 0.0042 |
| tACS_IAF+2 | ka_ka | value | 0.6866 | <0.001 |
| tACS_IAF+2 | pa_pa | value | 0.7611 | <0.001 |
| tACS_IAF+2 | ta_ta | value | 0.6547 | <0.001 |

Test statistics and corresponding p-values evaluating the normal distribution of perceptual responses to congruent and incongruent audio-visual stimuli. Data represent baseline (wo\_tACS) and active tACS conditions (IAF, IAF-2 Hz, IAF+2 Hz).

**Table S4.** Omnibus Friedman test results evaluating stimulation effects.

| Condition | Statistic($\chi^2$) | df1 | p | effect_size(W) | p_adj |
| --- | --- | --- | --- | --- | --- |
| wo_tACS | 81.101 | 4 | 0.000000000<br>000000102 | 0.751 | 0.000000000000000<br>407 |
| tACS_IAF | 76.949 | 4 | 0.000000000<br>000000771 | 0.712 | 0.000000000000001<br>540 |
| tACS_IAF-2 | 61.916 | 4 | 0.000000000<br>001150000 | 0.573 | 0.000000000001150<br>000 |
| tACS_IAF+2 | 64.710 | 4 | 0.000000000<br>000296000 | 0.599 | 0.000000000000395<br>000 |

The table details the test statistic, degrees of freedom, effect sizes, and p-values for perceptual responses across baseline (wo\_tACS) and active tACS conditions. P-values are adjusted using the False Discovery Rate (FDR) method across conditions.

**Table S5.** Post-hoc pairwise Wilcoxon signed-rank comparisons within s@mula@on condi@ons.

| Condition | group1 | group2 | n1 | n2 | Statistic(V) | p | p_adj | star |
| --- | --- | --- | --- | --- | --- | --- | --- | --- |
| wo_tACS | Mg_pa_ka | Mg_ta | 27 | 27 | 54.0 | 0.0010 | 0.0020 | ** |
| wo_tACS | Mg_pa_ka | ka_ka | 27 | 27 | 0.0 | 0.0000 | 0.0000 | **** |
| wo_tACS | Mg_pa_ka | pa_pa | 27 | 27 | 0.0 | 0.0000 | 0.0000 | **** |
| wo_tACS | Mg_pa_ka | ta_ta | 27 | 27 | 0.0 | 0.0000 | 0.0000 | **** |
| wo_tACS | Mg_ta | ka_ka | 27 | 27 | 3.0 | 0.0000 | 0.0000 | **** |
| wo_tACS | Mg_ta | pa_pa | 27 | 27 | 96.5 | 0.0460 | 0.0510 | ns |
| wo_tACS | Mg_ta | ta_ta | 27 | 27 | 1.0 | 0.0000 | 0.0000 | **** |
| wo_tACS | ka_ka | pa_pa | 27 | 27 | 348.0 | 0.0000 | 0.0000 | **** |
| wo_tACS | ka_ka | ta_ta | 27 | 27 | 132.5 | 0.6270 | 0.6270 | ns |
| wo_tACS | pa_pa | ta_ta | 27 | 27 | 32.0 | 0.0003 | 0.0004 | *** |
| tACS_IAF | Mg_pa_ka | Mg_ta | 27 | 27 | 103.0 | 0.0670 | 0.0750 | ns |
| tACS_IAF | Mg_pa_ka | ka_ka | 27 | 27 | 0.0 | 0.0000 | 0.0000 | **** |
| tACS_IAF | Mg_pa_ka | pa_pa | 27 | 27 | 1.0 | 0.0000 | 0.0000 | **** |
| tACS_IAF | Mg_pa_ka | ta_ta | 27 | 27 | 0.0 | 0.0000 | 0.0000 | **** |
| tACS_IAF | Mg_ta | ka_ka | 27 | 27 | 3.0 | 0.0000 | 0.0000 | **** |
| tACS_IAF | Mg_ta | pa_pa | 27 | 27 | 47.0 | 0.0003 | 0.0004 | *** |
| tACS_IAF | Mg_ta | ta_ta | 27 | 27 | 0.0 | 0.0000 | 0.0000 | **** |
| tACS_IAF | ka_ka | pa_pa | 27 | 27 | 305.0 | 0.0001 | 0.0002 | *** |
| tACS_IAF | ka_ka | ta_ta | 27 | 27 | 144.0 | 0.5780 | 0.5780 | ns |
| tACS_IAF | pa_pa | ta_ta | 27 | 27 | 57.0 | 0.0020 | 0.0020 | ** |
| tACS_IAF-2 | Mg_pa_ka | Mg_ta | 27 | 27 | 91.0 | 0.0190 | 0.0240 | * |
| tACS_IAF-2 | Mg_pa_ka | ka_ka | 27 | 27 | 0.0 | 0.0000 | 0.0000 | **** |
| tACS_IAF-2 | Mg_pa_ka | pa_pa | 27 | 27 | 5.0 | 0.0000 | 0.0001 | **** |
| tACS_IAF-2 | Mg_pa_ka | ta_ta | 27 | 27 | 2.0 | 0.0000 | 0.0000 | **** |
| tACS_IAF-2 | Mg_ta | ka_ka | 27 | 27 | 5.5 | 0.0001 | 0.0002 | *** |
| tACS_IAF-2 | Mg_ta | pa_pa | 27 | 27 | 92.0 | 0.0350 | 0.0390 | * |
| tACS_IAF-2 | Mg_ta | ta_ta | 27 | 27 | 7.0 | 0.0001 | 0.0002 | *** |
| tACS_IAF-2 | ka_ka | pa_pa | 27 | 27 | 252.0 | 0.0005 | 0.0009 | *** |
| tACS_IAF-2 | ka_ka | ta_ta | 27 | 27 | 95.0 | 0.1680 | 0.1680 | ns |
| tACS_IAF-2 | pa_pa | ta_ta | 27 | 27 | 47.0 | 0.0100 | 0.0140 | * |
| tACS_IAF+2 | Mg_pa_ka | Mg_ta | 27 | 27 | 103.0 | 0.0400 | 0.0440 | * |
| tACS_IAF+2 | Mg_pa_ka | ka_ka | 27 | 27 | 0.0 | 0.0000 | 0.0000 | **** |
| tACS_IAF+2 | Mg_pa_ka | pa_pa | 27 | 27 | 7.0 | 0.0000 | 0.0001 | **** |
| tACS_IAF+2 | Mg_pa_ka | ta_ta | 27 | 27 | 2.0 | 0.0000 | 0.0000 | **** |
| tACS_IAF+2 | Mg_ta | ka_ka | 27 | 27 | 2.0 | 0.0000 | 0.0001 | **** |
| tACS_IAF+2 | Mg_ta | pa_pa | 27 | 27 | 50.0 | 0.0040 | 0.0060 | ** |
| tACS_IAF+2 | Mg_ta | ta_ta | 27 | 27 | 0.0 | 0.0000 | 0.0001 | **** |
| tACS_IAF+2 | ka_ka | pa_pa | 27 | 27 | 170.0 | 0.0030 | 0.0040 | ** |
| tACS_IAF+2 | ka_ka | ta_ta | 27 | 27 | 79.0 | 0.5840 | 0.5840 | ns |
| tACS_IAF+<br>2 | pa_pa | ta_ta | 27 | 27 | 28.0 | 0.0040 | 0.0060 | ** |

Comparisons of perceptual responses across specific syllable groups within each targeted stimulation condition. The results utilize the Wilcoxon signed-rank test to compare illusory and non-illusory percepts, with p-values adjusted via the False Discovery Rate (FDR) method.
Note: FDR correction (Benjamini–Hochberg) was applied separately within each stimulation condition across all ten pairwise comparisons among the five response categories (Mg\_pa\_ka, Mg\_ta, ka\_ka, pa\_pa, ta\_ta). Adjusted p-values therefore differ from those in Table S6, which corrects only across the four focal Mg\_ta vs Mg\_pa\_ka comparisons.

**Table S6.** Wilcoxon signed-rank test results comparing illusory and non-illusory responses to the McGurk stimulus.

| Condition | Statistic (V) | p_value | Eff. size | p_adj(fdr) | Magnitude |
| --- | --- | --- | --- | --- | --- |
| wo_tACS | 324 | 0.001 | 0.6242103 | 0.005 | large |
| tACS_IAF | 248 | 0.067 | 0.3630060 | 0.067 | moderate |
| tACS_IAF+2 | 275 | 0.039 | 0.3979179 | 0.053 | moderate |
| tACS_IAF-2 | 287 | 0.019 | 0.4537365 | 0.038 | moderate |

Statistical comparisons between illusory (Mg\_ta) and non-illusory (Mg\_pa\_ka) responses across experimental conditions. The table includes the test statistic (V), effect sizes, magnitude interpretations, and FDR-adjusted p-values for baseline and tACS interventions (IAF, IAF-2 Hz, IAF+2 Hz).
Note: FDR correction (Benjamini–Hochberg) was applied across the four focal comparisons of illusory (Mg\_ta) versus non-illusory (Mg\_pa\_ka) responses, one per stimulation condition (wo\_tACS, tACS\_IAF, tACS\_IAF-2, tACS\_IAF+2). This is the pre-specified comparison of interest and is the one reported in the main text and Figure 5.

**Table S7.** Friedman test results for specific perceptual responses across stimulation conditions.

| Response | n | statistic($\chi^2$) | df | p | p_adj | Kendall_W |
| --- | --- | --- | --- | --- | --- | --- |
| Mg_ta | 27 | 3.8366534 | 3 | 0.279 | 0.360 | 0.047 |
| Mg_pa_ka | 27 | 3.7619048 | 3 | 0.288 | 0.360 | 0.046 |
| Congruent /pa/ | 27 | 10.5807692 | 3 | 0.014 | 0.071 | 0.131 |
| Congruent /ta/ | 27 | 0.6490385 | 3 | 0.885 | 0.885 | 0.008 |
| Congruent /ka/ | 27 | 6.5648536 | 3 | 0.087 | 0.218 | 0.081 |

Omnibus non-parametric evaluation of the percentage response for congruent and incongruent stimuli, using stimulation type as the independent variable. The table displays test statistics, degrees of freedom, adjusted p-values, and Kendall's W for effect size across baseline and active tACS conditions.

**Table S8.** Post-hoc pairwise Wilcoxon tests comparing specific perceptual responses across stimulation conditions.

| Response | group1 | group2 | Statistic (V) | p | p_adj | star | Eff. size | magnitude |
| --- | --- | --- | --- | --- | --- | --- | --- | --- |
| Mg_ta | wo_tAC<br>S | tACS_IA<br>F | 246.0 | 0.075 | 0.226 | ns | 0.3352<br>3615 | moderate |
| Mg_ta | wo_tAC<br>S | tACS_IA<br>F-2 | 190.0 | 0.722 | 0.722 | ns | 0.0624<br>2328 | small |
| Mg_ta | wo_tAC<br>S | tACS_IA<br>F+2 | 215.0 | 0.322 | 0.449 | ns | 0.1872<br>7660 | small |
| Mg_ta | tACS_IA<br>F | tACS_IA<br>F-2 | 98.5 | 0.052 | 0.226 | ns | 0.3792<br>0469 | moderate |
| Mg_ta | tACS_IA<br>F | tACS_IA<br>F+2 | 140.0 | 0.374 | 0.449 | ns | 0.1780<br>2195 | small |
| Mg_ta | tACS_IA<br>F-2 | tACS_IA<br>F+2 | 102.5 | 0.227 | 0.449 | ns | 0.1950<br>7891 | small |
| Mg_pa_ka | wo_tAC<br>S | tACS_IA<br>F | 95.0 | 0.042 | 0.147 | ns | 0.3814<br>7561 | moderate |
| Mg_pa_ka | wo_tAC<br>S | tACS_IA<br>F-2 | 160.0 | 0.703 | 0.703 | ns | 0.0670<br>4723 | small |
| Mg_pa_ka | wo_tAC<br>S | tACS_IA<br>F+2 | 126.0 | 0.213 | 0.321 | ns | 0.2381<br>4185 | small |
| Mg_pa_ka | tACS_IA<br>F | tACS_IA<br>F-2 | 253.5 | 0.049 | 0.147 | ns | 0.3838<br>1529 | moderate |
| Mg_pa_ka | tACS_IA<br>F | tACS_IA<br>F+2 | 208.5 | 0.409 | 0.491 | ns | 0.1664<br>6809 | small |
| Mg_pa_ka | tACS_IA<br>F-2 | tACS_IA<br>F+2 | 56.5 | 0.214 | 0.321 | ns | 0.1794<br>3902 | small |
| Congruent<br>/pa/ | wo_tAC<br>S | tACS_IA<br>F | 66.0 | 0.002 | 0.013 | * | 0.5687<br>0443 | large |
| Congruent<br>/pa/ | wo_tAC<br>S | tACS_IA<br>F-2 | 149.0 | 0.509 | 0.611 | ns | 0.1318<br>8245 | small |
| Congruent<br>/pa/ | wo_tAC<br>S | tACS_IA<br>F+2 | 105.0 | 0.045 | 0.134 | ns | 0.3884<br>9565 | moderate |
| Congruent<br>/pa/ | tACS_IA<br>F | tACS_IA<br>F-2 | 220.0 | 0.264 | 0.396 | ns | 0.2150<br>4457 | small |
| Congruent<br>/pa/ | tACS_IA<br>F | tACS_IA<br>F+2 | 152.5 | 0.798 | 0.798 | ns | 0.0508<br>9280 | small |
| Congruent<br>/pa/ | tACS_IA<br>F-2 | tACS_IA<br>F+2 | 81.5 | 0.240 | 0.396 | ns | 0.2570<br>0163 | small |
| Congruent<br>/ta/ | wo_tAC<br>S | tACS_IA<br>F | 148.0 | 0.773 | 0.913 | ns | 0.0370<br>7326 | small |
| Congruent<br>/ta/ | wo_tAC<br>S | tACS_IA<br>F-2 | 163.0 | 0.456 | 0.794 | ns | 0.1251<br>6762 | small |
| Congruent<br>/ta/ | wo_tAC<br>S | tACS_IA<br>F+2 | 102.0 | 0.436 | 0.794 | ns | 0.1601<br>9805 | small |
| Congruent<br>/ta/ | tACS_IA<br>F | tACS_IA<br>F-2 | 102.5 | 0.472 | 0.794 | ns | 0.0803<br>1465 | small |
| Congruent<br>/ta/ | tACS_IA<br>F | tACS_IA<br>F+2 | 82.5 | 0.913 | 0.913 | ns | 0.0567<br>8259 | small |
| Congruent<br>/ta/ | tACS_IA<br>F-2 | tACS_IA<br>F+2 | 42.0 | 0.529 | 0.794 | ns | 0.1157<br>5442 | small |
| Congruent<br>/ka/ | wo_tAC<br>S | tACS_IA<br>F | 122.0 | 0.178 | 0.356 | ns | 0.2658<br>9613 | small |
| Congruent<br>/ka/ | wo_tAC<br>S | tACS_IA<br>F-2 | 99.5 | 0.092 | 0.345 | ns | 0.3239<br>5692 | moderate |
| Congruent<br>/ka/ | wo_tAC<br>S | tACS_IA<br>F+2 | 103.5 | 0.115 | 0.345 | ns | 0.3238<br>6329 | moderate |
| Congruent<br>/ka/ | tACS_IA<br>F | tACS_IA<br>F-2 | 112.0 | 0.648 | 0.972 | ns | 0.1372<br>9100 | small |
| Congruent<br>/ka/ | tACS_IA<br>F | tACS_IA<br>F+2 | 131.5 | 0.855 | 1.000 | ns | 0.0580<br>7025 | small |
| Congruent<br>/ka/ | tACS_IA<br>F-2 | tACS_IA<br>F+2 | 76.0 | 1.000 | 1.000 | ns | 0.0263<br>3667 | small |

Pairwise comparisons evaluating changes in individual response types (e.g., Mg\_ta, congruent /pa/) between the baseline (wo\_tACS) and active tACS frequencies. The table reports test statistics, adjusted p-values, and effect size magnitudes to outline the frequency-specific modulations of multisensory integration.

**Table S9.** Highest Density Interval (HDI) and Region of Practical Equivalence (ROPE) results for Generalised Linear Mixed Model (GLMM) parameters.

| Parameter | Median | 95% HDI Lower | 95% HDI Upper | % in ROPE | Decision on H0 |
| --- | --- | --- | --- | --- | --- |
| Intercept | 2.595 | 2.373 | 2.835 | 0.000 | Rejected |
| (IAF) | 0.170 | 0.121 | 0.219 | 0.000 | Rejected |
| (IAF-2) | 0.452 | 0.353 | 0.552 | 0.000 | Rejected |
| (IAF+2) | 0.625 | 0.517 | 0.734 | 0.000 | Rejected |
| McGurk_ta | -1.817 | -1.855 | -1.780 | 0.000 | Rejected |
| pa | -0.873 | -0.910 | -0.835 | 0.000 | Rejected |
| ta | 0.030 | -0.012 | 0.072 | 1.000 | Accepted |
| (IAF) X McGurk_ta | -0.510 | -0.571 | -0.450 | 0.000 | Rejected |
| (IAF-2) X McGurk_ta | -0.714 | -0.825 | -0.600 | 0.000 | Rejected |
| (IAF+2) X McGurk_ta | -0.963 | -1.085 | -0.839 | 0.000 | Rejected |
| (IAF) X pa | 0.152 | 0.087 | 0.216 | 0.038 | Undecided |
| (IAF-2) X pa | -0.308 | -0.428 | -0.191 | 0.000 | Rejected |
| (IAF+2) X pa | -0.478 | -0.610 | -0.347 | 0.000 | Rejected |
| (IAF) X ta | -0.041 | -0.114 | 0.031 | 0.965 | Undecided |
| (IAF-2) X ta | -0.224 | -0.363 | -0.085 | 0.017 | Undecided |
| (IAF+2) X ta | -0.055 | -0.209 | 0.096 | 0.722 | Undecided |
| Note. Estimates are on the logit scale. ROPE = [-0.1, 0.1]. H0 = practical equivalence to zero (no effect). |  |  |  |  |  |

Bayesian parameter estimates detailing the modulatory influence of tACS on stimulus perception. The table presents the median estimates on the logit scale, 95% HDI bounds, the percentage of the posterior within the ROPE [-0.1, 0.1], and the decision on the null hypothesis of practical equivalence to zero. Reference (wo\_tACS, /ka/)

**Table S10.**
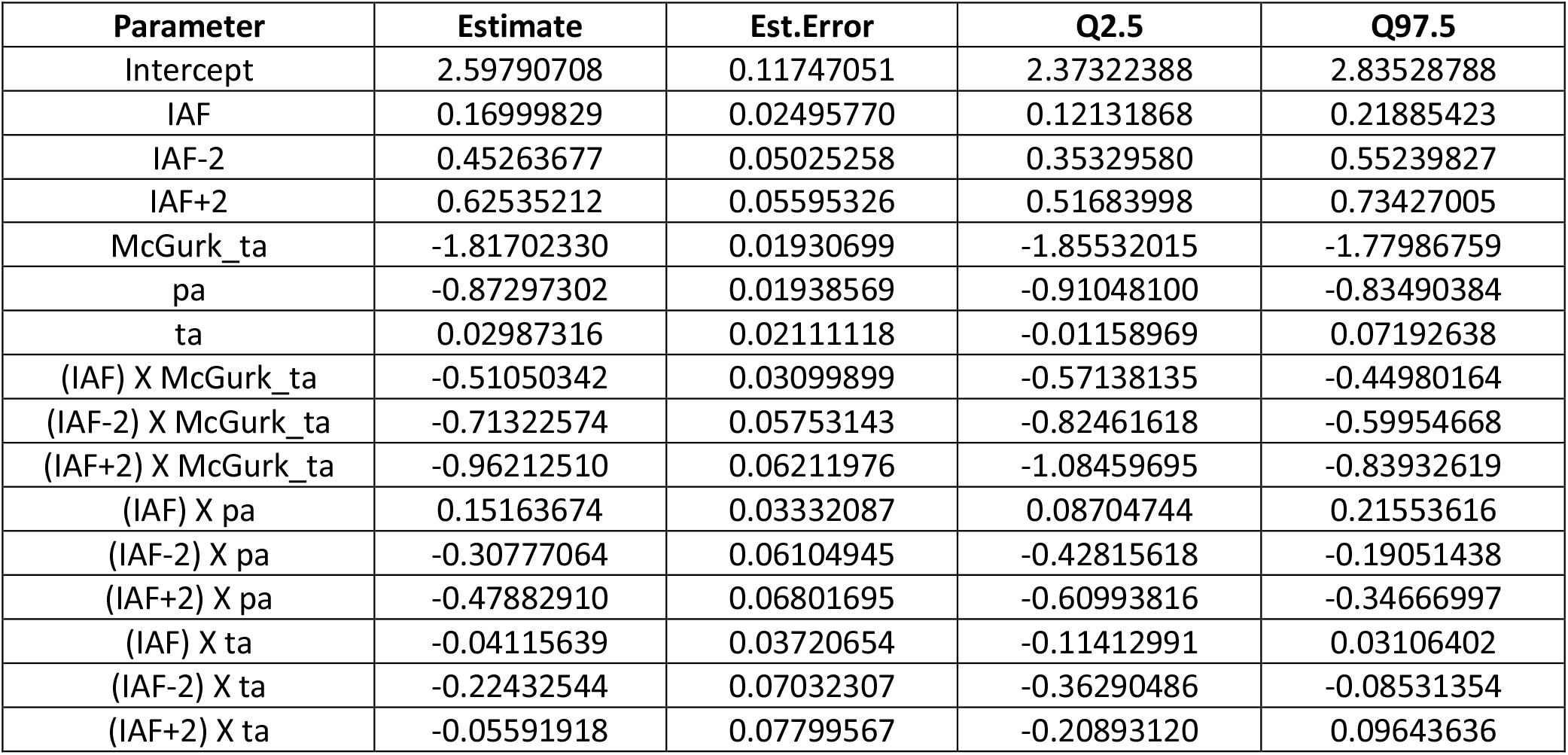
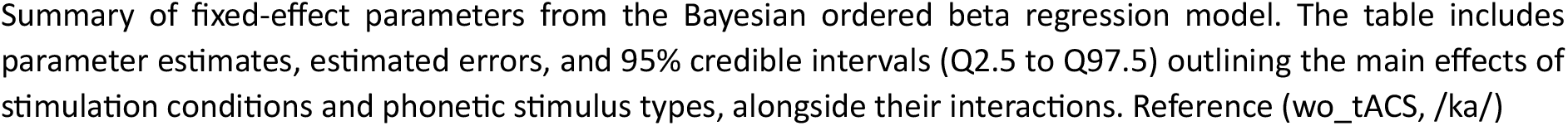
Population-level effects summary for the Generalized Linear Mixed Model (GLMM).

| Parameter | Estimate | Est.Error | Q2.5 | Q97.5 |
| --- | --- | --- | --- | --- |
| Intercept | 2.59790708 | 0.11747051 | 2.37322388 | 2.83528788 |
| IAF | 0.16999829 | 0.02495770 | 0.12131868 | 0.21885423 |
| IAF-2 | 0.45263677 | 0.05025258 | 0.35329580 | 0.55239827 |
| IAF+2 | 0.62535212 | 0.05595326 | 0.51683998 | 0.73427005 |
| McGurk_ta | -1.81702330 | 0.01930699 | -1.85532015 | -1.77986759 |
| pa | -0.87297302 | 0.01938569 | -0.91048100 | -0.83490384 |
| ta | 0.02987316 | 0.02111118 | -0.01158969 | 0.07192638 |
| (IAF) X McGurk_ta | -0.51050342 | 0.03099899 | -0.57138135 | -0.44980164 |
| (IAF-2) X McGurk_ta | -0.71322574 | 0.05753143 | -0.82461618 | -0.59954668 |
| (IAF+2) X McGurk_ta | -0.96212510 | 0.06211976 | -1.08459695 | -0.83932619 |
| (IAF) X pa | 0.15163674 | 0.03332087 | 0.08704744 | 0.21553616 |
| (IAF-2) X pa | -0.30777064 | 0.06104945 | -0.42815618 | -0.19051438 |
| (IAF+2) X pa | -0.47882910 | 0.06801695 | -0.60993816 | -0.34666997 |
| (IAF) X ta | -0.04115639 | 0.03720654 | -0.11412991 | 0.03106402 |
| (IAF-2) X ta | -0.22432544 | 0.07032307 | -0.36290486 | -0.08531354 |
| (IAF+2) X ta | -0.05591918 | 0.07799567 | -0.20893120 | 0.09643636 |

Summary of fixed-effect parameters from the Bayesian ordered beta regression model. The table includes parameter estimates, estimated errors, and 95% credible intervals (Q2.5 to Q97.5) outlining the main effects of stimulation conditions and phonetic stimulus types, alongside their interactions. Reference (wo\_tACS, /ka/)

**Table S11.** Bayes Factors for Bayesian Generalized Linear Mixed Model parameters.

| Parameter | log_BF | Effects | Labels | Raw BF10 | log<sub>10</sub>_BF10 |
| --- | --- | --- | --- | --- | --- |
| Intercept | 55.419 | fixed | High | 1.17e+24 | 24.068 |
| IAF | 12.274 | fixed | High | 2.14e+05 | 5.331 |
| IAF-2 | 17.108 | fixed | High | 2.69e+07 | 7.430 |
| IAF+2 | 21.268 | fixed | High | 1.72e+09 | 9.236 |
| McGurk_ta | 316.862 | fixed | High | 4.09e+137 | 137.611 |
| pa | 150.245 | fixed | High | 1.78e+65 | 65.251 |
| ta | -3.216 | fixed | Strong (H<sub>0</sub>) | 0.040 | -1.397 |
| (IAF) X McGurk_ta | 29.123 | fixed | High | 4.45e+12 | 12.648 |
| (IAF-2) X McGurk_ta | 25.209 | fixed | High | 8.87e+10 | 10.948 |
| (IAF+2) X McGurk_ta | 39.313 | fixed | High | 1.18e+17 | 17.073 |
| (IAF) X pa | 4.536 | fixed | Strong | 93.31 | 1.970 |
| (IAF-2) X pa | 5.229 | fixed | High | 186.61 | 2.271 |
| (IAF+2) X pa | 11.089 | fixed | High | 6.55e+04 | 4.816 |
| (IAF) X ta | -3.065 | fixed | Strong (H<sub>0</sub>) | 0.047 | -1.331 |
| (IAF-2) X ta | 1.775 | fixed | Moderate | 5.90 | 0.771 |
| (IAF+2) X ta | -2.677 | fixed | Strong (H<sub>0</sub>) | 0.069 | -1.163 |

Strength of evidence estimations for fixed-effect model parameters. The table details Log and Log10-transformed Bayes Factors (BF10) assessing the conditional main effects and interactions of stimulation type and stimulus conditions on perceptual responses. Labels denote the evidence of strength (as described in Method section 2.3.2) with (H<sub>0</sub>) marked for the values supporting null hypothesis. Reference (wo\_tACS, /ka/)

**Table S12.** Hypothesis testing for McGurk-specific contrasts.

| Comparison | Estimate | Est.Error | CI.Lower | CI.Upper | Evid. Ratio | Star | p_value | p_adj | BF10 | log10(BF10) |
| --- | --- | --- | --- | --- | --- | --- | --- | --- | --- | --- |
| (IAF) vs. wo_tACS | -0.341 | 0.018 | -0.377 | -0.306 | 8.96e-17 | * | 0.000 | 0.000 | 1.12e+16 | 16.048 |
| (IAF-2) vs. wo_tACS | -0.261 | 0.028 | -0.316 | -0.206 | 2.36e-139 | * | 0.000 | 0.000 | 4.23e+138 | 138.626 |
| (IAF+2) vs. wo_tACS | -0.337 | 0.027 | -0.390 | -0.283 | 2.00e-192 | * | 0.000 | 0.000 | 5.01e+192 | 192.700 |
| (IAF) vs. (IAF-2) | -0.080 | 0.029 | -0.137 | -0.022 | 2.77e+00 | * | 0.006 | 0.009 | 3.61e-01 | -0.442 |
| (IAF) vs. (IAF+2) | -0.004 | 0.028 | -0.059 | 0.051 | 1.03e+02 |  | 0.884 | 0.884 | 9.68e-03 | -2.014 |
| (IAF-2) vs. (IAF+2) | 0.076 | 0.036 | 0.006 | 0.147 | 8.53e+00 | * | 0.033 | 0.039 | 1.17e-01 | -0.931 |

Post-hoc hypothesis testing isolating the effect of tACS conditions on the illusory McGurk percept. The table lists parameter estimates, credible intervals, Evidence Ratios, adjusted p-values, and Bayes Factors (BF10 and Log10(BF10)) comparing baseline (wo\_tACS) to active tACS frequencies, as well as offsets directly to the IAF and within offset comparison.

**Table S13.** Interaction contrasts and credible differences across stimulus pairs.

| Condition_pair wise | Stimulus_pairwise | estimate | lower.HPD | upper.HPD | p_adj | BF10 | log10(BF10) |
| --- | --- | --- | --- | --- | --- | --- | --- |
| wo_tACS - tACS_IAF | McGurk_ta - ka | 0.510 | 0.450 | 0.571 | <0.001 | 7.62e+14 | 14.882 |
| wo_tACS - (tACS_IAF-2) | McGurk_ta - ka | 0.714 | 0.600 | 0.825 | <0.001 | Inf | > 192.7 |
| wo_tACS - (tACS_IAF+2) | McGurk_ta - ka | 0.963 | 0.837 | 1.082 | <0.001 | 2.12e+17 | 17.326 |
| tACS_IAF - (tACS_IAF-2) | McGurk_ta - ka | 0.203 | 0.088 | 0.324 | 0.001 | 9.61e+00 | 0.983 |
| tACS_IAF - (tACS_IAF+2) | McGurk_ta - ka | 0.451 | 0.325 | 0.576 | <0.001 | Inf | > 192.7 |
| (tACS_IAF-2) - (tACS_IAF+2) | McGurk_ta - ka | 0.250 | 0.098 | 0.417 | 0.003 | 5.14e+00 | 0.711 |
| wo_tACS - tACS_IAF | ka - pa | 0.152 | 0.088 | 0.216 | <0.001 | 5.58e+02 | 2.747 |
| wo_tACS - (tACS_IAF-2) | ka - pa | -0.308 | -0.430 | -0.193 | <0.001 | 6.41e+06 | 6.807 |
| wo_tACS - (tACS_IAF+2) | ka - pa | -0.478 | -0.608 | -0.346 | <0.001 | Inf | > 192.7 |
| tACS_IAF - (tACS_IAF-2) | ka - pa | -0.458 | -0.585 | -0.332 | <0.001 | Inf | > 192.7 |
| tACS_IAF - (tACS_IAF+2) | ka - pa | -0.630 | -0.768 | -0.496 | <0.001 | Inf | > 192.7 |
| (tACS_IAF-2) - (tACS_IAF+2) | ka - pa | -0.171 | -0.339 | 0.000 | 0.058 | 2.79e-01 | -0.554 |
| wo_tACS - tACS_IAF | ka - ta | -0.041 | -0.114 | 0.030 | 0.296 | 4.49e-02 | -1.348 |
| wo_tACS - (tACS_IAF-2) | ka - ta | -0.224 | -0.359 | -0.082 | 0.002 | 5.91e+00 | 0.771 |
| wo_tACS - (tACS_IAF+2) | ka - ta | -0.055 | -0.213 | 0.091 | 0.509 | 6.52e-02 | -1.186 |
| tACS_IAF - (tACS_IAF-2) | ka - ta | -0.184 | -0.333 | -0.039 | 0.020 | 5.92e-01 | -0.228 |
| tACS_IAF - (tACS_IAF+2) | ka - ta | -0.013 | -0.174 | 0.137 | 0.869 | 4.04e-02 | -1.394 |
| (tACS_IAF-2) - (tACS_IAF+2) | ka - ta | 0.169 | -0.031 | 0.364 | 0.121 | 1.81e-01 | -0.742 |
| wo_tACS - tACS_IAF | McGurk_ta - pa | 0.662 | 0.605 | 0.717 | <0.001 | Inf | > 192.7 |
| wo_tACS - (tACS_IAF-2) | McGurk_ta - pa | 0.405 | 0.317 | 0.495 | <0.001 | 1.30e+16 | 16.113 |
| wo_tACS - (tACS_IAF+2) | McGurk_ta - pa | 0.483 | 0.389 | 0.577 | <0.001 | Inf | > 192.7 |
| tACS_IAF - (tACS_IAF-2) | McGurk_ta - pa | -0.257 | -0.346 | -0.164 | <0.001 | 3.68e+10 | 10.566 |
| tACS_IAF - (tACS_IAF+2) | McGurk_ta - pa | -0.179 | -0.280 | -0.081 | 0.001 | 5.43e+00 | 0.735 |
| (tACS_IAF-2) - (tACS_IAF+2) | McGurk_ta - pa | 0.079 | -0.046 | 0.196 | 0.243 | 4.47e-02 | -1.350 |
| wo_tACS - tACS_IAF | McGurk_ta - ta | 0.470 | 0.404 | 0.533 | <0.001 | 5.73e+14 | 14.758 |
| wo_tACS - (tACS_IAF-2) | McGurk_ta - ta | 0.489 | 0.376 | 0.599 | <0.001 | 2.40e+23 | 23.380 |
| wo_tACS -<br>(tACS_IAF+2) | McGurk_ta - ta | 0.906 | 0.782 | 1.023 | <0.001 | 5.79e+15 | 15.762 |
| tACS_IAF -<br>(tACS_IAF-2) | McGurk_ta - ta | 0.019 | -0.097 | 0.139 | 0.777 | 2.11e-02 | -1.675 |
| tACS_IAF -<br>(tACS_IAF+2) | McGurk_ta - ta | 0.436 | 0.312 | 0.564 | <0.001 | 4.55e+15 | 15.658 |
| (tACS_IAF-2) -<br>(tACS_IAF+2) | McGurk_ta - ta | 0.416 | 0.256 | 0.567 | <0.001 | 1.08e+10 | 10.032 |
| wo_tACS -<br>tACS_IAF | pa - ta | -0.193 | -0.259 | -0.122 | <0.001 | 4.31e+35 | 35.634 |
| wo_tACS -<br>(tACS_IAF-2) | pa - ta | 0.082 | -0.034 | 0.207 | 0.208 | 7.08e-02 | -1.150 |
| wo_tACS -<br>(tACS_IAF+2) | pa - ta | 0.424 | 0.293 | 0.558 | <0.001 | 8.56e+18 | 18.933 |
| tACS_IAF -<br>(tACS_IAF-2) | pa - ta | 0.276 | 0.148 | 0.400 | <0.001 | 1.48e+03 | 3.169 |
| tACS_IAF -<br>(tACS_IAF+2) | pa - ta | 0.615 | 0.478 | 0.754 | <0.001 | 4.08e+42 | 42.610 |
| (tACS_IAF-2) -<br>(tACS_IAF+2) | pa - ta | 0.339 | 0.173 | 0.513 | <0.001 | 3.48e+02 | 2.541 |

Differences of differences (interaction contrasts) computed on the log-odds scale to evaluate if the tACS-related shifts in perception varied significantly by stimulus type. The table provides estimates, Highest Posterior Density (HPD) intervals, adjusted p-values, and Bayes Factors (BF10 and Log10(BF10)) demonstrating the selectivity of tACS modulation on incongruent versus congruent trials.

## Supplementary Material 4 – Correlation Analysis

From the Kuramoto oscillator model, it can be seen that the order parameter (R) value is approximately from 70% to 80% for M=10, *τ* =5ms

R is from 30% to 40% for M=20, *τ* =2.5ms

R is from 90% to 100% for M= 5, *τ* = 10*ms*

Now, our goal is to fit order parameter values to the %/ta/ responses for each of the 17 participants. For this, we shall use interpolation/extrapolation technique.1.

1. For participant 2 (P2), avg. of % /ta/response =(%/ta/ response at (IAF+2) + %/ta/ response at (IAF-2))/2=73.34% which is close to 75% Now, to get close to 75% ((70%+80%)/2) order parameter value, P2 will have M=10, *τ* =5ms The R value for P2 for this M and *τ* at (IAF+2) and (IAF-2) are listed in Table 1.
2. For P3, %/ta/ response is 80%. Hence, to get R close to 80%, we can use extrapolation to find the value of M for P3 as follows

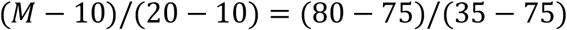

⇒M= 8.75, *τ* = 50/8.75 = 5.71*ms* The R value for P3 for this M and *τ* at (IAF+2) and (IAF-2) are listed in Table 1.
3. For P4, avg. of %/ta/ response is close to 100%. Hence, to get R close to 100%, P4 will have M=3.75, *τ* =13ms The R value for P4 for this M and *τ* at (IAF+2) and (IAF-2) are listed in Table 1.
4. For P5, avg. of %/ta/ response is close to 3%. Hence, to get R close to 3%, we can use extrapolation to find the value of M for P5 as follows

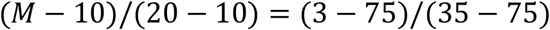

⇒M= 28, *τ* = 50/28 = 1.79*ms* The R value for P5 for this M and *τ* at (IAF+2) and (IAF-2) are listed in Table 1.
5. For P7, avg. of %/ta/ response is close to 85%. Hence, to get R close to 85%, we can use extrapolation to find the value of M for P7 as follows

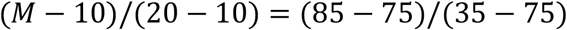

⇒M= 7.5, *τ* = 50/7.5 = 6.7*ms*. The R value for P7 for this M and *τ* at (IAF+2) and (IAF-2) are listed in Table 1.
6. For P8, avg. of %/ta/ response is 45%. Hence, to get R close to 45%, we can use interpolation to find the value of M for P8 as follows

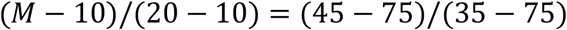

⇒M= 17.5, *τ* = 50/17.5 = 3*ms*. The R value for P8 for this M and *τ* at (IAF+2), IAF and (IAF-2) are listed in Table 1.
7. For P9, %/ta/ response is 30%. Hence, to get R close to 30%, we can use extrapolation to find the value of M for P9 as follows (*M* − 10)/(20 − 10) = (30 − 75)/(35 − 75) ⇒M= 21.25, *τ* = 50/21.25 = 2.35*ms*. The R value for P9 for this M and *τ* at (IAF+2) and (IAF-2) are listed in Table 1.
8. For P10, %/ta/ response is close to 90%. Hence, to get R close to 90%, we can use extrapolation to find the value of M for P10 as follows

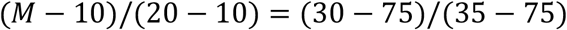

⇒M= 6.25, *τ* = 50/6.25 = 8*ms*. The R value for P10 for this M and *τ* at (IAF+2) and (IAF-2) are listed in Table 1.
9. For P12, %/ta/ response is 100%. Hence, to get R close to 100%, we can use extrapolation to find the value of M for P12 as follows

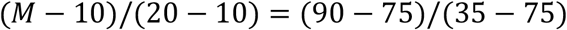

⇒M= 3.75, *τ* = 50/3.75 = 13*ms*. The R value for P12 for this M and *τ* at (IAF+2) and (IAF-2) are listed in Table 1.
10. For P13, avg. of %/ta/ response is close to 12%. Hence, to get R close to 12%, we can use extrapolation to find the value of M for P13 as follows

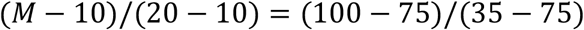

⇒M= 25.75, *τ* = 50/25.75 = 2*ms*. The R value for P13 for this M and *τ* at (IAF+2), IAF and (IAF-2) are listed in Table 1.
11. For P16, %/ta/ response is close to 55%. Hence, to get R close to 55%, we can use interpolation to find the value of M for P16 as follows

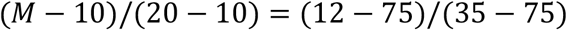

⇒M= 15, *τ* = 50/15 ≈ 3*ms*. The R value for P16 for this M and *τ* at (IAF+2) and (IAF-2) are listed in Table 1.
12. For P18, %/ta/ response is close to 95%. Hence, to get R close to 95%, P18 will have M=5, *τ* =10ms. The R value for P18 for this M and *τ* at (IAF+2) and (IAF-2) are listed in Table 1.
13. For P19, avg. of %/ta/ response is close to 85%. Hence, to get R close to 85%, P19 will have M= 7.5, *τ* =6.7ms. The R value for P19 for this M and *τ* at (IAF+2) and (IAF-2) are listed in Table 1.
14. For P20, avg. of %/ta/ response is 15%. Hence, to get R close to 15%, we can use extrapolation to find the value of M for P20 as follows

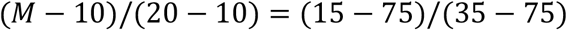

⇒M= 25, *τ* = 50/25 = 2*ms*. The R value for P20 for this M and *τ* at (IAF+2) and (IAF-2) are listed in Table 1.
15. For P22, %/ta/ response is close to 95%. Hence, to get R close to 95%, P22 will have M=5, *τ* =10ms. The R value for P22 for this M and *τ* at (IAF+2) and (IAF-2) are listed in Table 1.
16. For P23, %/ta/ response is 100%. Hence, to get R close to 100%, P23 will have M=3.75, *τ* =13ms. The R value for P23 for this M and *τ* at (IAF+2), IAF and (IAF-2) are listed in Table 1.
17. For P26, %/ta/ response is 100%. Hence, to get R close to 100%, P26 will have M=3.75, *τ* =13ms. The R value for P26 for this M and *τ* at (IAF+2), IAF and (IAF-2) are listed in Table 1.

Although R values are included in Table 3 for 10 participants at M= 10, *τ* = 5*ms*, we have also checked R values for these 10 participants at other M, *τ* and *β* values. In all these cases, values of order parameter at (IAF-2) and (IAF+2) are close to 1, whereas values of order parameter at IAF are close to zero. From Table 5, we can conclude that model-predicted order-parameter asymmetry (R values at IAF-2 minus IAF+2) vs. behavioural asymmetry (the proportion of McGurk /ta/ responses at IAF−2 minus IAF+2) across 27 participants shows a statistically significant, positive correlation. Spearman’s correlation is greater than Pearson’s, indicating that the relationship between the model-predicted order-parameter asymmetry and behavioural asymmetry is monotonic (consistently increasing or decreasing) but non-linear. The Fisher z-transformation is used to calculate a confidence interval (CI) for the Pearson correlation, whereas the bootstrapping method is used for the Spearman correlation. In Table 3, participant 21 is marked red because it shows a discrepancy: the order parameter value at (IAF+2) is greater than at (IAF-2) for M= 10, *τ* = 5*ms* and *β* = 1 after invoking the effective coupling strategy.

We have not included the without-tACS effect in the correlation analysis because if we put *I* = 0 in eq. (S18), we get,

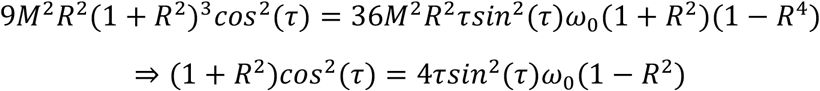

For small *τ, sin*(*τ*) ≈ *τ* and *cos*(*τ*) ≈ 1

Therefore, we can write,

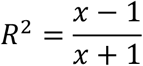

where *x* = 4*τ*^3^*ω*_0_

For *R* to be real, *R*^2^ ≥ 0 which implies *x* ≥ 1

At *τ* = 5*ms*, we get, *ω*_0_ ≥ 2 × 10^6^ which implies *f* ≥ 318309.89 *Hz* which is not in the desired frequency range. Similarly, if we replace M with *M*_*eff*_, the same scenario occurs. Hence, no real solution of *R* exists without the tACS case.

## Notes

### Competing Interest Statement

The authors have declared no competing interest.

